# MRI-based Parcellation and Morphometry of the Individual Rhesus Monkey Brain: a translational system referencing a standardized ontology

**DOI:** 10.1101/699710

**Authors:** R. Jarrett Rushmore, Sylvain Bouix, Marek Kubicki, Yogesh Rathi, Edward H. Yeterian, Nikos Makris

**Author notes:** Contributions are equal (Edward H. Yeterian, Nikos Makris). Corresponding Author: Nikos Makris, M.D., Ph.D., Massachusetts General Hospital, 149 Thirteenth Street, Charlestown, MA 02129 USA.

## Abstract

The rhesus macaque is the closest animal model to the human, and investigations into the brain of the rhesus monkey has shed light on the function and organization of the primate brain at a scale and resolution not yet possible in studies of the human brain A cornerstone of the linkage between non-human primate and human studies of the brain is magnetic resonance imaging, which allows for an association to be made between the detailed structural and physiological analysis of the non-human primate and that of the human brain. To further this end, we present a novel parcellation system for the rhesus monkey brain, referred to as the monkey Harvard Oxford Atlas (mHOA) which is based on the human Harvard-Oxford Atlas (HOA) and grounded in an ontological and taxonomic framework. Consistent anatomical features were used to delimit and parcellate brain regions in the macaque, which were then categorized according to functional systems. This system of parcellation will be expanded with advances in technology and like the HOA, will provide a framework upon which the results from other experimental studies (e.g., functional magnetic resonance imaging (fMRI), physiology, connection, graph theory) can be interpreted.

## INTRODUCTION

A fundamental goal of current basic and clinical neuroscience is to relate function to structure (e.g., Mesulam, 2000; Swanson, 2012). The validity of these function-structure relationships is closely associated with the accuracy of anatomical information (Mesulam, 2000; Swanson, 2012; Cieslik et al., 2013). At a cortical level, a key aspect of structure is represented by the cytoarchitecture of the different cortical areas, each of which have specific patterns of connectivity. In the non-human primate, various mesoscale-level descriptions of the cytoarchitecture and connections have been achieved (Bohland et al., 2009b), which has enabled the definition of a brain circuit diagram (BCD). The concept of a BCD interrelates gray matter cortical and subcortical brain regions with connecting axonal pathways. In this concept is also encapsulated the definition of a fiber tract, which necessarily includes origins and terminations in the gray matter in addition to a connecting white matter stem (Makris et al., 1997) (Fig 1A). As a result, the complete definition of a BCD necessitates these two types of information, namely, subcortical and cortical gray matter maps and white matter fiber tracts. This mesoscale level (e.g., Swanson, 2012; Bohland et al., 2009b) of structural neuroanatomical understanding has not been achieved to date in humans, where research has focused primarily on stems of fiber pathways and is methodologically unable to resolve their origins and terminations (Bohland et al., 2009b; Maier-Hein et al., 2017; Schilling et al., 2019) (Fig 2A, B). Until appropriate technological advancements are made, a comparative approach based on structural homologies can be used to extrapolate precise neuroanatomical data about origins and terminations from the monkey to enhance our understanding of the human BCD, and thereby provide translational relevance in basic and clinical neuroscience (Schmahmann & Pandya, 2005) (Fig 3). The insights from these homologies are needed because one can invasively experiment on monkeys and mimic to a large degree changes in human behavior, or even investigate biological consequences of structurally severing certain brain circuits.

**Figure 1.**
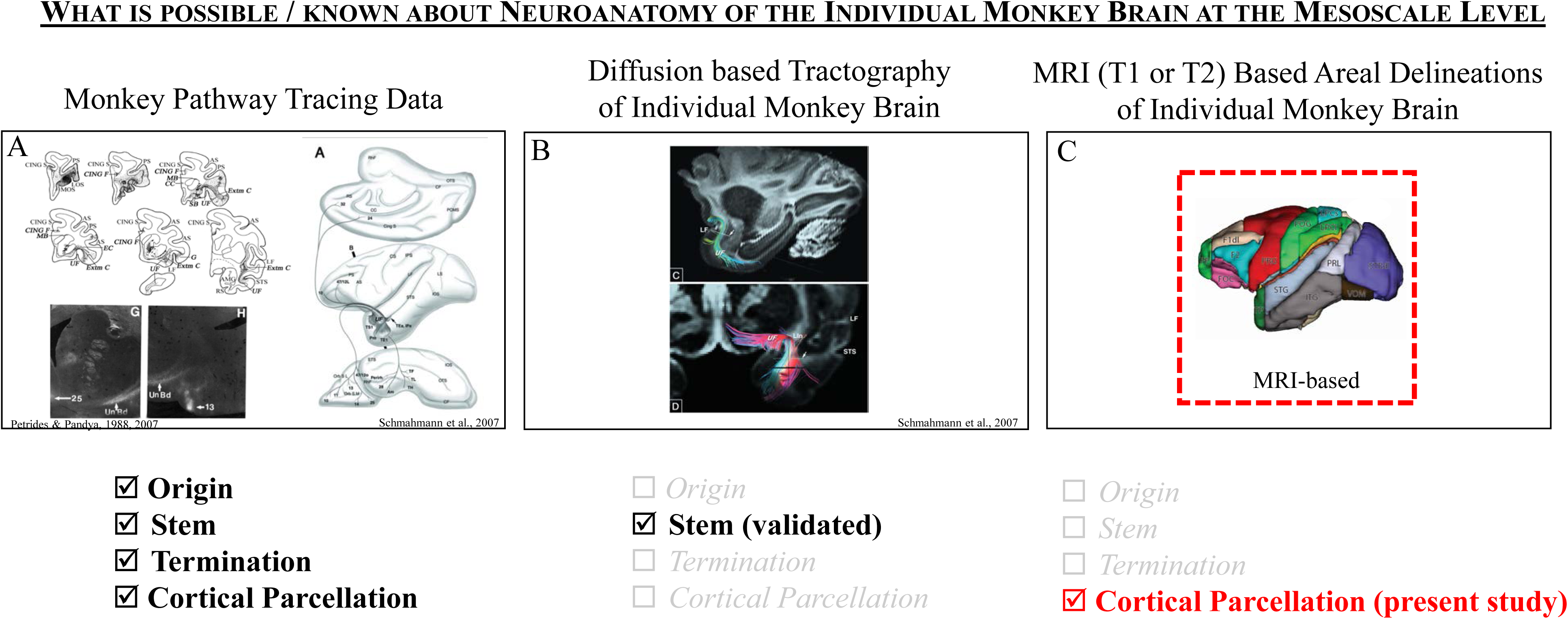
The state of neuroanatomical knowledge in the individual monkey brain. A. Tract tracing data have been accumulated to reveal the connections of most areas of the monkey brain. These invasive methodologies reveal the origin, stem and termination of axonal bundles, and also illuminate divisions in cortical and subcortical structure by virtue of the use of histological stains applied to adjacent or the same brain sections as those for tract tracing. Importantly, these methods do not apply to the in vivo brain. B. Non-invasive diffusion MRI tractography is a method to infer the presence of fiber tracts based on the diffusion of water molecules within white matter. These methods do not have the capability of revealing origins and terminations, but are capable of illustrating stems of fiber pathways. C. MRI-based parcellations divide brain regions, but, by themselves, do not specify origins, terminations or stems of pathways. The three methods together are needed to define the complete brain circuit diagram of the individual monkey *in vivo*. Upper left figure in A is copyright 2007 Society for Neuroscience and is used with permission; lower left figure in A is used with permission from Petrides & Pandya (1988). Right hand figure in A, and figure B is used with permission from Schmahmann et al., 2007.

**Figure 2.**
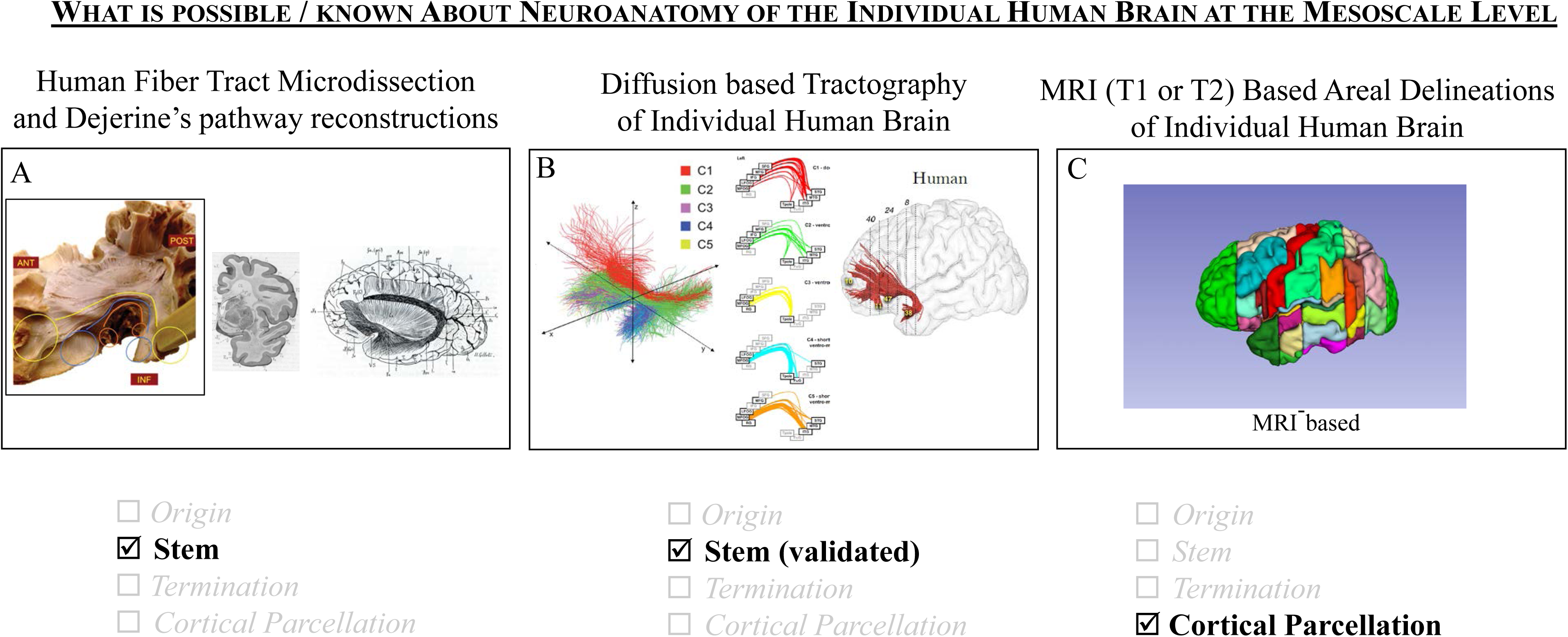
The state of neuroanatomical knowledge in the individual human brain. A. Since human brains in vivo are not amenable to the use of invasive tract tracing techniques, direct knowledge of pathways is derived from post-mortem microdissection, and from classical histological studies. These methods do not visualize origins or terminations and are validated only for pathway stems (see previous paper). B. Diffusion MRI tractography methods allow for the inference of pathways, but like the same techniques in the monkey (see Figure 1A), provide validated data only for stems. C. Cortical parcellation schemes, such as that of the Harvard Oxford Atlas, are specified in the individual brain based on consistent anatomical landmarks. These schemes can form the foundation for connectional data derived from non-invasive methods, but do not provide validation of terminations and origins of pathways. Unlike the monkey, combining these three methods in the human are insufficient to define the complete human brain circuit diagram. Left hand figure in A, and figures in B used with permission from Hau et al., 2012. Central and right-hand figures in A from Dejerine and Dejerine-Klumpe (1895).

**Figure 3.**
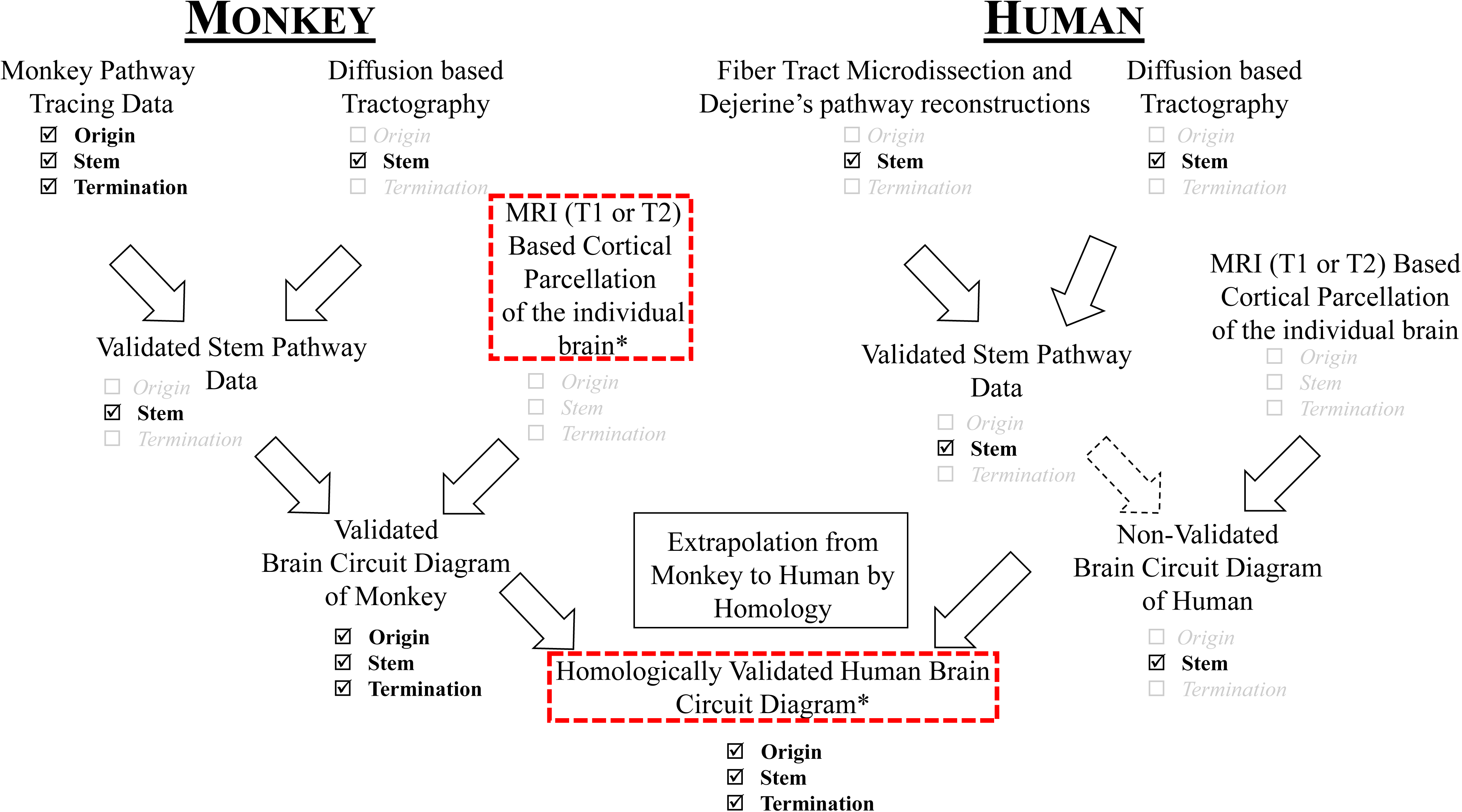
The combination and sequence of approaches in the monkey (Figure 1) that will validate diffusion MRI tractographic technology in the context of cortical parcellation and will complete the non-human primate brain circuit diagram in vivo (left). The same approach in the human (right) will lead to an incomplete brain circuit diagram because of the inability to visualize terminations or origins of pathways. However, the validated and confirmed monkey brain circuit diagram will be able to produce a more complete human brain circuit diagram through homological comparisons.

Current advances in Magnetic Resonance Imaging (MRI) technology have enabled the study of brain anatomy in a non-invasive way and *in vivo*, an approach that has yielded tremendous clinical benefit. More specifically, neuroimaging techniques such as T1- and T2-weighted MRI have provided information about gray matter structure, and diffusion imaging (dMRI) has been used to infer the location of white matter tracts in monkeys and humans. It should be clarified that the level of detail in gray and white matter that is possible with histological techniques at the mesoscale level has not been obtained using MRI in either monkeys or humans. Regarding gray matter, the closest level of correspondence between human cytoarchitectonic cortical maps and MRI-based parcellation maps is guided by identifiable morphological landmarks present in the individual brain such as sulci and gyri or structural correlations with T1 and T2 biophysical parameters (Caviness et al., 1996; Rademacher et al., 1992; Fischl et al., 2004; Glasser & Van Essen, 2011). In the monkey, an approach to cortical parcellation that takes into consideration individual variation in cortical gray matter structure has not been achieved to date. This precision self-referential approach based on individual neuroanatomical features (Kennedy et al., 1997, 1998; Makris et al., 2002), as opposed to an atlas-based approach, is necessary for two reasons. First, it will define more precisely monkey BCDs using MRI technology. Second, this will improve validation of human BCDs through a homological approach, given that the validation of human fiber tracts is limited to stems (Fig 2B). This logic is illustrated in Fig 3. An additional necessary objective is to integrate this logic into an established computational neuroanatomy system in the field of neuroimaging.

The purpose of this manuscript is to describe a system of parcellating the macaque cerebral cortex using conventional neuroimaging and tools, which follows the same type of theoretical, neuroanatomical, and ontological reasoning as in the Harvard-Oxford Atlas (HOA) of the human brain (Caviness et al., 1996; Makris et al., 1999; 2004, 2005). More precisely, we use the human HOA paradigm and apply it to analysis of the monkey brain. We provide an anatomical framework that can be related to cytoarchitectonic fields as well as functional regions and systems, specifically primary, unimodal association, and heteromodal association. This framework is based on the specific morphology of individual brains through the use of sulci, other visible landmarks, and anatomical convention. Unlike procedures based on fitting individual brains to an atlas or template that removes interindividual variability in brain structure to identify commonalities in a common space, the HOA system preserves variability in brain structure by creating a self-referential system for each brain. As proof of concept, we applied this system in one young adult macaque monkey brain MRI dataset, which constitutes a monkey equivalent of the human HOA (mHOA). Furthermore, it presents a translational system for integrating information from monkey to human using structural imaging. At the same time, it incorporates a uniform ontological framework similar to that described by the HOA system and compatible with such ontological approaches as NeuroNames (Martin & Bowden, 1996), the Federative Internationale Programme for Anatomical Terminology (FIPAT) (e.g., ten Donkelaar et al., 2017, 2018), and Swanson’s Neuroanatomical Terminology (Swanson, 2015). This ontological approach makes the structural delineation of individual brains more systematic and organized in its lexicon and terminologies, which is particularly important in comparative neuroanatomy when extrapolating from one species to another. It will also provide a basis for addressing similarities in structure and function in an evolutionary context and also for interrelating and organizing current data basing and data mining approaches. We anticipate that this parcellation scheme will be applied in basic and clinical systems neuroscience and will have translational value in validating the human BCD. The systematic and individualized system for parcellating cortical and subcortical gray matter, the mHOA, is a novel and essential first step for more precise formulations of primate BCDs.

## METHODS

All procedures involving animals were approved by the IACUC at Boston University School of Medicine and at Massachusetts General Hospital. A 6.7-year-old female rhesus monkey was anesthetized with ketamine and xylazine (20mg/kg;0.2-0.4mg/kg) and placed in an MRI-compatible stereotaxic head holder as previously described (Makris et al., 2010). The monkey was scanned using a 1.5 T Siemens Sonata magnet at the MGH-NMR Center at the Charlestown Navy Yard. The MP-RAGE volumes were acquired with 0.8×0.8mm2 in plane resolution with 1.0mm thick slices with the following parameters: TR=2.73ms, TE=2.8ms (min), TI=300ms, flip angle =degrees, matrix = 256×256, bandwidth=190Hz/pixel, NEX=4. Approximately 128 slices were acquired with zero gap, increasing slice thickness to cover the brain.

After image acquisition, general segmentation procedures were used to separate the cortex from white matter and subcortical areas (Figure 4). The cortex was initially segmented as a single region of interest and labeled as cerebral cortex. The following subcortical regions were segmented: caudate, putamen, nucleus accumbens, globus pallidus, thalamus, amygdala, basal forebrain, hippocampus, brainstem, ventral diencephalon and cerebellum. General segmentation and subdivision of cerebral cortex was carried out as in previous approaches to the HOA with the Cardviews program (Filipek et al., 1994; Caviness et al., 1996; Makris et al., 1999), and 3D Slicer. 3D Slicer was used to construct 3D volumes and to calculate parcellation unit volumes.

**Figure 4:**
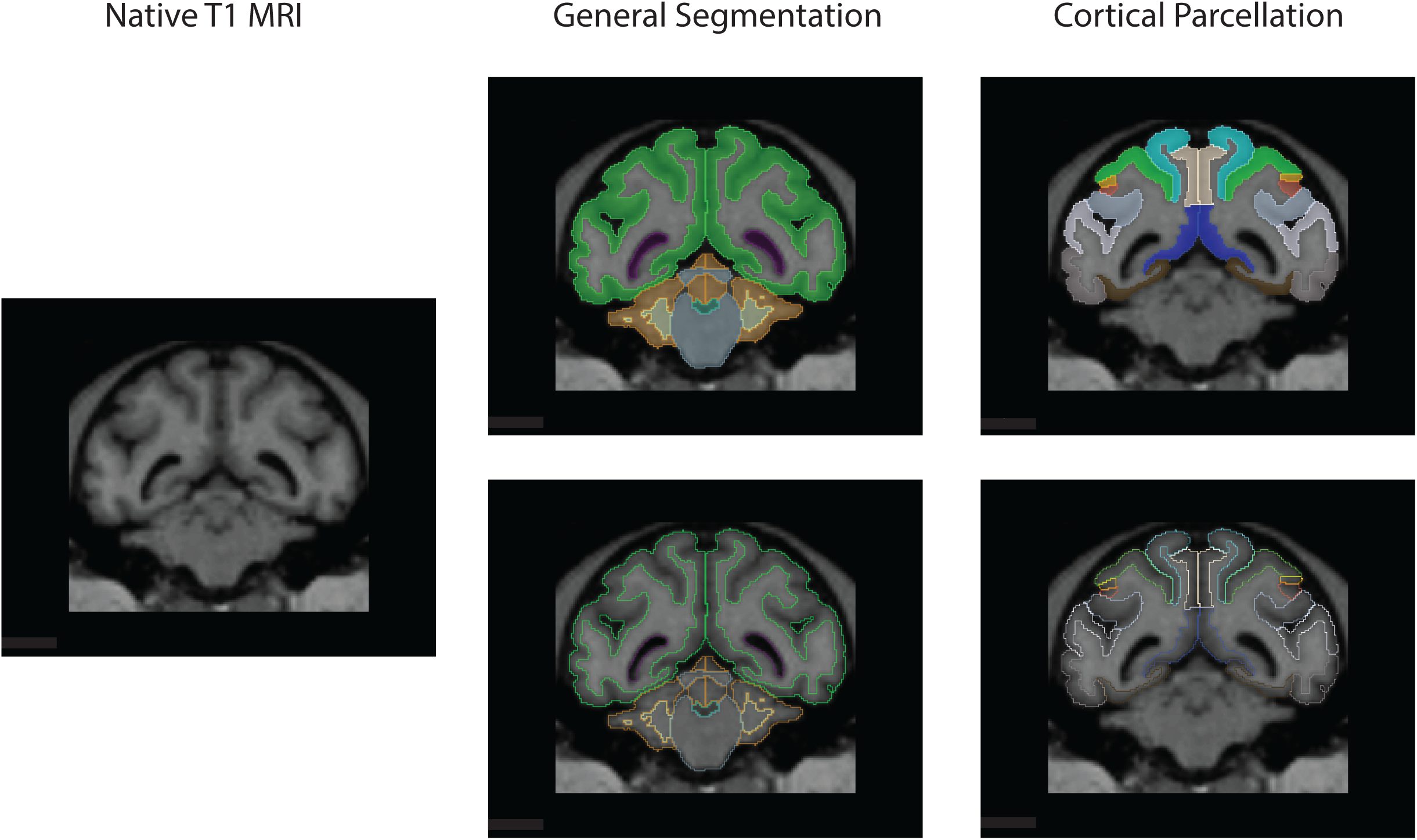
System of Parcellation. An individual T1 image set (left) was acquired and a cortical ribbon and subcortical structures were divided (middle) through a general segmentation process (green – cortical ribbon, orange, cerebellar cortex, yellow – cerebellar white matter, blue-fourth ventricle, blue-grey – brainstem). The cortical ribbon was further divided into regions according to the system described in this paper (right): dark blue – CGP; dark gray – ITG; brown – VMO; gray – STG; light blue – LPCs; light brown – MPC; light green – LPCi; white-PRL.

## RESULTS

Herein, we accomplished three main objectives. First, we developed a parcellation framework of the cerebral cortex of the rhesus monkey based on cortical morphological features in a fashion that closely reflects its underlying structure. Second, we made this system available for neuroimaging use by parcellating one exemplar rhesus monkey brain using T1 MRI. Third, we have referenced our system to standard ontological systems used for the macaque and human brain (i.e., Neuronames (Bowden & Martin, 1995; Bowden & Dubach, 2003; Bowden et al., 2012; Swanson, 2015; FIPAT cf TNA p58, items 1988-2199)). Finally, the system was developed to generate an analogous framework for rhesus sulci/gyri/regions as in the human brain, which we propose here as the macaque Harvard Oxford Atlas (mHOA).

We adopted the use of a systematic and systems-based parcellation of the cerebral cortex as has been used by our group for the human HOA (e.g., Rademacher et al., 1992; Caviness et al., 1996; Makris et al., 2010). This system is based on the division of the cortical surface by discrete anatomical conventions using consistent sulcal landmarks in conjunction with a series of coronal planes that are defined by common surface neuroanatomical landmarks. By adopting such an approach, individual brains can be parcellated according to their specific anatomical features using neuroimaging-based morphometric analysis.

### Sulcal anatomy of the lateral cerebral surface

The divisions of the lateral, mid-sagittal, ventral, and intrasylvian surfaces are shown in Figure 4 and listed in Table 1.

**Table 1:**
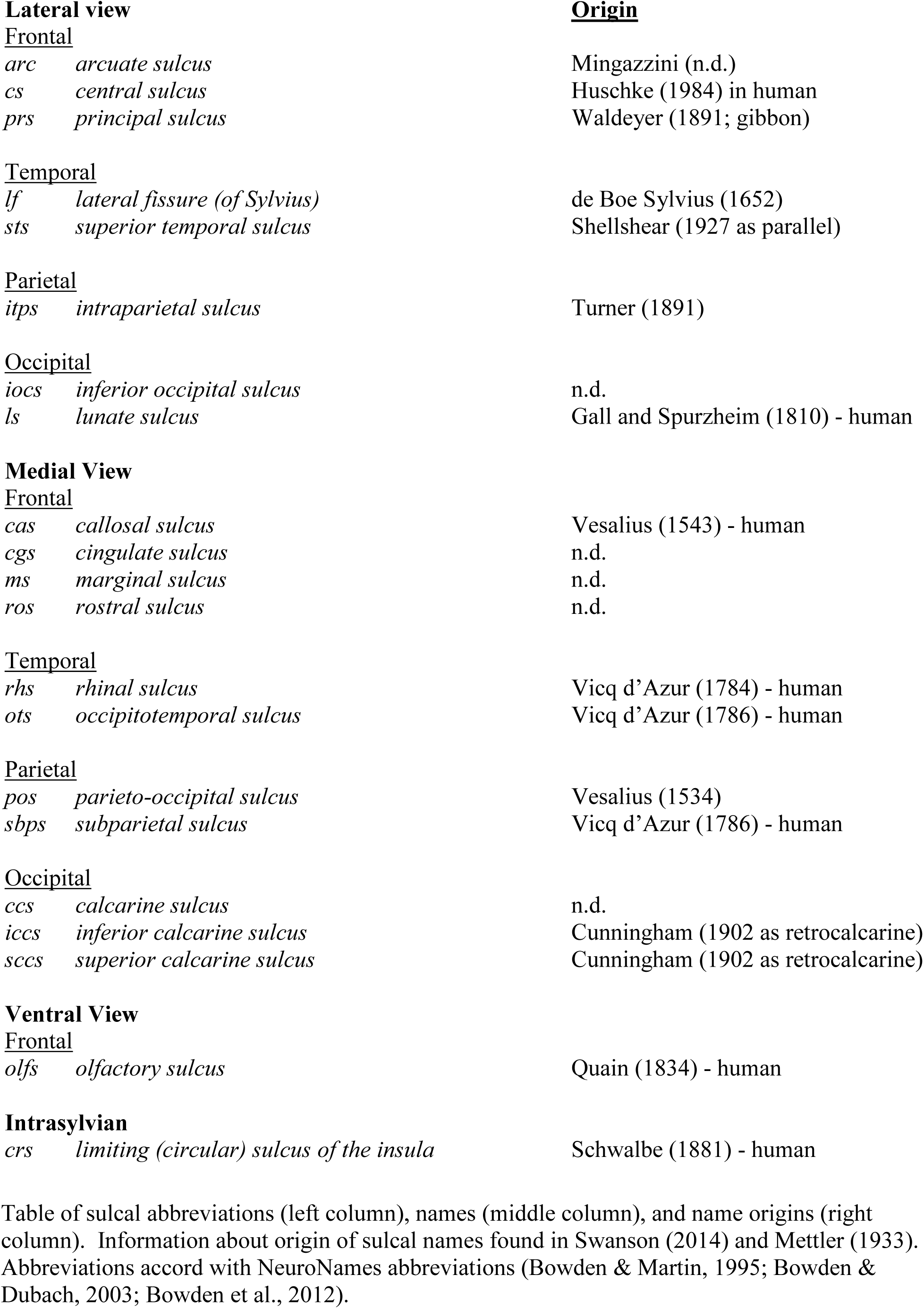
Sulcal Abbreviations

#### 1. Frontal Lobe

*Central sulcus* (NeuroNames (NN) ID 48). The central sulcus is the posterior border of the frontal lobe. It extends from the dorsal aspect of the interhemispheric fissure ventrolaterally on the lateral cerebral surface to end above the lateral fissure.

*Arcuate sulcus* (NN ID 2379). The arcuate sulcus is located rostral to the central sulcus. It is shaped like the number 7 such that the concavity of the sulcus faces rostrally. Its superior limb (ramus) is oriented roughly in the horizontal plane and its inferior limb (ramus) arcs inferiorly and rostrally, but does not reach the hemispheric margin at its lateral inferior convexity. A small sulcus referred to as posterior ramus or the spur of the arcuate extends for a short distance into the precentral gyrus from the caudal apogee of the arcuate sulcus.

*Principal sulcus* (NN ID 66). This sulcus is in the horizontal plane and extends towards the frontal pole from the concavity of the arcuate sulcus. It does not connect to the arcuate sulcus nor does it reach the frontal pole.

#### 2. Temporal Lobe

*Lateral fissure* (also referred to as the Sylvian fissure; NN ID 49). This is an extremely deep sulcus in the depths of which is found the insula, a region of cortex encircled and buried by the banks of the lateral fissure. The antero-ventral half of the fissure separates the frontal lobe from the temporal lobe, and the more posterior-dorsal half separates the parietal from the temporal lobe. The superior lip of the sulcus is referred to as the frontal operculum within the frontal lobe and as the parietal operculum in the parietal lobe. The ventral lip of the sulcus is referred to as the supratemporal plane throughout its course.

*Superior temporal sulcus* (NN ID 129). This sulcus roughly parallels the lateral fissure ventrally and is the major sulcus of the temporal lobe on the lateral surface. It extends from the temporal polar region into the parietal cortex, where it ends close to or at the union of the intraparietal sulcus and the lunate sulcus.

#### 3. Parietal Lobe

*Intraparietal sulcus* (NN ID 97). This deep sulcus is located caudal to the central sulcus. It progresses caudally and dorsally above the lateral fissure to end at a union with the lunate sulcus and the superior temporal sulcus at the dorsal hemispheric margin.

#### 4. Occipital lobe

*Lunate sulcus* (NN ID 150). This sulcus begins at a location caudal to the superior third of the STS and travels dorsally, roughly in the coronal plane, until it intersects with the intraparietal sulcus and the superior temporal sulcus at the dorsal hemispheric margin. From this union, a merged sulcus travels towards the midline and gives rise to the parieto-occipital sulcus.

*Inferior occipital sulcus* (NN ID 144). This sulcus begins below the inferior tip of the lunate sulcus and extends caudally, following a quasi-horizontal trajectory to the hemispheric margin

### Sulcal Anatomy of the Medial Cerebral Surface

*Callosal sulcus* (NN ID 36). This sulcus occupies the interface between the corpus callosum and the gray matter of the medial cerebral surface. It follows the contour of the corpus callosum.

*Rostral sulcus* (NN ID 76). The rostral sulcus is found beneath the rostrum of the corpus callosum, at the inferior aspect of the medial frontal lobe. It extends from underneath the callosum to the rostral tip of the hemisphere in a roughly horizontal plane.

*Cingulate sulcus* (NN ID 43). The cingulate sulcus in the rhesus macaque begins superior to the rostral tip of the rostral sulcus, and follows the contour of both the hemispheric margin and the corpus callosum as it travels caudally to the parietal lobe. Above the caudal portion of the body of the corpus callosum, it rises superiorly to become the marginal sulcus (marginal ramus of the cingulate sulcus).

*Subparietal sulcus* (NN ID 102). As the cingulate sulcus rises to become the marginal ramus, a sulcus similar in position to the main cingulate sulcus also becomes evident and is called the subparietal sulcus. It extends caudally along the course of the callosum, but often is disconnected from the cingulate sulcus.

*Parieto-occipital sulcus* (NN ID 52). This deep sulcus begins on the dorsal aspect of the cerebrum and extends to the medial cerebral aspect, where it separates the parietal from the occipital lobe.

*Calcarine sulcus* (NN ID 44). In the macaque, this deep sulcus extends caudally from below the splenium of the corpus callosum under the parieto-occipital sulcus. Near the occipital pole, it branches into a superior calcarine sulcus (also known as the dorsal ramus) and an inferior calcarine sulcus (ventral ramus).

### Sulcal anatomy of the ventral surface of the cerebrum

*Rhinal sulcus* (NN ID 41). The rhinal sulcus is on the ventromedial surface of the temporal lobe and courses caudally from the temporal pole. The caudal extent of the rhinal sulcus often is located medial to the rostral tip of the occipitotemporal sulcus.

*Occipitotemporal sulcus* (NN ID 55). This sulcus courses along the ventral surface of the temporal lobe into the occipital lobe turning ventromedially towards the calcarine sulcus.

*Olfactory sulcus* (NN ID 78). This sulcus runs rostro-caudally parallel to the interhemispheric fissure on the ventromedial surface of the frontal lobe.

### Sulcal anatomy within the lateral fissure

*Limiting sulcus of the insula* (NN ID 51). This sulcus, also known as the circular sulcus of the insula, circumscribes the insula and is present throughout almost the entire extent of the insula, with its superior portion forming the superior border of the insula, and its inferior portion forming the insula’s lower border.

### Limiting Planes

As in previous approaches, the cortex was separated into parcellation units based in part on boundaries derived from the location of the limiting sulci detailed above (Rademacher et al., 1992). These boundaries were supplemented and completed by a set of twelve coronal planes defined on the basis of anatomical landmarks (Figure 5; Table 2). By convention, these planes are referred to as coronal planes A-L. Seven coronal planes extend over more than one surface of the brain (e.g., medial, lateral or ventral; coronal planes A, B, E, G, I, J, L). In addition, there are several non-coronal planes (three along the horizontal axis, and two along oblique orientations) that form discrete borders and that are depicted in lower case (non-coronal planes a-e). Oblique planes (a, c) are positioned by first determining the endpoints of the non-coronal limiting plane with respect to limiting planes or sulci, then defining this plane progressively through the coronal planes to connect these endpoints (Figure 5). The interplay between the different cardinal planes, with the aid of drawing tools, allows the demarcation of these oblique planes (e.g., Rademacher et al., 1992). It should be noted that as in Rachemacher, the planes were designated based on rostral-caudal position (i.e., plane A is the most rostral, B is the second most rostral., etc.).

**Figure 5.**
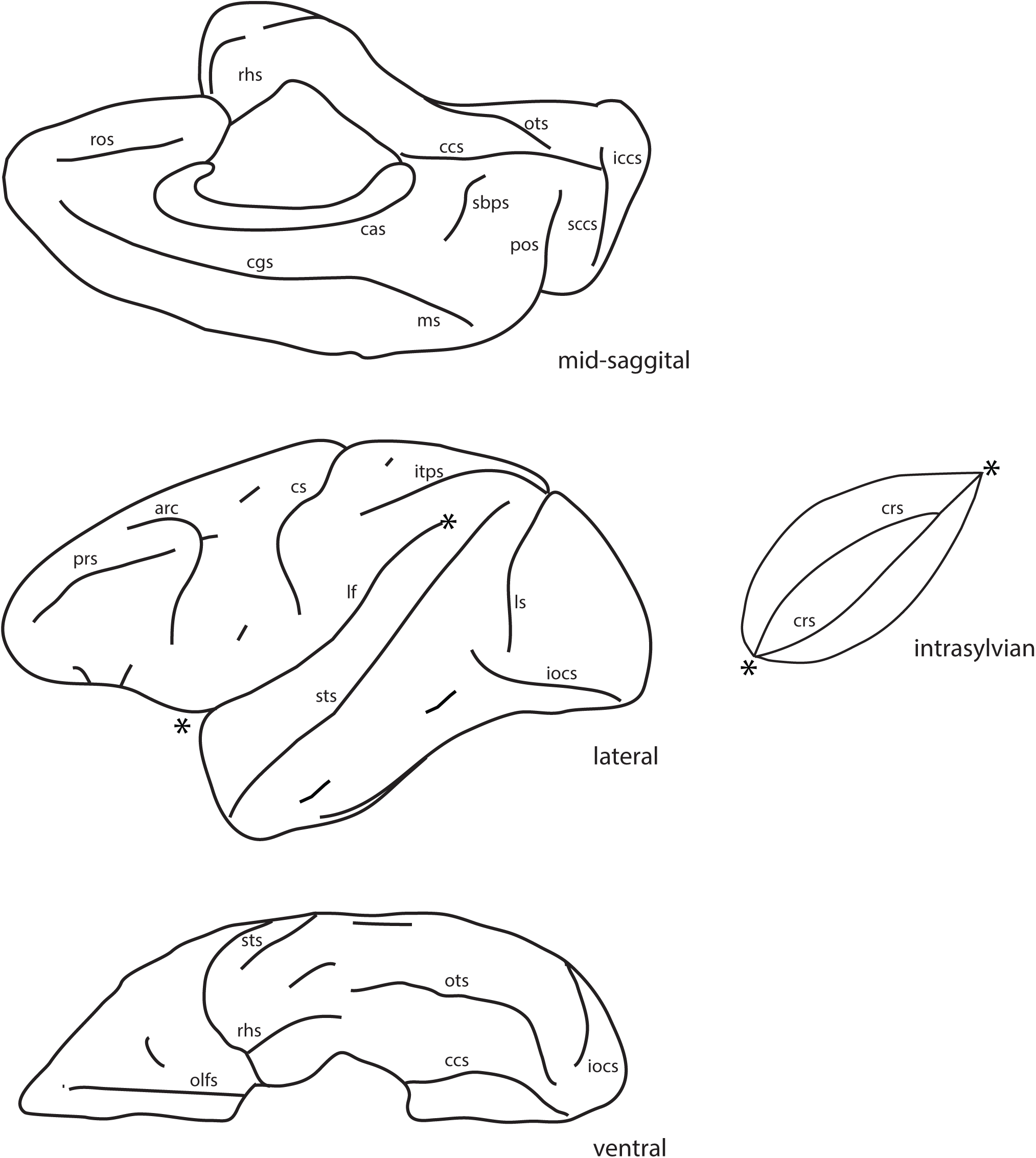
The sulci of the macaque monkey brain from the mid-sagittal (upper), lateral (middle), and ventral (lower) views. The cerebral cortex within the lateral fissure (between the asterisks) is opened to show the constituent regions (to the right of the lateral view). Abbreviations: arc: arcuate sulcus, cas: callosal sulcus, ccs: calcarine sulcus, cgs: cingulate sulcus,, iccs: inferior calcarine sulcus, iocs: inferior occipital sulcus, itps: intraparietal sulcus, crs: limiting sulcus of the insula, lf: lateral fissure, ls: lunate sulcus, ms: marginal sulcus, olfs: olfactory sulcus, ots: occipitotemporal sulcus, pos: parieto-occipital sulcus, prs: principal sulcus, ros: rostral sulcus, rhs: rhinal sulcus, sccs: superior calcarine sulcus, sbps: subparietal sulcus, sts: superior temporal sulcus

**Table 2:**
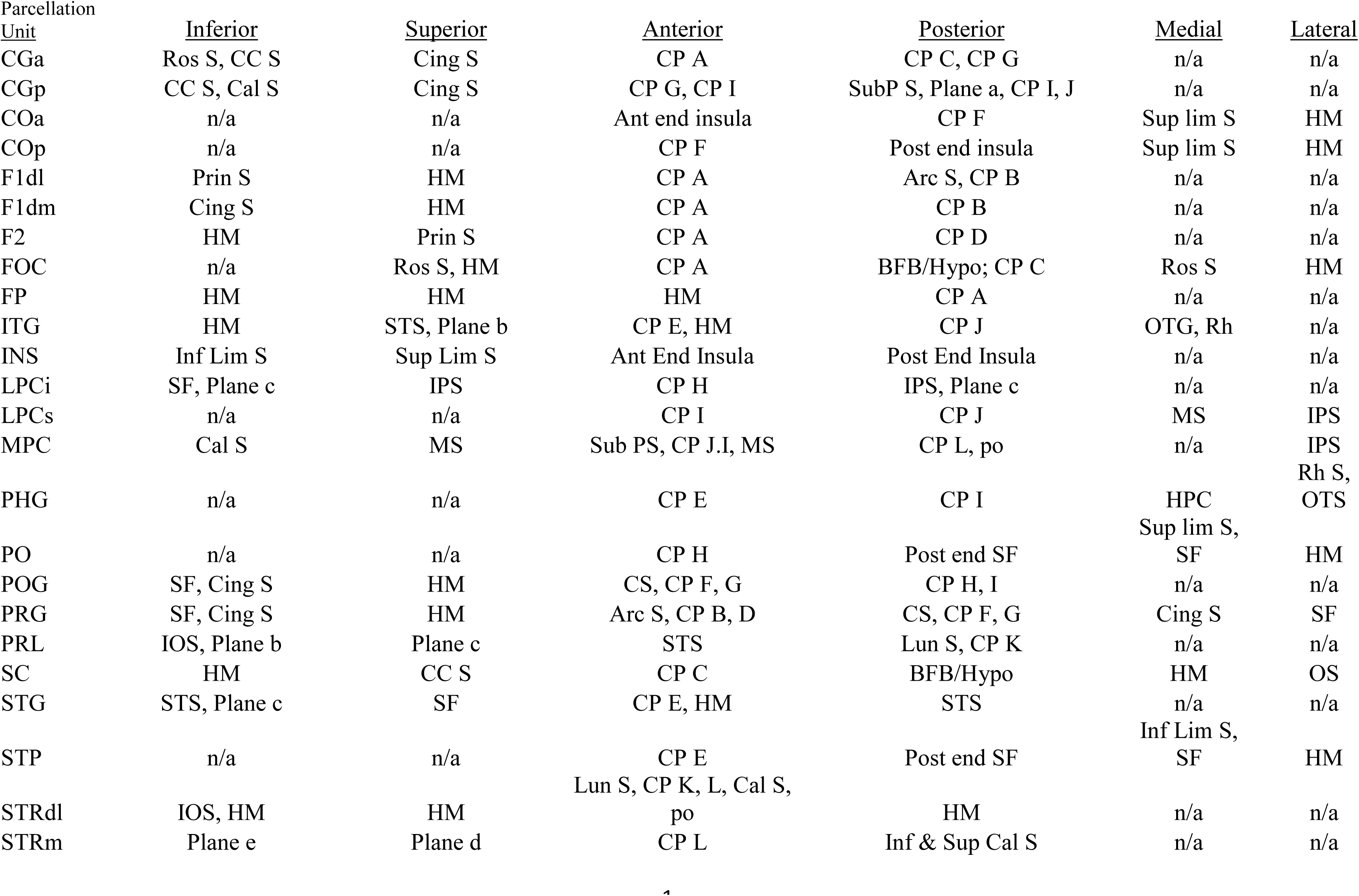
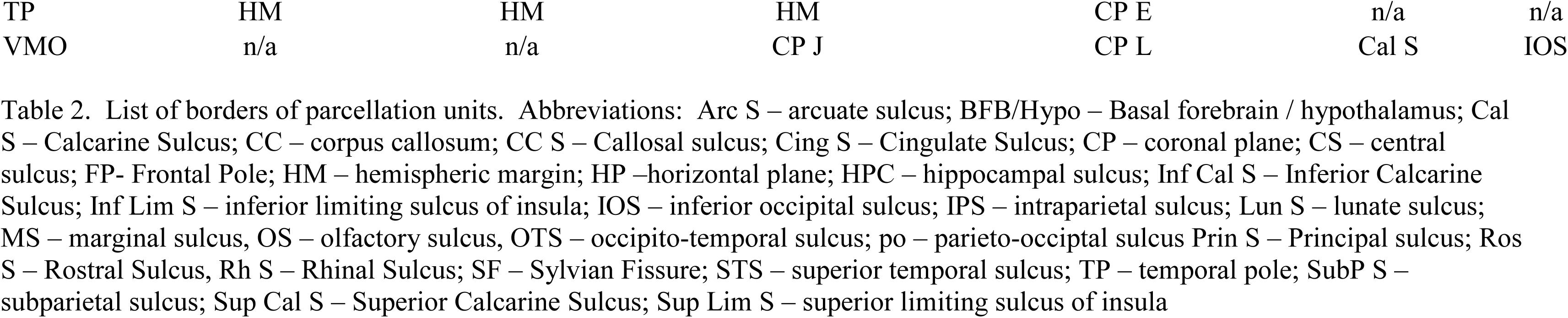
Parcellation unit boundaries

**Table 3:**
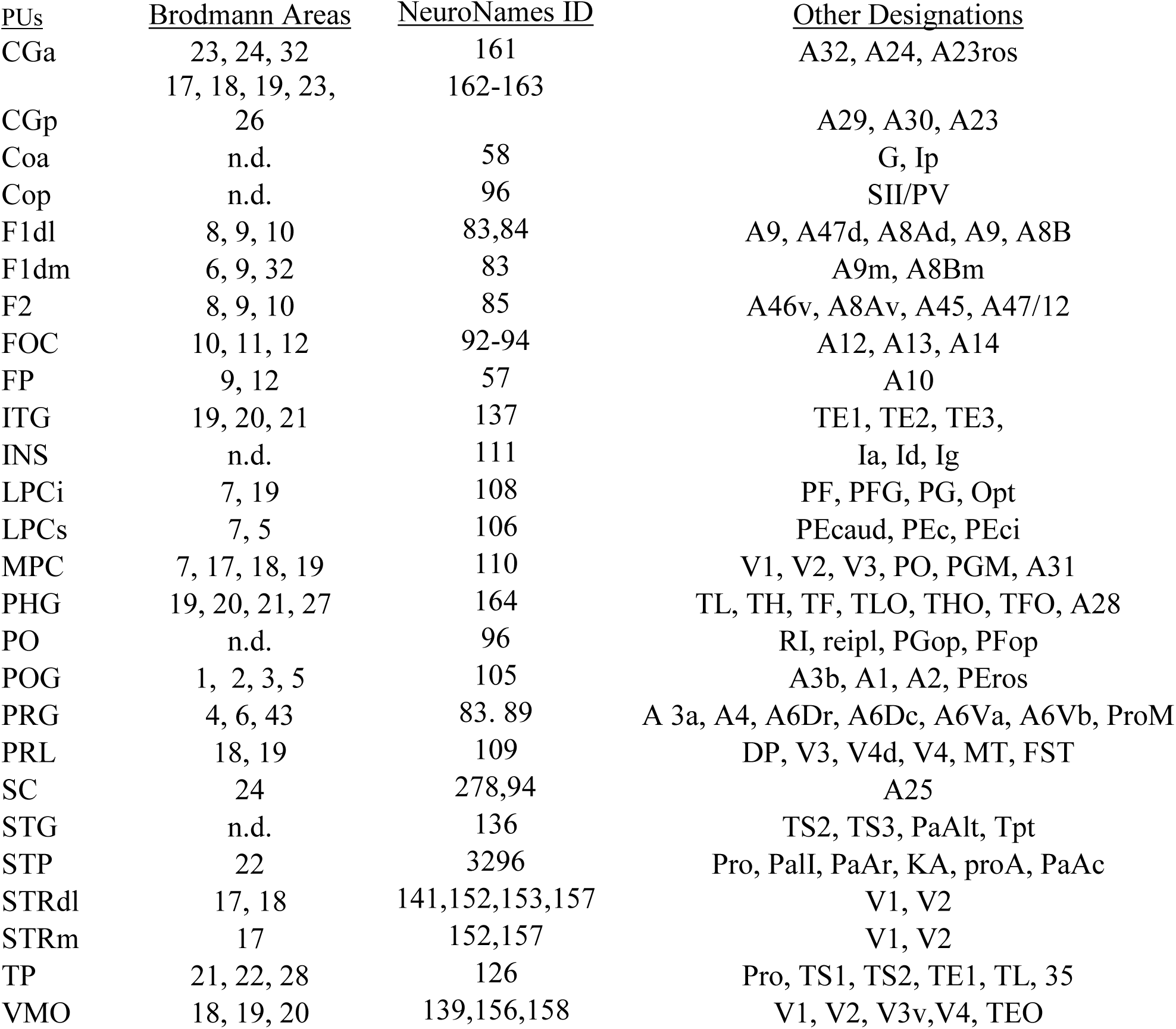
Functional and structural demarcations of parcellation units

#### 1. Coronal Limiting Planes

Coronal plane A is defined by the anterior tip of the rostral sulcus on medial aspect of the brain. It then extends from the medial surface to the inferior, lateral and superior brain surfaces. It forms the caudal boundary of the frontal pole (FP) parcellation unit.

Coronal plane B is defined based on the terminus of the superior ramus of the arcuate sulcus. From this point, coronal plane B extends superiorly to the medial aspect of the hemisphere to end in the cingulate sulcus. This plane forms the posterior border of both the dorsomedial and dorsolateral superior frontal parcellation units (F1dm, F1dl).

Coronal plane C extends from the rostral-most portion of the genu of the corpus callosum to the inferior hemispheric margin. It extends on the ventral aspect of the frontal lobe to end in the olfactory sulcus. This plane separates the subcallosal area (SC) caudally from the fronto-orbital cortex (FOC) rostrally and laterally and the anterior cingulate gyrus (CGa) rostrally.

Coronal plane D progresses ventrally from the tip of the inferior ramus of the arcuate sulcus to the inferior margin of the hemisphere, and separates the inferior frontal cortex parcellation unit (F2) from the precentral gyrus parcellation unit (PrG).

Coronal plane E is defined by a coronal plane rostral to the fronto-temporal junction (i.e., where the frontal lobe adjoins to the temporal lobe in proximity to the limen insulae). It extends around the tip of the temporal lobe to delimit the temporal pole parcellation unit (TP).

Coronal plane F extends from the inferior tip of the central sulcus to the Sylvian fissure. It separates the precentral gyrus (PrG) from the postcentral gyrus (PoG). This plane extends onto the superior lip of the Sylvian fissure to end in the superior part of the circular sulcus of the insula, thus also separating the opercular area anterior to the central sulcus (anterior central opercular (COa) from the opercular area posterior to the central sulcus (COp) (Makris et al., 2004).

Coronal plane G extends from the superior tip of the central sulcus over the interhemispheric fissure to the medial cerebral surface. It progresses through the cingulate sulcus and ends in the callosal sulcus. This plane separates the precentral (PrG) and post central (PoG) parcellation units on the dorsal aspect of the midsagittal surface, and inferiorly separates the anterior cingulate gyrus (CGa) from the posterior cingulate gyrus (CGp) on the medial hemispheric surface.

Coronal plane H is defined by a plane originating from the rostral tip of the intraparietal sulcus. This plane connects this tip to the depth of the Sylvian fissure inferiorly and thus creates a border between the postcentral gyrus (PoG) and the rostral aspect of the inferior lateral parietal cortex (LPCi). In the parietal operculum, it also separates the posterior central opercular parcellation unit (COp) from the parietal opercular (PO) parcellation unit.

Coronal plane I is located at the rostral tip of the calcarine sulcus under the splenium of the corpus callosum. It defines the borders of several parcellation units: 1) on the parahippocampal gyrus it is the border between the parahippocampal (PHG) and ventromedial occipital (VMO) parcellation units; 2) on the dorsal aspect of the cerebrum, this plane extends from the intraparietal sulcus superiorly, passing through the post-central dimple and ending at the cingulate sulcus to separate the post-central gyrus (PoG) parcellation unit from the superior aspect of the lateral parietal (LPCs) parcellation unit; and 3) it forms an intersection with oblique plane a (see below) to form part of the anterior border of the medial parietal (MPC) parcellation unit.

Coronal plane J is formed by the rostral tip of the inferior occipital sulcus. This plane extends ventrally to separate the inferior temporal gyrus (ITG) from the lateral aspect of the ventromedial occipital area (VMO). On the medial aspect of the hemisphere, this plane separates the ventral aspect of the medial parietal cortex (MPC) from the posterior cingulate (CGp) parcellation units.

Coronal plane K is formed by the inferior tip of the lunate sulcus dorsally and extends ventrally to intersect the inferior occipital sulcus. It separates PRL from STRdl in the lateral aspect of the occipital lobe.

Coronal plane L is formed by the end of the parieto-occipital sulcus and extends to the ventral and ventrolateral surface of the hemisphere to intersect with the caudal portion of the inferior occipital sulcus. It separates VMO from STRdl on the lateral, medial and ventral aspects of the occipital lobe, and comprises the rostral border of the STRm parcellation unit.

#### 2. Non-coronal limiting planes

Plane a extends rostrally in a curvilinear fashion from the subparietal sulcus to follow the contour of the cingulate gyrus until it reaches coronal plane I. It thus forms part of the superior border of the posterior cingulate parcellation unit (CGp) with the medial parietal cortex (MPC).

Plane b is a horizontal plane that extends anteriorly from the anterior tip of the inferior occipital sulcus until its intersection with the superior temporal sulcus, thus separating the prelunate parcellation unit (PRL) from the inferior temporal gyrus (ITG) parcellation unit.

Plane c is an oblique line between the superior-posterior tip of the Sylvian fissure and the confluence point of the lunate sulcus with the intraparietal sulcus. This line also passes through the superior tip of the superior temporal sulcus. It forms the ventral boundary of the inferior portion of the lateral parietal cortex (LPCi), and the superior border of the superior temporal gyrus (STG) and the prelunate area (PRL).

Planes d and e extend rostrally from the tips of the superior and inferior calcarine sulci until they intersect coronal plane L. These limiting planes thus provide the superior and inferior borders of the STRm parcellation unit.

### Morphometric Analysis

Parcellation units were identified by the sulcal and limiting planes defined above, and the sum of the voxels for each unit was used to generate volumes for each area on each side of the brain (Figure 6, Table 4). Total cerebral volumes were compared to values calculated from Blinkov and Glezer (1968), and to a subset of rhesus macaque MRI volumetric data (Makris et al., 2010). For the former comparison, average cortical thickness and total surface area of the rhesus macaque cerebral cortex were used to calculate volume, after correcting for fixation, as shown in detail in Table 4. Results of this comparison show that the individual hemispheric volume estimates from the present study were within 9.8-11% of the value estimated from Blinkov & Glezer (1968), and well within the published range of variation for young and old rhesus macaque monkeys (total hemispheric gray matter volume in current study = 44.6 cm^3^; young monkeys: 46 cm^3^ ±6.8 (SD), old monkeys: 40.5 cm^3^ ±4.7 (SD) cf. Table 1, Makris et al., 2010).

**Figure 6.**
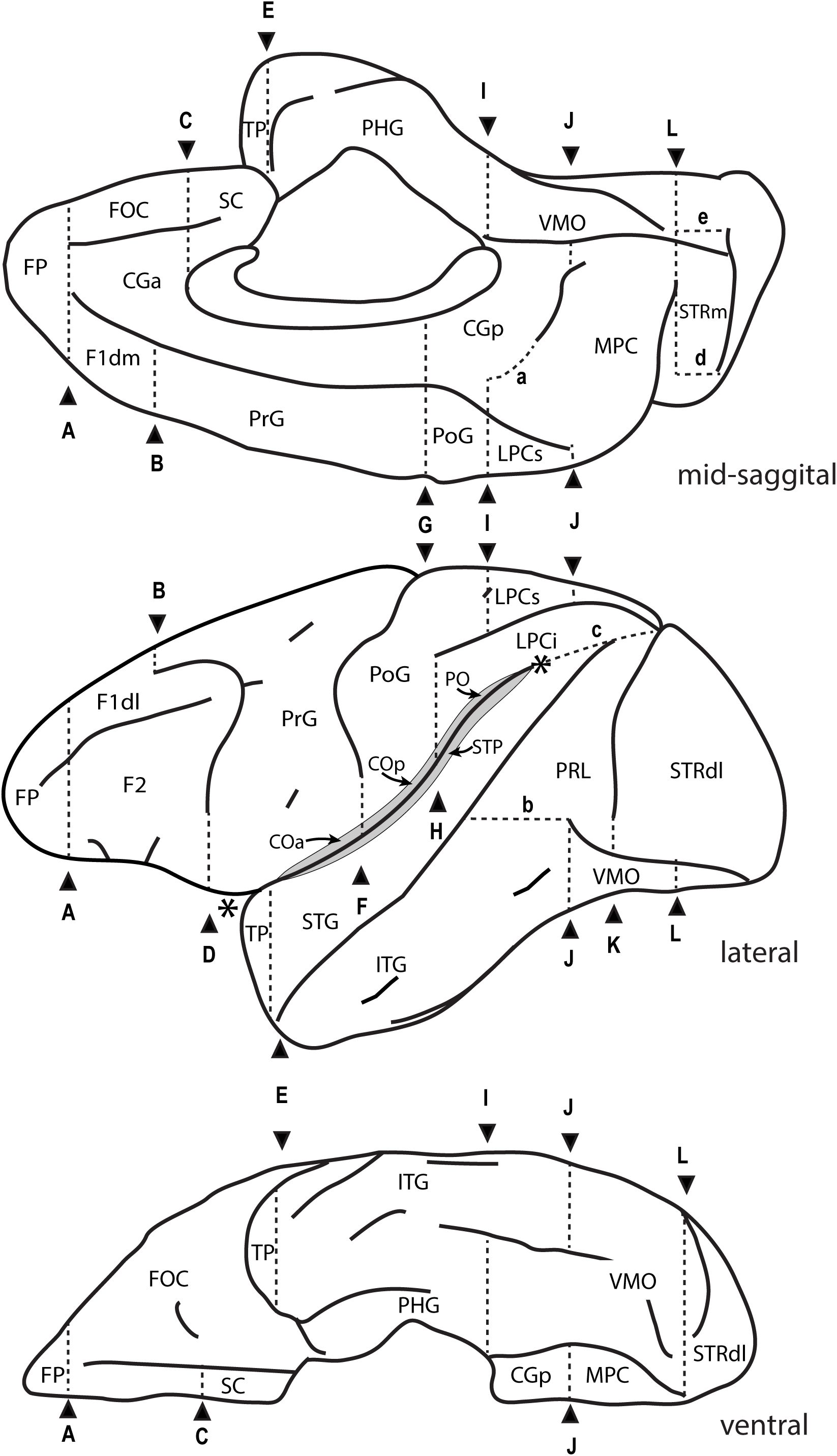
Parcellation units (PUs) in the mid-sagittal (upper), lateral (middle), and ventral (lower) views. The vertical lines (A-L) represent coronal limiting planes, and non-coronal planes are denoted by lower-case letters (a-c). Areas within the insular cortex are displayed in shading. Abbreviations: CGa:anterior cingulate, CGp:posterior cingulate, COa: central operculum-anterior, COp: central operculum – posterior, F1dl: dorso-lateral superior frontal, F1dm: dorso-medial superior frontal, F2: inferior frontal gyrus, FOC: fronto-orbital cortex, FP: frontal pole, ITG: inferior temporal gyrus, LPCi: inferior portion of lateral parietal cortex, LPCs: superior portion of lateral parietal cortex, MPC: medial parietal cortex, PO: parietal operculum, PoG: postcentral gyrus, PRL: prelunate gyrus, PHG: parahippocampal gyrus, PrG: precentral gyrus, SC: subcallosal area, STG: superior temporal gyrus, STP: supratemporal plane, STRdl: dorsolateral portion of striate cortex, STRm: medial portion of striate cortex, TP: temporal pole, VMO: ventromedial occipital

**Table 4:** Parcellation Unit Volumes

| *Parcellation Unit* | *Left Volume (mm<sup>3</sup>)* | *Left volume (%)* | *Right Volume (mm<sup>3</sup>)* | *Right volume (%)* |
| --- | --- | --- | --- | --- |
| Frontal Lobe |  |  |  |  |
| COa | 200.2 | 0.9 | 186.9 | 0.8 |
| F1dl | 808.0 | 3.6 | 899.7 | 4.1 |
| F1dm | 248.0 | 1.1 | 277.3 | 1.3 |
| F2 | 654.2 | 2.9 | 584.3 | 2.7 |
| FOC | 1113.3 | 4.9 | 1162.08 | 5.3 |
| FP | 376.3 | 1.7 | 299.2 | 1.4 |
| PRG | 2815.5 | 12.5 | 2713.3 | 12.3 |
| Total Frontal Lobe | 6215.5 | 27.6 | 6122.8 | 27.8 |
| Occipital Lobe |  |  |  |  |
| STRdl | 2840.8 | 12.6 | 2653.4 | 12.0 |
| STRm | 492.0 | 2.2 | 472.3 | 2.1 |
| VMO | 1079.0 | 4.8 | 1066.9 | 4.8 |
| Total Occipital Lobe | 4411.8 | 19.6 | 4192.6 | 19.0 |
| Temporal Lobe |  |  |  |  |
| ITG | 1739.5 | 7.7 | 1771.0 | 8.0 |
| PRL | 933.6 | 4.1 | 906.1 | 4.1 |
| STG | 1333.6 | 5.9 | 1380.8 | 6.3 |
| STP | 639.8 | 2.8 | 614.2 | 2.8 |
| TP | 385.0 | 1.7 | 426.2 | 1.9 |
| Total Temporal Lobe | 5031.5 | 22.3 | 5098.4 | 23.1 |
| Parietal Lobe |  |  |  |  |
| COp | 201.6 | 0.9 | 190.6 | 0.9 |
| POG | 887.7 | 3.9 | 857.1 | 3.9 |
| LPCi | 947.4 | 4.2 | 983.2 | 4.5 |
| LPCs | 605.3 | 2.7 | 595.4 | 2.7 |
| MPC | 1357.1 | 6.0 | 1275.5 | 5.8 |
| PO | 344.0 | 1.5 | 364.2 | 1.7 |
| Total Parietal Lobe | 4343.0 | 19.3 | 4265.9 | 19.3 |
| Medial Paralimbic |  |  |  |  |
| CGa | 802.9 | 3.6 | 722.7 | 3.3 |
| CGp | 544.2 | 2.4 | 485.1 | 2.2 |
| PHG | 629.0 | 2.8 | 625.6 | 2.8 |
| SC | 97.6 | 0.4 | 87.0 | 0.4 |
| Total Medial Paralimbic | 2073.6 | 9.2 | 1920.5 | 8.7 |
| Insula | 464.6 | 2.1 | 449.1 | 2.0 |
| **Total Cerebral Cortex** | 22540.2 | 100.0 | 22049.3 | 100.0 |
| Cerebral Cortex –<br>Calculated (Blinkov &<br>Glezer, 1968) | 20085 |  | 20085 |  |
| Percent Difference | 11.0% |  | 9.8% |  |

### Ontology

There are several approaches to the systematic codification of neuroanatomical terminology. The Federative International Programme for Anatomical Terminology (FIPAT) is a well-known set of human anatomical terminologies, including the Terminologia Anatomica (2017), the Terminologia Histologica (2008), the Terminologia Embryologica (2017) and the Terminologia Neuroanatomica (2017) (ten Donkelaar et al., 2018). The goal of the FIPAT initiative has been to systematically codify anatomical terminologies to create a common framework, taxonomy, and reference to describe human anatomy, embryology and neuroanatomy. Another approach to codify the ontology of the human nervous system is that of Swanson (2015), which takes a developmental perspective. Both of these systems are focused on human neuroanatomy ontology.

A project that has emphasized both human and non-human primate ontology is the NeuroNames initiative (Bowden & Martin, 1995, Martin & Bowden,1996; Bowden & Dubach, 2003; Bowden et al., 2012). This approach emphasizes the interrelationships of neuroanatomical structures among species, and is therefore of utility in contextualizing the present parcellation framework. To this end, we include a taxonomic structure of how our parcellation units relate to cerebral cortical landmarks and terminologies derived principally from the NeuroNames framework (Table 5). The structure and divisions outlined in Table 5 and Figure 8 are in large part consistent with the human-based ontologies referenced above, excepting certain species-specific differences.

**Figure 7:**
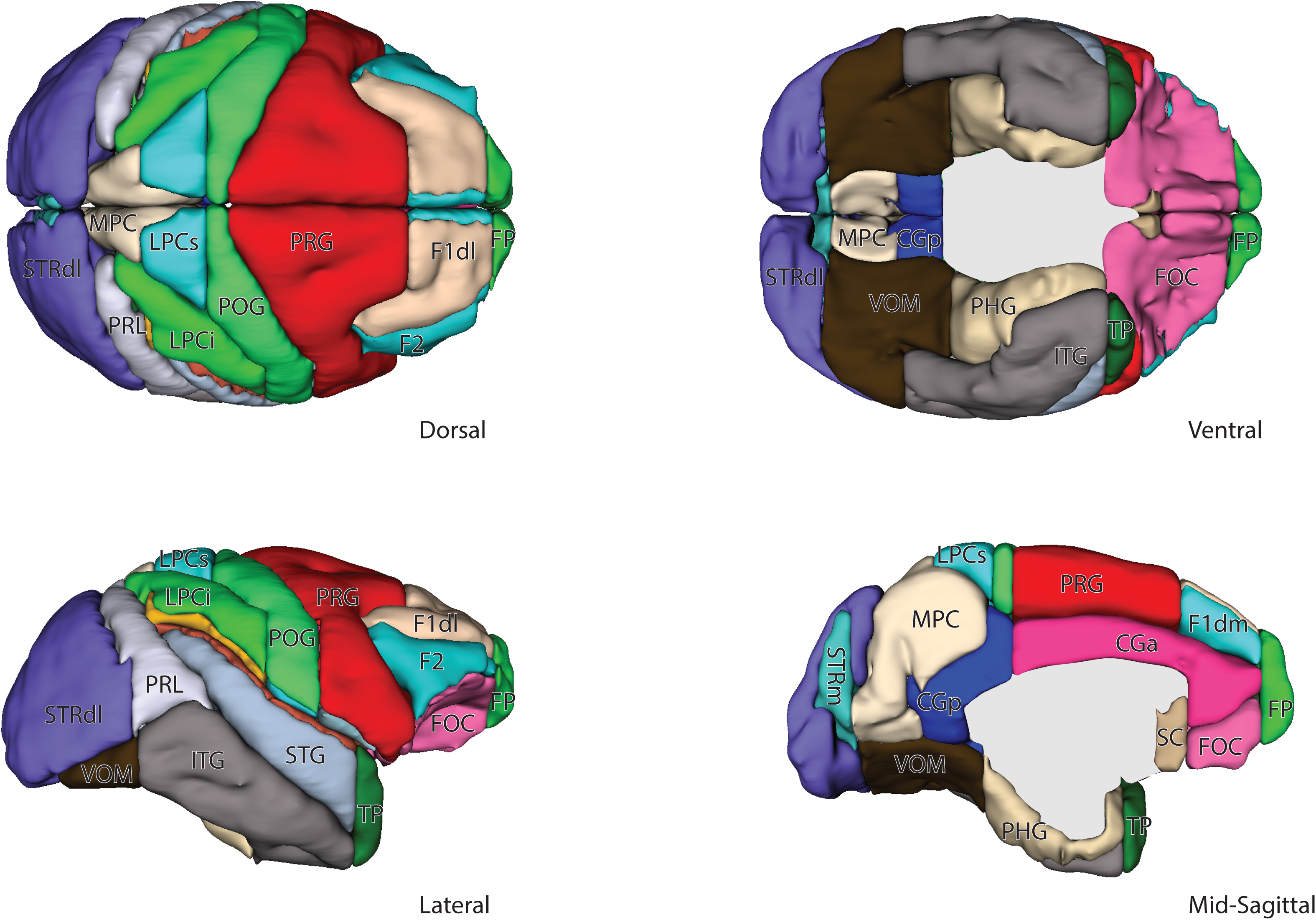
Three-dimensional view of parcellation units in the four cardinal views of the macaque brain. Abbreviations as in Figure 6.

**Figure 8.**
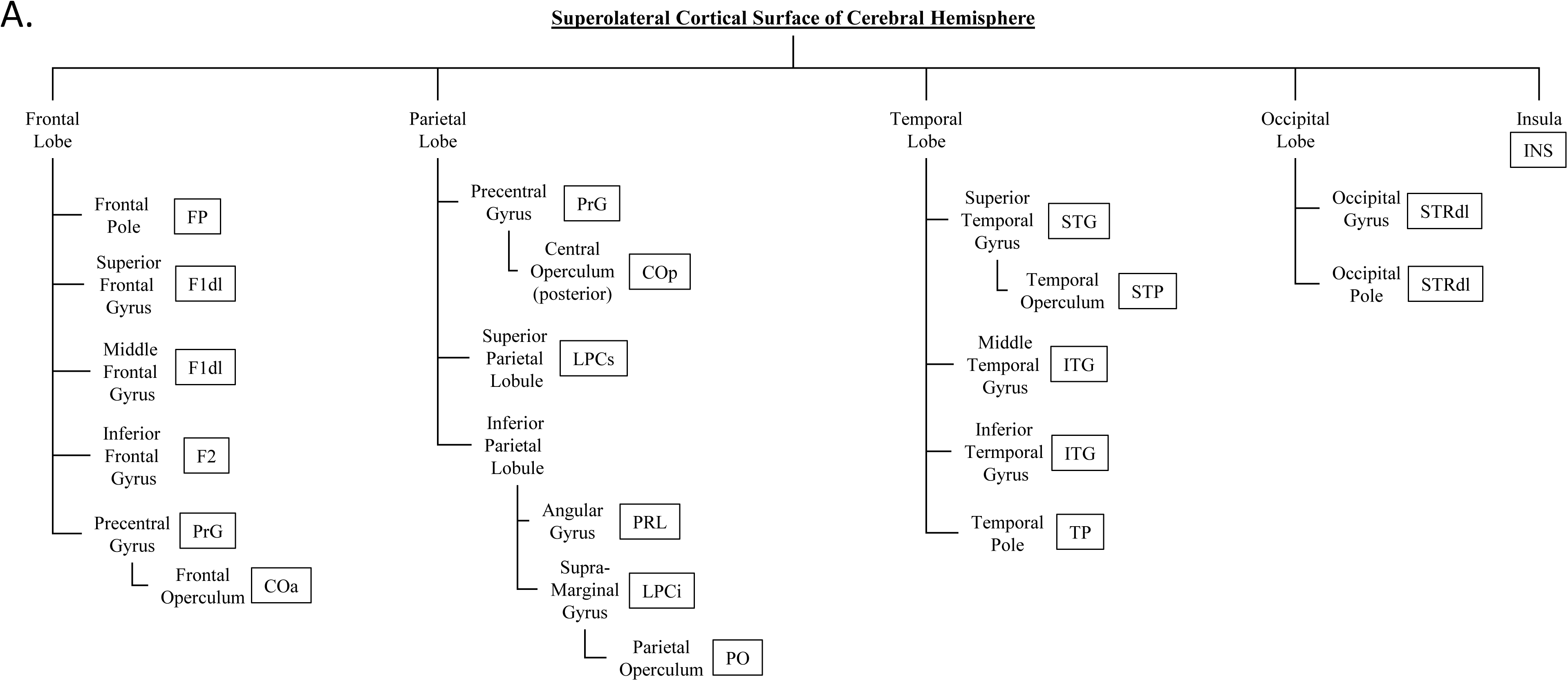
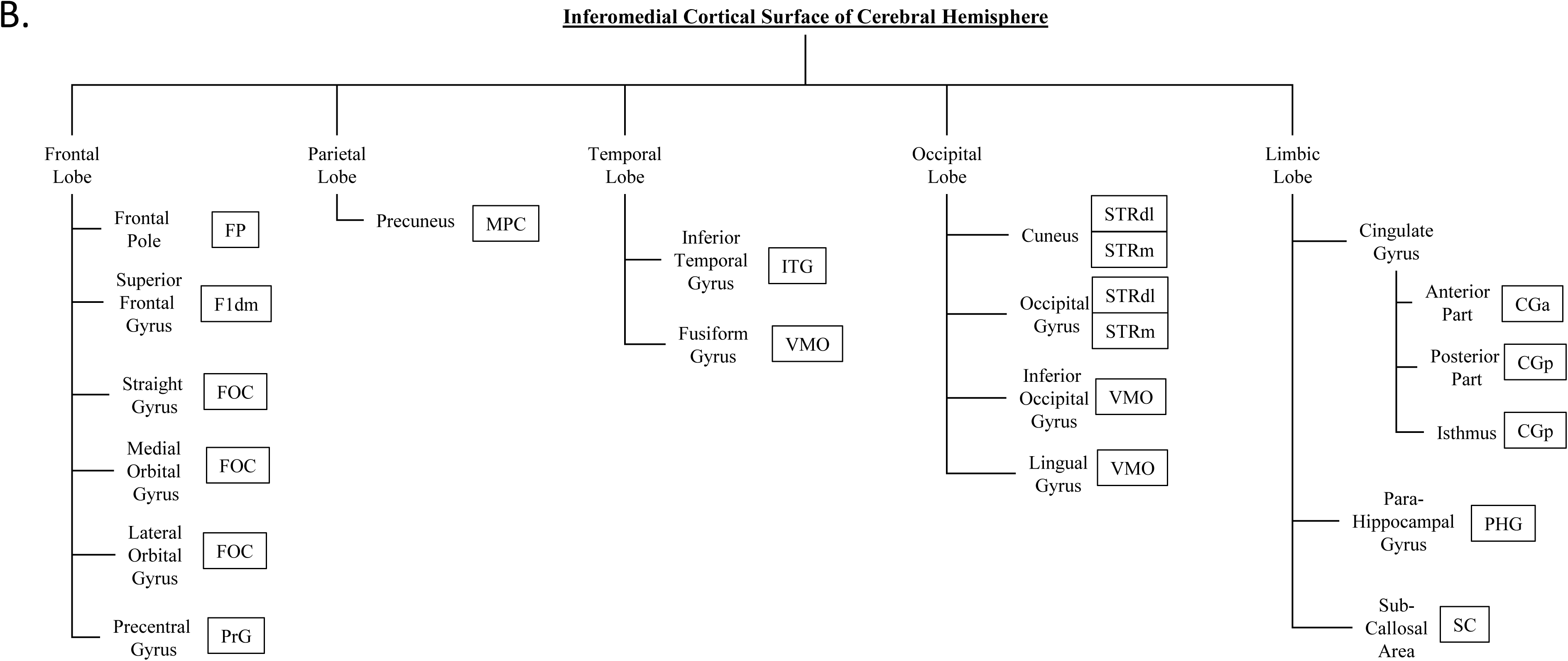
Visual representation of relationships between taxonomical structures (regions) tabulated in Table 5. Parcellation units are boxed to the right of the anatomical structure with which they are associated.

**Table 5:**
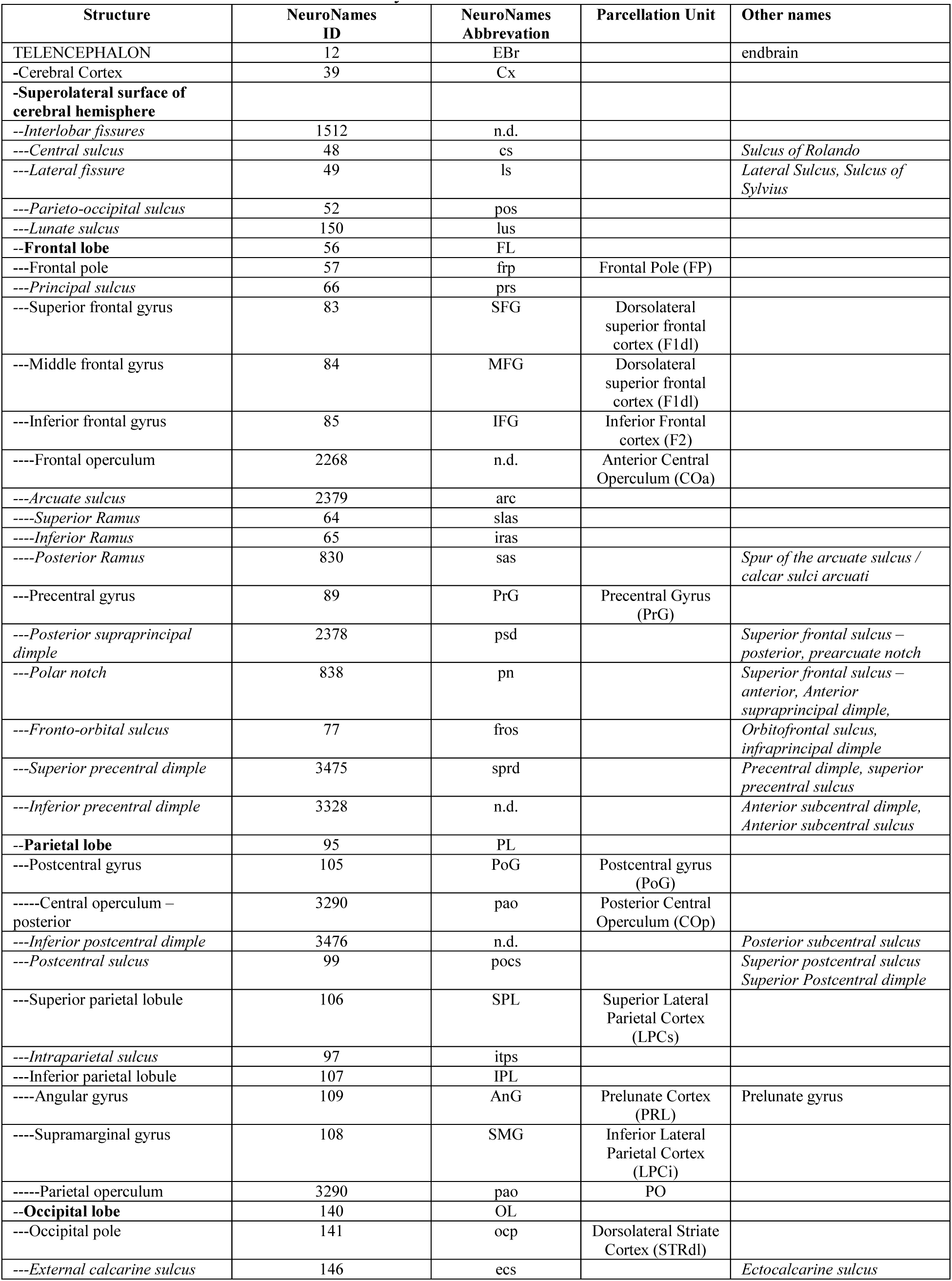
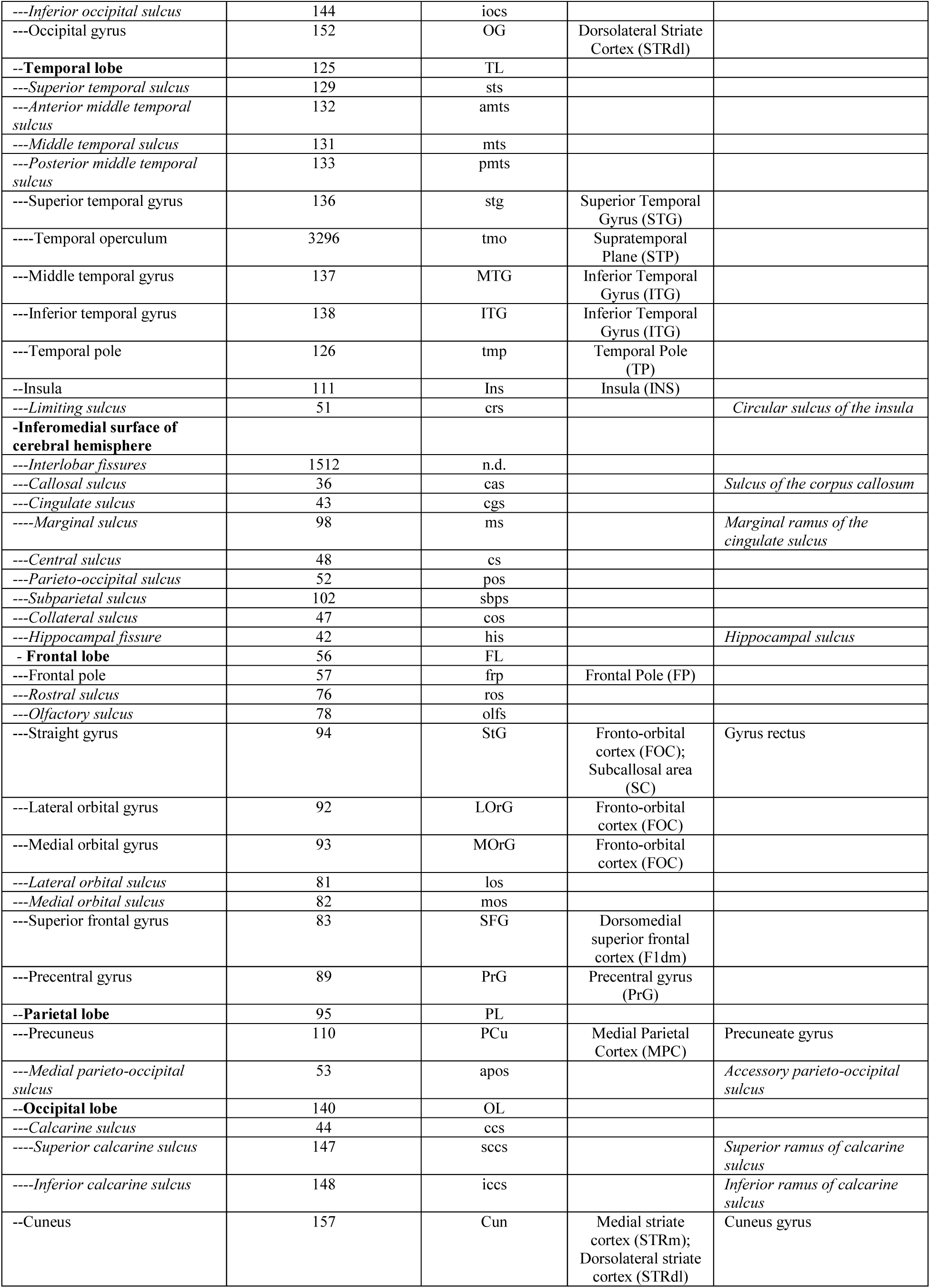
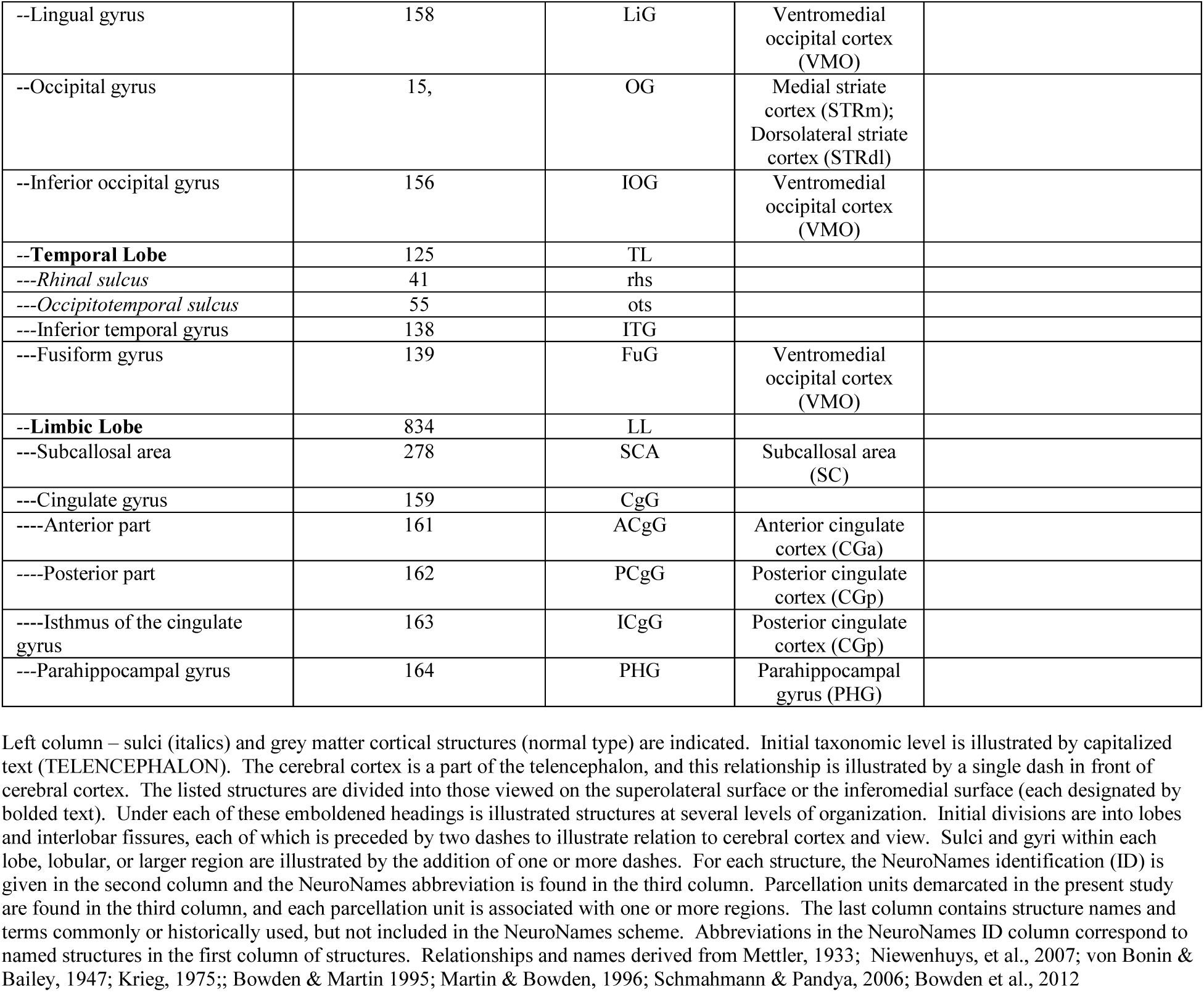
Table of structures in the rhesus monkey cerebral cortex.

## DISCUSSION

The present system parcellates the rhesus macaque brain according to specific anatomical boundaries. These boundaries delimit broad anatomical regions of interest (ROIs) or parcellation units (PUs) that then can be evaluated through morphometric criteria such as quantitative measures of volume or thickness, and can be used in future studies as a structural basis upon which different types of data can be overlaid (e.g., fMRI, dMRI tractography, magnetoencephalography). This approach is based on similar studies using anatomically reliable borders in the human brain (Rademacher et al., 1992; Caviness et al., 1996; Makris et al., 1999), and separates the cortical surface into discrete areas that belong to neural systems and ultimately are composed of areas with similar hodological, architectonic and functional characteristics. For the purpose of illustration, this theoretical approach was applied here in one rhesus monkey to demonstrate feasibility. We emphasize that although the parcellation map of the single macaque subject presented herein could be used as a brain atlas, the principal relevance of this investigation lies with the development of a process for generating a cortical map based on individual morphological anatomy. The latter is the key feature for precision neuroanatomy in the macaque and also a necessary component to complete the validation of the macaque brain circuit diagram (BCD) using neuroimaging and, importantly, to improve through homology the formulation of the BCD in humans.

### Parcellation validity

The parcellation units, as specified above, consist of cerebral cortical regions that have been defined first by architectonic criteria, and in some cases refined by physiological and/or immunohistochemical markers (e.g., Saleem & Logothetis, 2007). The cytoarchitectonic divisions have been recognized as a foundational aspect of connectivity (e.g., Pandya & Yeterian, 1985; Mesulam, 1988, 2000; Pandya et al., 2015; Hilgetag et al, 2016; Beul et al., 2017; Goulas et al., 2018). These structural features underpin the concept that brain regions are organized into neural systems that mediate specific aspects of behavior (e.g., Mesulam, 2000; Pandya et al., 2015). At a fundamental level, therefore, a basic unit for the analysis of systems-level function is the delineation of cytoarchitectonic areas. At the current level of technology, MRI is unable to resolve the cytoarchitectonic fields *in vivo.* As a result, the specific designation of the present parcellation units based on sulcal patterns and limiting planes in the monkey brain presumes that these boundaries represent, to varying degrees, brain areas cytoarchitectonically defined by previous investigators (e.g., Brodmann, 1909; von Bailey & Bonin, 1947; Sanides, 1962; Rademacher et al., 1992, 1993). It should be noted that beyond primary cortical areas, the relationship between cytoarchitectonically-derived boundaries and sulcal and gyral patterns is increasingly variable in the primate brain (e.g., Ebberstaller, 1890; Cunningham, 1892; Bailey & von Bonin, 1951; Sanides, 1962; Eidelberg & Galaburda, 1984; Ono et al., 1990; Rademacher et al., 1993).Thus, there is an absence of a 1:1 correspondence between parcellation units and cerebral cortical cytoarchitecture, a fact also reflected in the ontological affiliation of the parcellation units. This morphometric approach is the basis of cortical parcellation for the Harvard-Oxford Atlas in humans (Rademacher et al., 1992, 1993; Filipek et al., 1994; Caviness et al., 1996).

### The macaque Harvard Oxford Atlas (mHOA) Approach to Morphometric Analysis

Although segmentation and parcellation of cortical regions based on anatomical landmarks has been performed in human structural MRI studies following the HOA guidelines (Rademacher et al,. 1992; Fillipek et al., 1994; Caviness et al., 1996), this has not yet been comprehensively accomplished for the macaque monkey. As in the human, such a system has a potential to guide fundamental morphometric studies and questions of aging, sex differences, interindividual variability, and asymmetry in normality and disease. Additionally, systematic parcellation of the cortical surface can provide a topographical framework for the mapping of other data modalities, including functional imaging data, multisite electrophysiological data, magnetoencephalography, and diffusion MRI tractography. These topographical maps are critical components in data analysis using current mathematical and statistical models (Bullmore & Sporns, 2009, Lynall et al., 2010).

The cortical parcellation approach used in the present paper also has the potential to inform the human BCD. Due to technical and ethical considerations, much of the knowledge of human neuroanatomy is inherently comparative and is based on invasive experimental results from other species (see preceding paper). This homological approach means that human brain connectivity lacks the gold standard needed as a solid basis to build the BCD (Fig 1-3). A comprehensive BCD of the macaque monkey would take advantage of the results of decades of invasive neuroanatomical data and would use homology to advance the human BCD. This paper provides one of the two components of such a macaque BCD, namely a description of the system used to derive a cortical parcellation map based on a systems-level approach. Furthermore, it addresses ontological definitions including a comparative approach to cortical areas, a topic of current interest and relevance in basic neuroscience (Martin & Bowden, 1996). What follows is a description of the different elements of the cortical parcellations. Given that this is the first paper of its kind in the macaque monkey, namely one that addresses both computational neuroscientists and neuroanatomists involved with imaging, we considered it relevant to discuss in some detail the neural systems underlying the parcellation units.

### Functional and Anatomical Zones

The system detailed above allows for a way to investigate the relationship between brain structure, as reflected in the division of the cortical surface into parcellation units and associated functions. A number of structural maps based on cytoarchitecture (e.g., Brodmann, 1909; von Bonin and Bailey, 1947) have provided initial subdivisions and a coarse-grained division of regions. Subsequent studies built upon and further refined the maps based on hodological and physiological properties (e.g., Vogt et al. 2015, Pandya & Yeterian, 1985; Glasser et al. 2011, Van Essen et al., 2012; Glasser et al. 2016). The perfect ontological system has yet to be established as organizational principles based on function and structure are still being brought to terms, but a broad organizational scheme can be elaborated based on the existence of functional systems.

A classical way to think about brain systems is the characterization of various regions as performing one of three different kinds of general function (Rademacher et al., 1992; Mesulam, 2000). Primary regions can be classified either as sensory or motor, and constituent neurons are limited to the processing of a single modality (e.g., vision, audition, motor). These regions are characterized by a strong connection to the periphery. Primary sensory areas contain a map of sensory space produced by dense and organized input from relevant sensory receptors and these areas are the first regions to receive this high density information. By contrast, the primary motor area has a topographically mapped organization and sends outputs to command discrete muscular activation. Linked to the primary sensory regions are unimodal association areas. Neurons in these areas receive signals from individual primary sensory areas, and further process and elaborate signals into more complex representation of sensory space and objects within (Pandya & Kuypers, 1969; Powell & Jones, 1970). These regions process a single modality, and receive very limited input from other modalities. Unimodal association areas of the motor cortex have neurons that house more complex motoric representations, which are then relayed to the primary motor cortex for deconvolution into more component actions. Finally, heteromodal association areas represent a further elaboration of signals by merging processing from one or more modalities. For example, heteromodal regions house multisensory neurons that respond to both auditory and visual signals, or have neurons that respond to visual information as well as relaying motor commands.

#### Primary Cortices

##### a. Primary Auditory Cortex

The primary auditory cortex (AI) is a region located in the inferior opercular surface of the lateral (Sylvian fissure). It is characterized by a prominent middle granular layer (koniocortex), and it receives a large number of projections from the ventral portion of the medial geniculate nucleus of the thalamus (vMGN; Pandya & Sanides, 1973; Galaburda & Pandya, 1983; Cipilloni & Pandya, 1991; Morel et al., 1993; Kosaki et al., 1997; Kaas & Hackett, 2000; Scott et al., 2017). This region is located in the STP parcellation unit.

##### b. Primary Visual Cortex

The visual cortex in the macaque monkey has the most elaborate cytoarchitecture of any cerebral area, with a highly granular layer 4 subdivided into five discrete layers based on cytoarchitecture (Polyak, 1957) and physiology (Hubel & Wiesel, 1969, 1974a, b). The primary visual cortex is the single largest architectonic region of the cerebral cortex, and occupies the majority of the lateral aspect of occipital lobe, from the lunate sulcus to the occipital pole. It continues over the occipital pole to reach the medial aspect of the hemisphere slightly beyond the dorsal and ventral rami of the calcarine sulcus, and extends rostrally into the depths of the stem of the calcarine sulcus. This region corresponds to area 17 of Brodmann (1909) and area OC of von Bonin & Bailey (1947). The projection of the visual field onto the surface of area 17 has been well mapped (Daniel & Whitteridge, 1961; Hubel & Wiesel, 1974; Gattass et al., 1981; Van Essen et al., 1984; Tootell et al., 1988; Polimeni et al., 2006). The central field representation is located on the lateral and rostral border of area 17, with the vertical meridian representation extending from this point superiorly and inferiorly along the caudal bank of the lunate sulcus. The horizontal meridian representation extends caudally from the foveal representation, passes through the occipital pole and turns to progress rostrally into the depth of the calcarine sulcus. While there remains some uncertainty about the precise rostro-medial limits of area V1, maps based on cytoarchitecture (Brodmann, 1909; von Bonin & Bailey, 1947) and physiology (Gattass et al., 1981) place the border slightly rostral to the vertical rami of the calcarine sulcus, where regions representing the most superior and inferior aspects of the contralateral visual hemifield are located. Accordingly, the parcellation unit STRdl represents the majority of the visual representation – extending from the fovea to a peripheral azimuth of approximately 6-10 degrees of visual angle, whereas the medial parcellation unit STRm contains more peripheral representations of the contralateral visual hemifield (>10 degrees). The far peripheral representation is contained within the depths of the calcarine (horizontal meridian representation) and would be contained within ventromedial occipital (VMO) parcellation unit for the peripheral and upper visual field, and the medial parietal cortex (MPC) parcellation unit for the lower peripheral visual field.

##### c. Primary Somatosensory Cortex

The primary somatosensory cortex is a group of several areas in the rostral parietal lobe caudal to the central sulcus, each of which forms a thin strip parallel to the central sulcus. Brodmann’s area 3b (referred to as area PB by von Bonin and Bailey (1947)) has a prominent granular layer IV similar to other primary sensory areas, and is located within the caudal bank of the central sulcus. More caudal regions (areas 1, 2) also receive input from somatosensory thalamus and are considered primary somatosensory areas (Woolsey et al., 1942; Werner & Whitsel, 1968; Paul et al., 1972; Dreyer et al., 1974, Dreyer et al., 1975; Nelson et al., 1980). Areas 3b, 1 and 2 are arrayed in sequential vertical strips moving posteriorly from the caudal bank of the central sulcus. They are characterized functionally by different receptor input and receptive field properties, and cytoarchitectonically by progressively fewer granule cells and more pyramidal cells with increasing distance from the central sulcus. The primary somatosensory regions extend dorsally from the superior aspect of the cingulate sulcus, over the vertex and inferiorly toward the Sylvian fissure. The neurons of each region are organized somatotopically such that the sensory representations of the contralateral foot are on the medial aspect, the leg and trunk representations on the dorsolateral aspect, and the contralateral hand and face representations on the ventrolateral aspect. This organization is observed in parallel for each region (i.e., 3b, 1, 2) of the primary somatosensory cortex. The primary somatosensory cortex is entirely encapsulated in the PoG parcellation unit.

##### d. Other primary sensory areas

The primary gustatory and olfactory regions are located adjacent to the connection of the temporal lobe with the frontal lobe. The primary gustatory region is located dorsally on the rostral part of the frontal operculum (FO parcellation unit; Sanides, 1968; Pritchard et al., 1986), whereas the primary olfactory cortex is found in the caudal portion of the orbitofrontal region (FOC parcellation unit; Powell et al., 1965; Price & Powell, 1971). Vestibular cortex is located in the caudal depths of the lateral fissure in a region known as the parieto-insular vestibular cortex (PIVC) that is located in parcellation unit PO (Akbarian et al., 1994; Chen et al., 2010).

##### e. Motor cortex

The motor cortex, Brodmann area 4 (or area F1 of Matelli and colleagues, 1985, 1991), is characterized by the lack of a noticeable layer IV (Geyer et al., 2000; see also Garcia-Cabezas & Barbas, 2014), and the presence of large pyramidal cells in layer V (Betz cells; Brodmann, 1909). This region lies anterior to and within the depths of the central sulcus, extending in a strip-like fashion from the medial aspect of the hemisphere over the dorsolateral to the ventrolateral cerebral surface. In this way, it parallels the shape and orientation of the primary somatosensory cortex. Similarly, the motor map of the body parallels the organization of the primary somatosensory cortex with the representation of the contralateral leg on the medial surface, trunk on the dorso-lateral cerebral surface, and face and hand on the ventro-lateral cerebral surface. The motor cortex, unlike the somatosensory cortex, is wider in area dorsolaterally, and tapers towards the Sylvian fissure. Along with premotor areas (see below), this region is included in the PrG parcellation unit.

#### Unimodal Association Areas

##### a. Auditory Association Areas

The primary auditory cortex is one of two or three core auditory regions located on the supratemporal plane. This region receives dominant thalamic input from the vMGN and is tonotopically organized. Early studies distinguished a medial koniocortical area (KAm) from a laterally adjacent koniocortical region (KAlt) in area KA (Pandya & Sanides, 1973). Later studies employed a different organization based in part on physiological properties, and situated AI caudally with respect to a more rostral region (area R; Morel et al., 1993). A further rostral area (area RT) differs from A1 and R in its physiological properties and because it receives a smaller projection from the vMGN (Kaas & Hackett, 2000; Scott et al., 2017). These core auditory areas *per se* are strongly interconnected, and also are connected to surrounding auditory association areas that are arrayed as a circumnavigating belt around the core regions. These belt regions extend to the lateral surface of the superior temporal gyrus, as well as rostrally, caudally, and medially, although the extent of the belt regions within the sulcus is debated and the nomenclature varies based on the study and authors (Pandya & Sanides, 1973; Galaburda & Pandya, 1983; Morel et al., 1993; Kosaki et al., 1997). Regardless of the conventions and nomenclature used, it is clear that the vast majority of the supratemporal plane contains core and auditory association regions, the vast majority of which are captured in the STP parcellation unit.

Connectional studies of the auditory association areas of the superior temporal gyrus and supratemporal plane have identified three main streams of projections targeting independent areas in the frontal, parietal, temporal and paralimbic areas (see, e.g., Pandya & Yeterian, 1985). These projections and the regions from which they emerge are conceived to be divided in terms of the information they process and relay. Signals related to auditory object recognition (e.g., sound of objects, species-specific vocalizations) are processed largely in more anterior association regions on the supratemporal plane and the superior temporal gyrus. By contrast, signals related to the location and salience of auditory stimuli in space are processed by regions on the superior aspect of the supratemporal plane and the superior temporal gyrus, and progress to parietal lobe areas that code for space (Kaas & Hackett, 2000; Rauschecker & Tian, 2000; Tian et al., 2001; Romanski et al., 1999; Rauschecker & Scott, 2009; Recanzone & Cohen, 2010; Ortiz-Rios et al., 2017).

##### b. Visual Association Cortex

The occipital lobe of the human and non-human primate was originally elaborated by Brodmann (1909) who divided it based on cytoarchitectonic features into area 17, which later was found to be coincident with the primary visual cortex, and areas 18 and 19. Subsequent studies based on physiological as well as more refined anatomical methods found that areas 18 and 19 housed a substantial number of distinct areas (Van Essen et al., 1981). These areas were typically named according to their proximity to area 17, also known as visual area 1 or V1. Thus, V2 forms a belt surrounding the rostral border of V1, V3 lies in front of V2, and V4 is located rostral to V3. Visual properties become more complex and neurons become increasingly selective to specific types of signals with increasing distance from area V1. Further studies subdivided these areas and indicated that stimulus selectivity was organized such that signals coding for space and movement were processed in more dorsal areas, whereas neurons responsive to specific objects were located in more ventral areas. The series of regions that together elaborate signals of space and movement were referred to as the dorsal or “where” stream, whereas the ventral regions were conceived to be the “what” stream that codes for object properties (Ungerleider & Mishkin, 1982). Dorsal stream visual association areas were found to project largely to heteromodal regions in the parietal lobe, whereas visual association areas of the ventral stream extended into regions of the inferior temporal gyrus to the temporal pole (Gross et al., 1969; Gross et al., 1972; Perrett et al., 1982; Desimone et al., 1984; Colby et al., 1988; Cavada & Goldman-Rakic, 1989a; Boussaoud et al., 1990; Tanaka et al., 1991; Fujita et al., 1992; Kobatake & Tanaka 1994; Andersen, 1995; Galletti et al., 2005; Ungerleider et al., 2008). Further studies suggested a more refined division of the dorsal stream into a dorsal-dorsal stream progressing to the superior parietal lobule and participating in on-line control of movements, and a ventral-dorsal stream leading from visual occipital association areas to inferior parietal regions and subserving the perception and recognition of space and action (Goodale & Milner, 1992; Rizzolatti & Matelli, 2003, also see Galletti et al., 2001, Galletti et al,. 2003).

The PRL parcellation unit contains many of the unimodal visual association areas, including V3, V4, V4t, DP, MT, and FST. The ITG parcellation unit contains many of the high level unimodal association areas of the ventral stream, including TEO, TE3 and TE2. Medial visual association cortices are encapsulated in the ventromedial occipital (VMO) parcellation unit, which contains ventral area V2 and ventral area V3 (V3v), whereas the medial parietal cortex (MPC) parcellation unit contains medial visual association areas including dorsal area V2, dorsal area V3, and area PO.

##### c. Somatosensory Association Cortex

The second somatosensory area (SII) contains its own somatotopic representation and is located posterior and ventral to the primary somatosensory areas, on the post-central parietal operculum (Krubitzer et al., 1995). This region receives input from primary somatosensory areas and subserves haptic perception (Sinclair & Burton, 1993), a function consistent with an object-oriented ventral stream, suggesting that somatosensory cortex may have a functional dichotomy of signals similar to the ventral and dorsal streams in auditory and visual cortices. A supplementary somatosensory area (SSA) located on the caudal inferior rim of the marginal sulcus of the cingulate sulcus also contains its own representation of the contralateral body surface (Morecraft et al., 2004), and is related to processing signals of navigation in space, similar to dorsal stream areas emanating from the auditory and visual unimodal association areas. Intermediate unimodal somatosensory areas in the rostral IPL (area 7b, PF) and SPL (rostral area 5, rostral PE) have responses that reflect the integration of receptor types independently processed in the two streams of the primary somatosensory cortex (Duffy & Burchfiel, 1971; Sakata et al., 1973).

The somatosensory association units are included in different parcellation units, including PO for SII, PoG for rostral area 5, and LPCi for rostral area 7 (PF). Since SSA is located in the marginal sulcus of the cingulate sulcus (Morecraft et al., 2004), it is present in the LPCs parcellation unit. Rostral area 5 is included in the PoG because of its functional similarity to primary somatosensory cortex. However, the rostral part of area 7b (PF) is heteromodal in terms of the responses of its component neurons (Hyvarinen et al., 1974; Leinonen et al., 1979; Dong et al., 1994) and is thus considered below in the heteromodal areas.

##### d. Motor Association Cortex

This collection of areas lies rostral to area 4. Originally defined by Brodmann as area 6 (Brodmann, 1909), these regions have been further elaborated on the basis of function and connectivity. Area 6 contains at least 4 subdivisions, a dorsal premotor area located above the arcuate spur, a ventral premotor area below the arcuate spur, a supplementary motor area (SMA, also known as the second motor area or MII) on the medial aspect of the hemisphere, and a pre-supplementary motor area (pre-SMA) rostral to SMA. The banks of the arcuate sulcus contain the frontal eye field, which is considered the premotor region for eye movements (Bizzi & Schiller, 1970; Latto & Cowey, 1971a,b; Bruce & Goldberg, 1985). Finer divisions of premotor areas based on cytoarchitectonic, connectional and functional differences have been proposed (Matelli et al,. 1985; Matelli et al., 1991; Belmalih et al, 2009) such that mesial, dorsal and ventral premotor regions contain at least two each. All of these regions project strongly to the primary motor cortex, and together function to contextualize and coordinate motor commands (e.g., Sasaki & Gemba, 1986). These regions, while unimodal in the sense that they process and relay motor commands, use visual and somatosensory information to integrate movements with hand shape to appropriately grasp an object, or with the position of the arm in space to correctly target reaching movements (Murata et al., 1996; Bonini et al., 2012). Many of the neurons of these regions respond to these goal-directed actions whether they are carried out directly, or simply observed to occur (mirror neurons; Rizzolatti et al., 1996; Raos et al, 2006; Bonini et al., 2014). The premotor areas, therefore, may be better classified as heteromodal (Bruni et al., 2017), even though previous approaches have considered them to be unimodal motor association areas. These premotor areas, together with the primary motor cortex (BA4), are included in the PrG parcellation unit.

#### Heteromodal Association Cortices

These regions are critical for integrating signals across sensory modalities, and also for integrating sensory signals, motor commands, and signals from limbic regions. Heteromodal areas contain interdigitated neurons with separate modality preferences, but also neurons that respond to multiple modalities and thus not linked to a single modality (Mesulam, 2000). Heteromodal regions are typically located where the outer rim of unimodal association areas from auditory, somatosensory and visual areas adjoin each other, namely in temporal and parietal and insular areas but also in prefrontal and paralimbic regions.

##### a. Prefrontal Association Cortex

Prefrontal regions are located rostral to the premotor cortices. For the purposes of the present parcellation scheme, the lateral prefrontal surface was separated into a dorsal and ventral portion based on the location of the principal sulcus, a dividing line that is largely respected based on connectional, cytoarchitectonic, and functional bases (e.g., Pandya & Yeterian, 1985; Yeterian et al., 2012; Pandya et al., 2015). In particular, unimodal association areas involved with perception and processing of objects project to lateral prefrontal areas inferior to the principal sulcus. By contrast, dorsally located unimodal association areas involved in the perception of space and the production of spatially accurate motor programs target prefrontal areas superior to the principal sulcus (Pandya & Yeterian, 1985; Pandya et al., 2015). Thus conceived, the more dorsal F1dl parcellation unit includes areas 8B, 8Ad, 9 and dorsal 46, whereas F1dm includes the medial portions of 8B and 9. The F2 parcellation unit includes areas 47/12, ventral 46, 8Av, and 45.

The fronto-orbital cortex (FOC) parcellation unit includes areas on the ventral surface of the frontal lobe, and extends from the lateral hemispheric margin ventrally and around the medial hemispheric margin to end in the rostral sulcus. This region includes areas 11, 13, and 14. These orbitofrontal regions are connected to unimodal association areas that respond to modality-specific object representations (e.g., Ungerleider & Mishkin, 1982; Pandya & Yeterian, 1985; Barbas & Pandya, 1989; Morecraft et al., 1992; Barbas, 1993; Carmichael & Price, 1995; Kaas & Hackett, 1999; Rolls et al., 2003). The orbitofrontal areas also connect with paralimbic cortex relating to affective visceral input, and receive gustatory and olfactory input, with some component neurons being bimodal for taste, gustation or vision (Rolls & Baylis, 1994; Rolls et al., 1996). These characteristics suggest that the orbitofrontal regions confer emotional valence or reward value relating to object identity, as defined through visual, gustatory, and olfactory input, or through ‘mouth-feel’ (Rolls et al., 2003). The frontal pole (FP) parcellation unit corresponds to area 10, an area thought to provide control of emotional regulation and visceromotor tone through projections to anterior cingulate (and the subcallosal region; see Joyce & Barbas, 2018) and to play a role in high level auditory processing (Medalla & Barbas, 2014).

##### b. Heteromodal association cortices of the temporoparietal junction

On the lateral aspect of the parietal cortex, a swath of cortical areas exists between auditory, visual and somatosensory association areas. This region is located mainly in posterior parietal cortex (superior and inferior parietal lobules) and extends to the temporal pole via the banks of the superior temporal sulcus, and to the medial surface via the medial parietal cortex. Cortical areas in this region contain mixed populations of modality-specific neurons (e.g., neurons that respond to visual stimuli adjacent to neurons that respond to auditory stimuli), and neurons that respond to more than one modality (i.e., visual, auditory, and coincident visual and auditory neurons). The functions of the areas in this region vary according to their location in the cortex.

The posterior parietal cortex on the superior parietal lobule (caudal Brodmann area 5, or areas PE, PEc), including medial parietal areas (area PGm, Brodmann area 31) contain neurons responsive to goal-directed movement and monitoring of the limbs or the eyes in space (e.g., Bakola et al., 2010, Baloka et al., 2013; Hadjidimitrakis et al., 2015; Piserchia et al., 2017). This region receives input from the dorsal ‘what’ streams of auditory, visual and somatosensory areas, and project strongly to dorsal premotor and prefrontal areas (Pandya & Yeterian, 1985). The parcellation unit LPCs includes these regions.

Regions of the inferior parietal lobule (area 7, PFG, PG, OPT) are important for spatial processing, temporal processing, aspects of decision making, as well for visuomotor integration for arm, hand and mouth (Hyvarinen, 1981; Gottlieb et al., 1998; Andersen, 1995; Platt & Glimcher, 1999; Leon & Shandlen, 2003). As such, they receive visual, auditory, vestibular and somatosensory input from both dorsal and ventral stream areas and project to ventral and dorsal banks of the principal sulcus along with parahippocampal and cingulate regions (Cavada & Goldman-Rakic, 1989a,b; Andersen et al., 1990; Friedman & Goldman-Rakic, 1994; Stanton et al., 1977).

Regions of the superior temporal sulcus are heteromodal in nature, serving as high level multimodal processing stations for object characterization, identity and processing. These areas project to ventral and orbital prefrontal areas, as well as to paralimbic regions in the parahippocampal and posterior cingulate cortices (Pandya & Yeterian, 1985; Pandya et al,. 2015).

#### Paralimbic association cortices

The paralimbic association cortices are proisocortical areas that represent a cytoarchitectonic transition between allo- and periallocortical (3-4 layered) limbic structures such as the primary olfactory (piriform) cortex and the hippocampus, and the isocortical (6-layered) neocortex. These paralimbic areas include parts of the medial, ventral, and orbitofrontal cortical areas adjacent to the corpus callosum, hippocampus and piriform cortex. They connect to limbic areas, to other paralimbic areas, and to circumscribed neocortical areas (Vogt & Pandya, 1987; Morecraft et al., 2004; Morris et al. 1999; Kobayashi & Amaral, 2003; Blatt et al., 2003). In doing so, they bridge regions of the neocortex that represent aspects of the external world and limbic system structures that convey motivational level, homeostatic signals, emotions and visceral inputs (Pandya & Yeterian, 1985; Mesulam, 2000; Pandya et al., 2015).

##### a. Subcentral Association Cortex

This region is associated with Brodmann area 25, and is highly interconnected with medial and posterior orbitofrontal prefrontal areas and subcortical emotional centers (e.g., amygdala) as well as subcortical areas involved in autonomic and visceral function (An et al., 1998; Rempel-Clower & Barbas, 1998; Ongur et al., 1998; Freedman et al., 2000; Chiba et al., 2001; Joyce & Barbas, 2018). It also receives signals from the temporal polar area and medial temporal lobe regions. Thus, this area appears to be a visceromotor region that receives input from prefrontal and temporal lobe areas as well as from the amygdala to relay signals to autonomic effector nuclei in the hypothalamus, brainstem and spinal cord directly (An et al., 1998; Ongur et al., 1999) or indirectly (Freedman et al., 2000).

##### b. The anterior cingulate cortex

The anterior cingulate cortex contains a number of individual areas. The CGa parcellation unit contains Brodmann areas 32 and 24, and is also consistent with the inclusion of both ACC and middle cingulate cortex, as set forth by Vogt and colleagues (2005). The areas within the CGa parcellation unit are connected with limbic regions (e.g., hippocampus) as well as proisocortical areas in the parahippocampal gyri, the posterior cingulate and retrosplenial cortex. There are also connections with somatosensory association areas, premotor areas, and heteromodal areas in the temporal and, to a lesser extent, the inferior parietal regions (Rosene & Van Hoesen, 1977; Petrides & Pandya, 1984; Matelli et al, 1986; Vogt & Pandya, 1987; Luppino et al., 1993; Carmichael & Price, 1995; Saleem et al., 2008; Morecraft et al., 2012). The anterior cingulate has extensive and widespread connections with orbital and lateral prefrontal regions (Pandya et al., 1971; Carmichael & Price, 1996; Gerbella et al., 2010; Petrides & Pandya, 2002; Petrides & Pandya 2007; Morecraft et al., 2012). A variety of functions have been ascribed to the anterior and middle cingulate cortex, including autonomic-visceromotor regulation, production of affective behaviors, nociception, cognitive control, error signal processing, motor control and response selection - differing functions that may be tied to further subdivisions of the anterior cingulate cortex (Devinsky et al., 1995; Paus, 2001).

##### c. The posterior cingulate/retrosplenial cortex

These posterior cingulate and retrosplenial regions have highly similar projections, the main difference being that the retrosplenial cortex has a projection to entorhinal cortex, whereas the posterior cingulate cortex does not (Vogt and Pandya, 1981; Morris et al., 1999; Kobayashi & Amaral, 2003; Kobayashi & Amaral 2007). The retrosplenial cortex maintains connections to other paralimbic areas (parahippocampal cortex), periallocortical and limbic regions (entorhinal cortex, hippocampal formation), and unimodal association and lateral prefrontal areas (Insausti & Amaral, 2008; Van Hoesen & Pandya, 1975; Suzuki & Amaral, 1994; Miyashita & Rockland, 2007; Aggleton et al., 2012; Kobayashi & Amaral, 2003; Kobayashi & Amaral 2007; Passarelli et al., 2018). The functions of the retrosplenial and posterior cingulate cortex, like the anterior cingulate, are debated. By virtue of its connection to both the prefrontal and entorhinal cortex, the retrosplenial cortex may provide linkage between frontal and hippocampal circuits to play a role in spatial memory and navigation, and working memory.

##### d. The parahippocampal cortex

The parahippocampal and adjacent perirhinal cortex contains a number of cortical areas that have strong connections to the heteromodal regions of the parietal, frontal and temporal areas, as well as unimodal inputs from visual and auditory cortices (Van Hoesen, 1982; Blatt et al., 2003). In addition, parahippocampal cortex receives input from other paralimbic areas such as the retrosplenial and cingulate regions, as well as limbic structures, most notably the hippocampus (Suzuki & Amaral, 1994; Lavenex et al., 2002; Blatt et al., 2003). The PHG contains three main architectonic areas, TH, TL and TF (Von Bonin & Bailey, 1947; Blatt et al., 2003), each of which have different patterns of connectivity with unimodal and heteromodal neocortical areas, and with hippocampal areas (Blatt & Rosene, 1998; Blatt et al., 2003).

Parahippocampal areas, in conjunction with adjacent perirhinal and entorhinal regions (Amaral et al., 1987; Suzuki & Amaral, 1994; Lavenex et al., 2002), are involved in processing of space and objects, as well as providing an important linkage with mnemonic limbic system structures. The PHG parcellation unit includes areas TL, TH and TF, as well as perirhinal areas and entorhinal areas.

##### e. The temporal pole

As with all paralimbic structures, the temporal pole has connections with neocortical areas, other paralimbic structures, and limbic regions such as the primary olfactory cortex, the hippocampus and the amygdala (Markowitsch et al., 1985; Moran et al., 1987; Carmichael & Price, 1995). Also similar to other paralimbic regions, the temporal pole is a collection of regions with differing connectivity (e.g., medial temporal polar regions are more heavily connected with olfactory areas, whereas lateral temporopolar areas connect with auditory areas (Moran et al., 1987; Kondo et al., 2003). The temporal pole is also highly connected with the orbitofrontal cortex as well as adjacent regions of the insula and the inferior and superior temporal gyrus (Markowitsch et al., 1985; Kondo et al., 2003). The temporal pole regions have been implicated in visual learning and memory (Horel et al., 1984; Gower, 1989; Bigelow et al., 2016).

#### Insula

The insula contains a number of architectonically discrete areas (Mesulam & Mufson, 1982a,b; Mufson & Mesulam, 1982; Evrard et al., 2014; Morecraft et al., 2015), and has been implicated in a wide number of functions (Augustine, 1996). A large number of unimodal, heteromodal, limbic and paralimbic areas send projections to insular cortex, including visual, olfactory, auditory, somatosensory, gustatory, vestibular, visceral, and motor regions (Mesulam & Mufson, 1982b; Mufson & Mesulam, 1982; Craig, 2014). Rostral insular regions are likely more heavily involved with visceral, gustatory and olfactory processing as well as motor function (Morecraft et al., 2015), given their proximity to associated cortices. By contrast, more caudal insular areas are linked to adjacent somatosensory, vestibular, auditory and visual areas. The wide number of inputs may permit the insula to integrate a broad range of signals from multiple modalities (Schneider et al., 1993; Augustine, 1996; Evrard et al., 2014). The anterior insula also contains von Economo neurons (Evrard, 2012), which are thought to play a role in social cue processing and autonomic regulation (Allman et al., 2010; Allman et al. 2011).

### Ontology

The organization of cerebral cortical regions in the non-human primate has been delineated on numerous occasions, and the pattern of organization has reflected the methods and knowledge at the time each schema was formulated (e.g., Brodmann, 1909; von Bonin & Baily, 1947; Braak, 1980). As technological capabilities improved, information from physiological, neuroanatomical, and immunohistochemical studies provided a higher level of detail of brain mapping (e.g., Paxinos et al,. 2009; Saleem et al., 2012). This detail has often led to further subdivisions of areas, but the resulting alterations in nomenclature and taxonomy have not always been brought into alignment or unified across the individual systems. What has been needed is a formal means by which nomenclature, origin, meaning and inter-relationship between structures (i.e., ontology) is systematized. This type of approach would thereby begin to create a comprehensive framework for description at several scales of analysis, including inter-species comparisons (Nichols, 2014; Bowden & Dubach, 2003; Imam et al., 2012; ten Donkelaar et al., 2017). Such comparisons have been performed in the monkey through the NeuroNames / NeuroMaps initiative (e.g., Bowden & Dubach, 2003; Bowden et al., 2007; Rohling et al., 2012; Bowden et al., 2012). To that end, we incorporated a taxonomic structure for the rhesus monkey cerebral cortex consistent with the NeuroNames system, as well as other systems developed to provide ontological frameworks for human neuroanatomical structures (e.g., ten Donkelaar et al., 2017, 2018; Swanson, 2015). It should be noted that the use of a standardized terminology required employing alternative terms for some that are more commonly used (e.g., marginal sulcus vs. marginal ramus of the cingulate sulcus). These alternate forms are collated for reference in Table 5. The present tabulation provides the beginning of a description that has importance for the organization of the proposed parcellation framework. It should be emphasized that the mHOA proposed ontology is designed to preserve homological correspondences between structures in the monkey and human brains as described in the HOA. Thus, it provides a means to comparatively evaluate the structures from the two species.

### Future Studies

The present system serves as the first step in systematically organizing, parcellating and contextualizing the rhesus monkey cerebral cortex according to the HOA framework, namely a means by which individual macaque and human brains may be parcellated according to homological principles. In the future, this system should extend to comprehensively include limbic system structures, deep telencephalic structures as well as diencephalic and brainstem regions. As herein, the approach should be to provide clear and reproducible anatomical borders. This system will also be extended to include more detail in the cortex and other brain regions as technology becomes increasingly able to reliably distinguish finer-grained areas on an individual basis. More extensive parcellation systems based on a higher number of anatomical divisions can and have been described (Martin & Bowden, 1996; Black et al., 2004; McLaren et al., 2009; Quallo et al., 2010; Paxinos et al., 2008; Frey et al., 2011; Van Essen et al., 2012; Saleem & Logothetis, 2012; Calabrese et al., 2015; Feng et al., 2017; Reveley et al., 2017). However, these approaches use divisions and borders that are not directly visible in MRI images. It is a subject of current debate to what extent we can make fine-grained and accurate MRI-based parcellations to fit atlases based on histologically prepared post-mortem material or a common stereotaxic atlas (e.g., Eickhoff et al., 2007). The advantage of such approaches is that they allow for a high resolution of anatomical detail. However, the main disadvantage in the use of atlas-based coordinate systems is that these systems are limited in accounting for variability in individual brain structure and morphometric measures derived from such atlases might be less sensitive to subtle structural pathology. There have been some measures of variability of gross anatomical structures in rhesus monkey brains (van der Gucht et al., 2006; Pereira-Pedro et al., 2017; Croxson et al., 2018), however, variability in the size and position of architectonic regions in the rhesus monkey has not yet been systematically or comprehensively studied. Intersubject variability or laterality has also not been well studied in rhesus macaques, but the studies that do exist as well as similar studies in humans have shown substantial intersubject variability in terms of size and shape, and in relationship to extant anatomically-defined borders (Van Essen et al., 1984; Maunsell & Van Essen, 1987; Geyer et al,. 1999; Amunts et al., 2000; Croxson et al., 2018). These and other studies also show that the location and size of higher order areas with respect to sulci and gyri are much more variable than primary or secondary regions (Raijkowska & Goldman-Rakic, 1995a,b; Amunts et al., 1999; Amunts et al., 2000; Uylings et al., 2005; Fischl et al., 2008; Hinds et al,. 2008; Croxson et al., 2018). This variability is not captured by techniques that fit individual brains to atlas or templates. The advantage of the mHOA system, as with the human HOA, is that it parcellates each brain based on its individual anatomy and therefore preserves variability. Previous studies have shown that the HOA approach to human brain volumetry is effective in capturing individual brain structure (Rademacher et al., 1992; Caviness et al., 1996; Makris et al., 2003, 2005). Unlike automated systems, the present approach entails manual and semi-automated procedures as was done for the human HOA (Rademacher et al., 1992; Caviness et al., 1996; Makris et al., 2003, 2005). It should be noted these manual systems provided the gold standard for validation of the currently available automated volumetry by FreeSurfer (Fischl et al., 2002, 2004). In the absence of an automated system to generate a monkey HOA, the present approach establishes a basis in this regard. This approach has been originated by our group in developing the field of MRI-based human brain ‘volumetrics’ (Caviness et al., 1999). Future monkey studies in our laboratory will use inter- and intra-rater reliability to refine the mHOA system and facilitate the creation of anatomically accurate automated methods.

As resolution increases and multimodal imaging technology extends the capacity to make strong linkages between signal and underlying anatomy, we fully anticipate that the borders between areas as defined by physiological properties, immunohistochemical distinctions, anatomical properties or connectivity differences will be more robustly and rigorously validated in MRI scans (Augustinack et al., 2005; Glasser et al., 2016). Presently, we have chosen to be conservative in our borders and use consistent limiting sulci and coronal limiting planes as a first step, which has shown to be an effective approach in current MRI-based morphometric analysis of the human brain, such as the HOA (Rademacher et al., 1992; Rademacher et al., 1993; Fischl et al., 2008).

## Conclusions

In this paper, we have developed a neuroanatomical system to parcellate the monkey cerebral cortex using MRI, namely the monkey HOA (mHOA). The procedure employs anatomical features of the individual macaque brain identifiable by MRI to delineate boundaries, and thereby provides information about the neuroanatomy of the specific experimental animal. A similar approach carried out in the human brain led to the creation of the HOA. By adopting a parallel approach in the context of a cross-species ontological framework, the present paper complements the existing human HOA. We anticipate that the mHOA will enable future studies to achieve a more complete definition, technically and conceptually, of the monkey brain circuit diagram (BCD) using MRI. Given that the human BCD relies on homology and thus is comparative in nature, neuroanatomical knowledge of the monkey brain will enhance the completeness of the BCD in the human. Importantly, this system is rule-based in nature and therefore can be adapted as imaging technological capabilities increase or as neuroanatomical conventions change and knowledge is accrued.

## Supporting information

Supplemental GS Images

Supplemental CP images

## Abbreviations

CGa: anterior cingulate
CGp: posterior cingulate
CP: coronal plane
CO: coronal plane
COa: central operculum - anterior
COp: central operculum - posterior
F1dl: dorso-lateral superior frontal
F1dm: dorso-medial superior frontal
F2: inferior frontal
FOC: fronto-orbital
FP: frontal pole
HP: horizontal plane
INS: insula
ITG: inferior temporal gyrus
LPCi: inferior portion of lateral parietal cortex
LPCs: superior portion of lateral parietal cortex
MPC: medial parietal cortex
PO: parietal operculum
PoG: postcentral gyrus
PRL: prelunate
PHG: parahippocampal gyrus
PrG: precentral gyrus
SC: subcallosal area
STG: superior temporal gyrus
STP: supratemporal plane
STRdl: dorsolateral portion of striate cortex
STRm: medial portion of striate cortex
TP: temporal pole
VMO: ventromedial occipital

## Acknowledgements

The authors acknowledge support from the National Institutes of Health (R01 MH112748).

## Supplementary Information

1. Supplementary Table 1: Volumes of General Segmentation Parcellations
2. Cortical Parcellation (CP) Results – Images for the cortical parcellations for each coronal section. Each coronal section is shown in native view (no overlay), with parcellation border shown as overlaid lines, and then as overlaid filled objects. Abbreviations are in margin, and correspond to those in Figure 5.
3. General Segmentation (GS) Results – Images of the general segmentation. Image conventions as in CP Results

