## Supplemental Table 1 for "MRI-based Parcellation and Morphometry of the Individual Rhesus Monkey Brain: a translational system referencing a standardized ontology"

Supplementary Table: General Segmentation Structure Volumes (mm^3^)

|  | **Left** | **Right** |
| --- | --- | --- |
| ***Paired Structures*** |  |  |
| Amygdala | 317.1 | 324 |
| Anterior Amygdala | 15.7 | 11.8 |
| Basal Forebrain | 216.9 | 173 |
| Caudate | 483.5 | 485 |
| Caudate/Putamen | 39.7 | 41.8 |
| Cerebellar Cortex | 3437.9 | 3530.2 |
| Cerebellar White Matter | 1184 | 1128.5 |
| Claustrum | 175.8 | 197.1 |
| Cortical Ribbon | 22540.2 | 22049.3 |
| Globus Pallidus | 346.2 | 361 |
| Hippocampus | 565.9 | 536.3 |
| Hypothalamus | 145 | 124.8 |
| Lateral Hypothalamus | 65.4 | 57.9 |
| Lateral Ventricle | 443.2 | 420.2 |
| Lateral Ventricle - Inf Horn | 63.5 | 60.6 |
| Nucleus Accumbens | 123.4 | 115.5 |
| Putamen | 903.7 | 857 |
| Thalamus | 765.4 | 746.2 |
| Ventral Diencephalon | 557.6 | 540.5 |
| ***Unpaired Structures*** |  |  |
| 3rd-Ventricle | 76.2 |  |
| 4th-Ventricle | 189.4 |  |
| Brainstem | 3938.6 |  |
| Optic-Chiasm | 100.8 |  |
