## Supplemental GS Images for "MRI-based Parcellation and Morphometry of the Individual Rhesus Monkey Brain: a translational system referencing a standardized ontology"

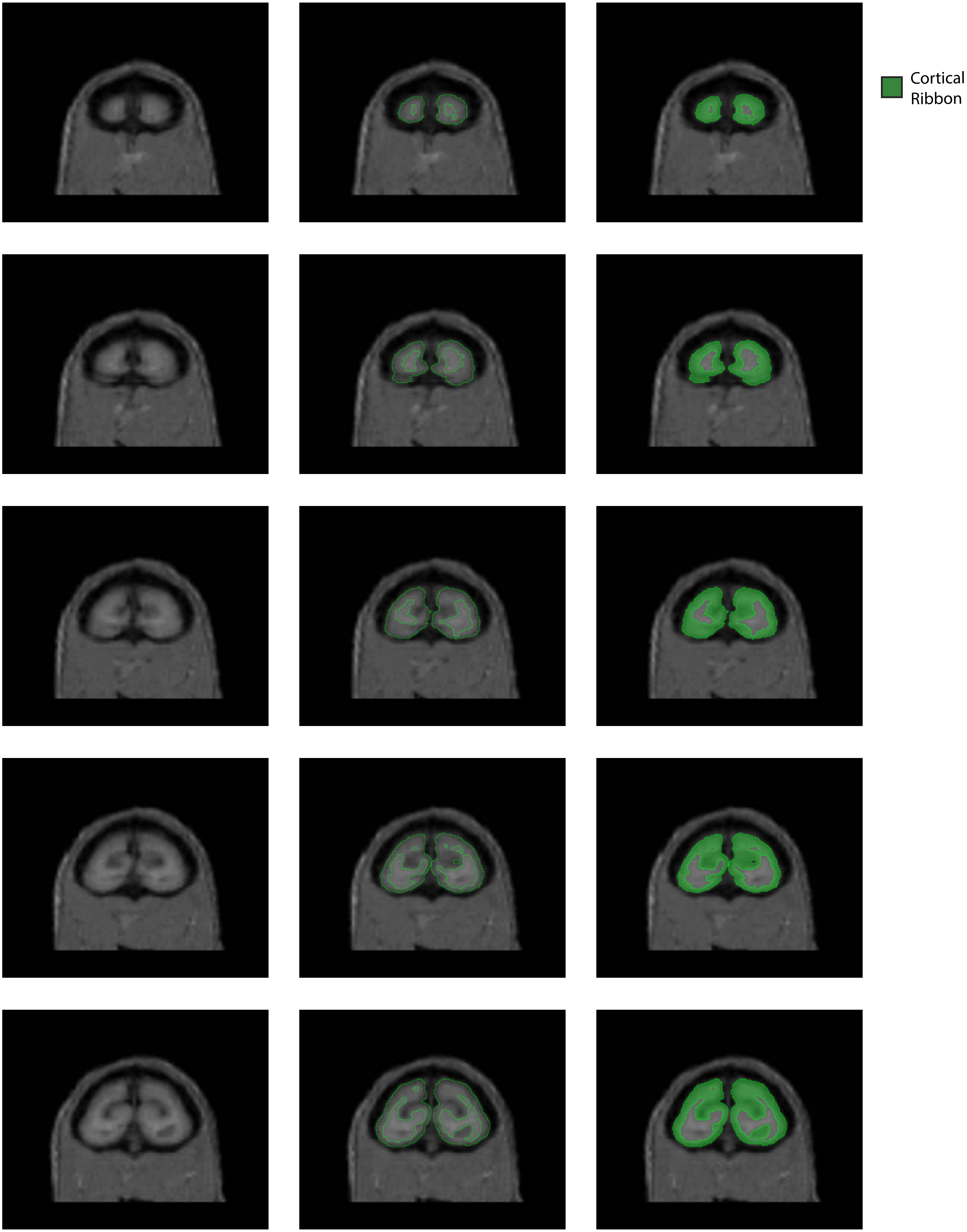
Cortical Ribbon

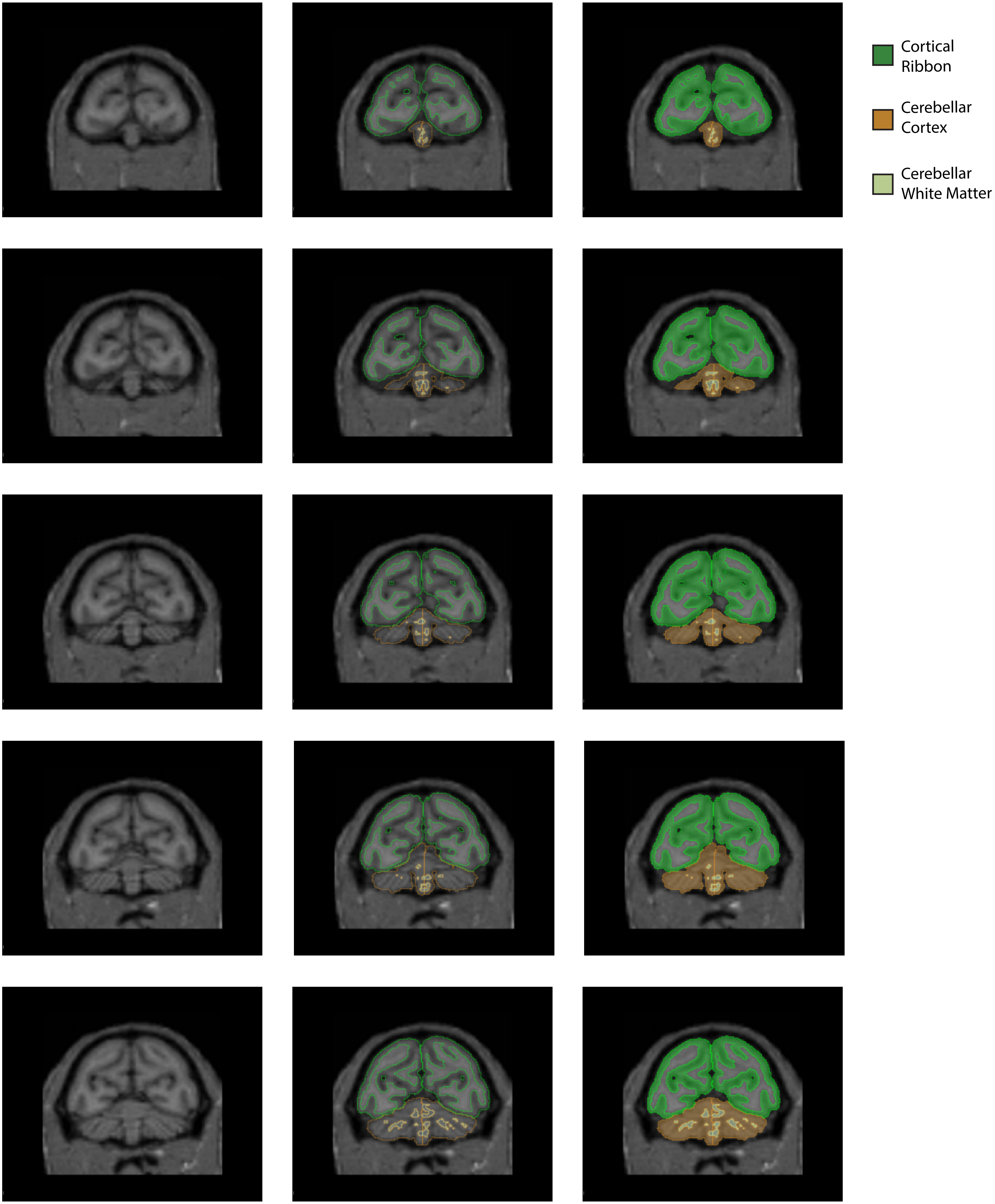
Cortical Ribbon
Cerebellar Cortex
Cerebellar White Matter

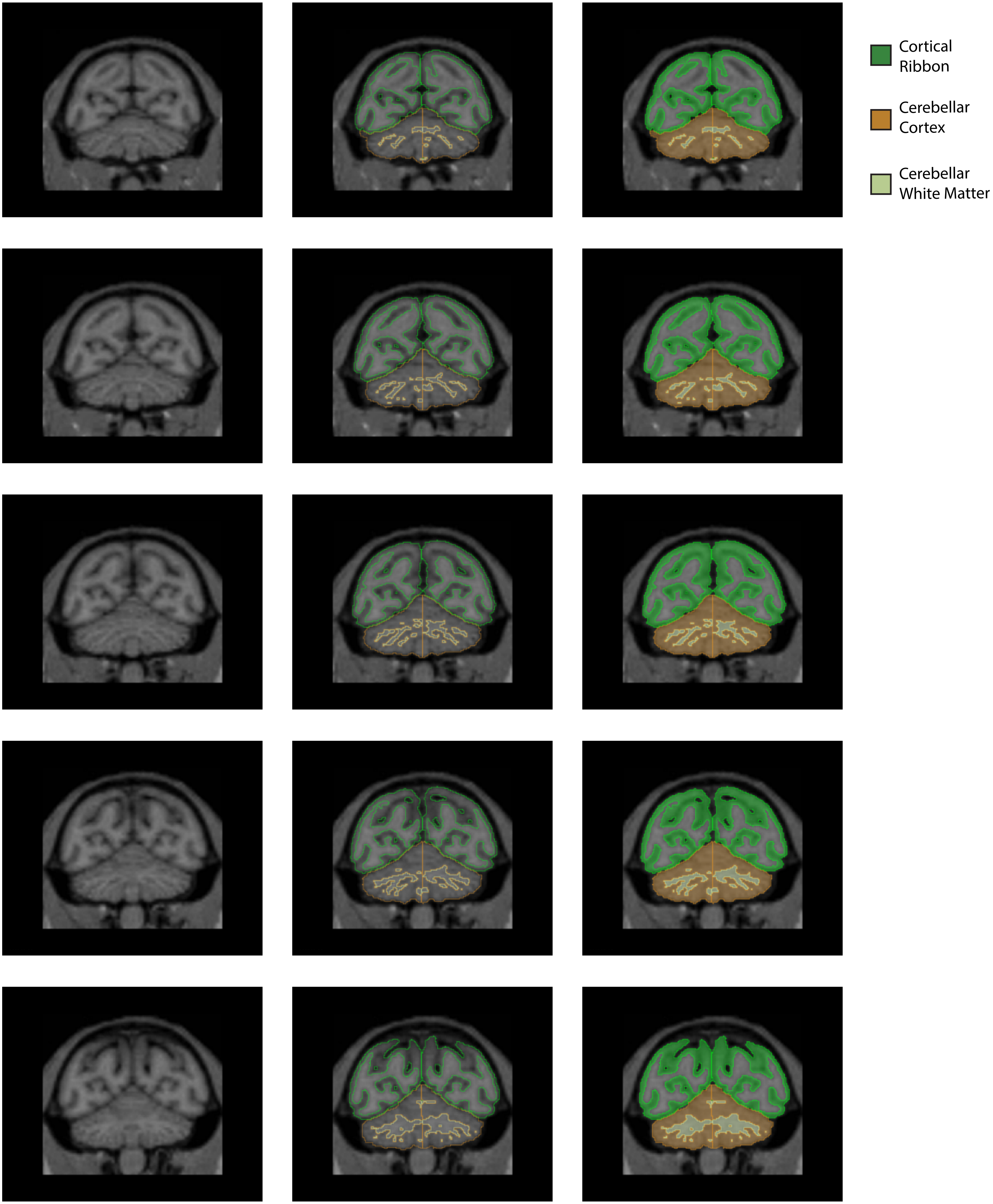

Cortical Ribbon
Cerebellar Cortex
Cerebellar White Matter

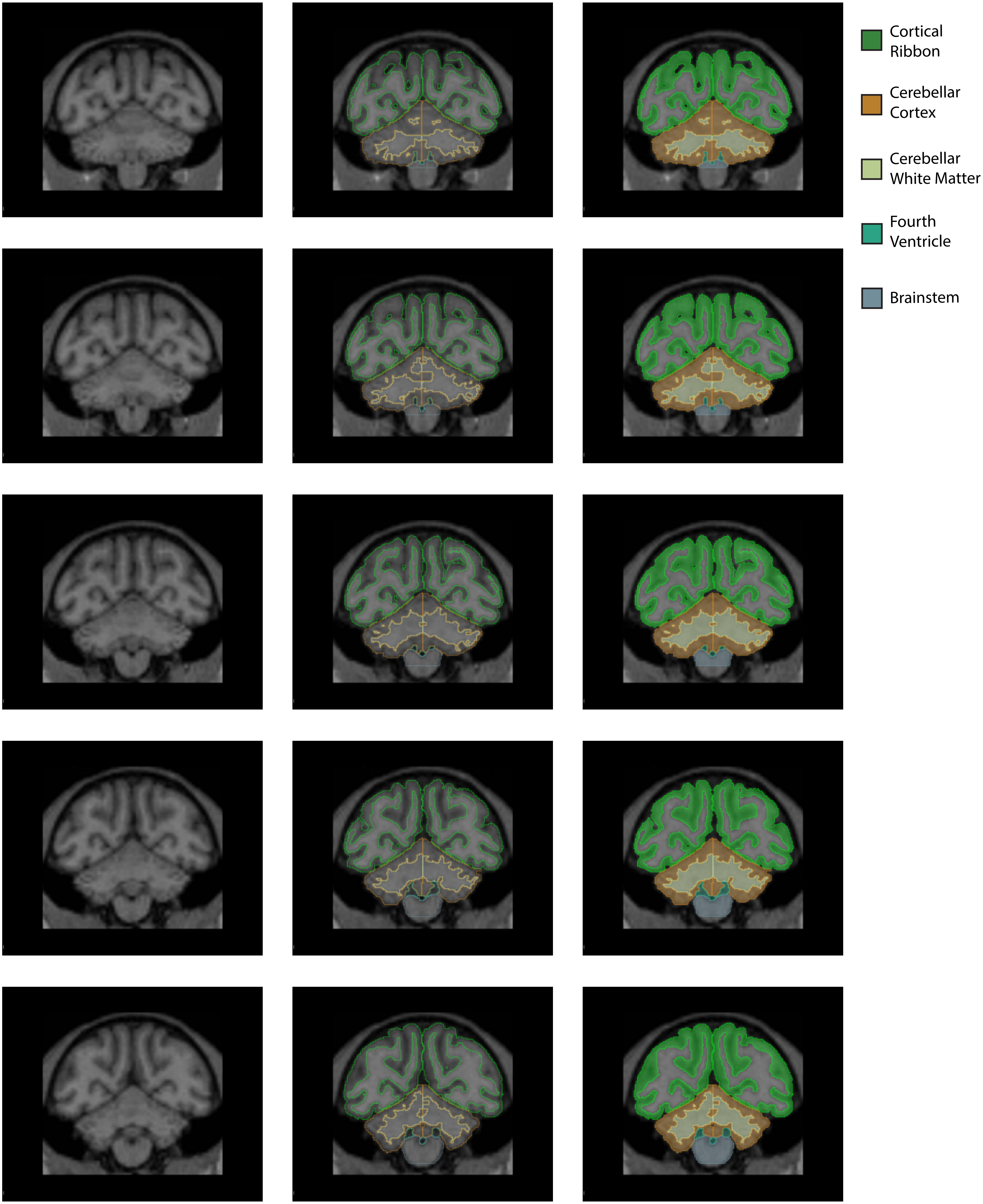

Cortical Ribbon
Cerebellar Cortex
Cerebellar White Matter
Fourth Ventricle
Brainstem

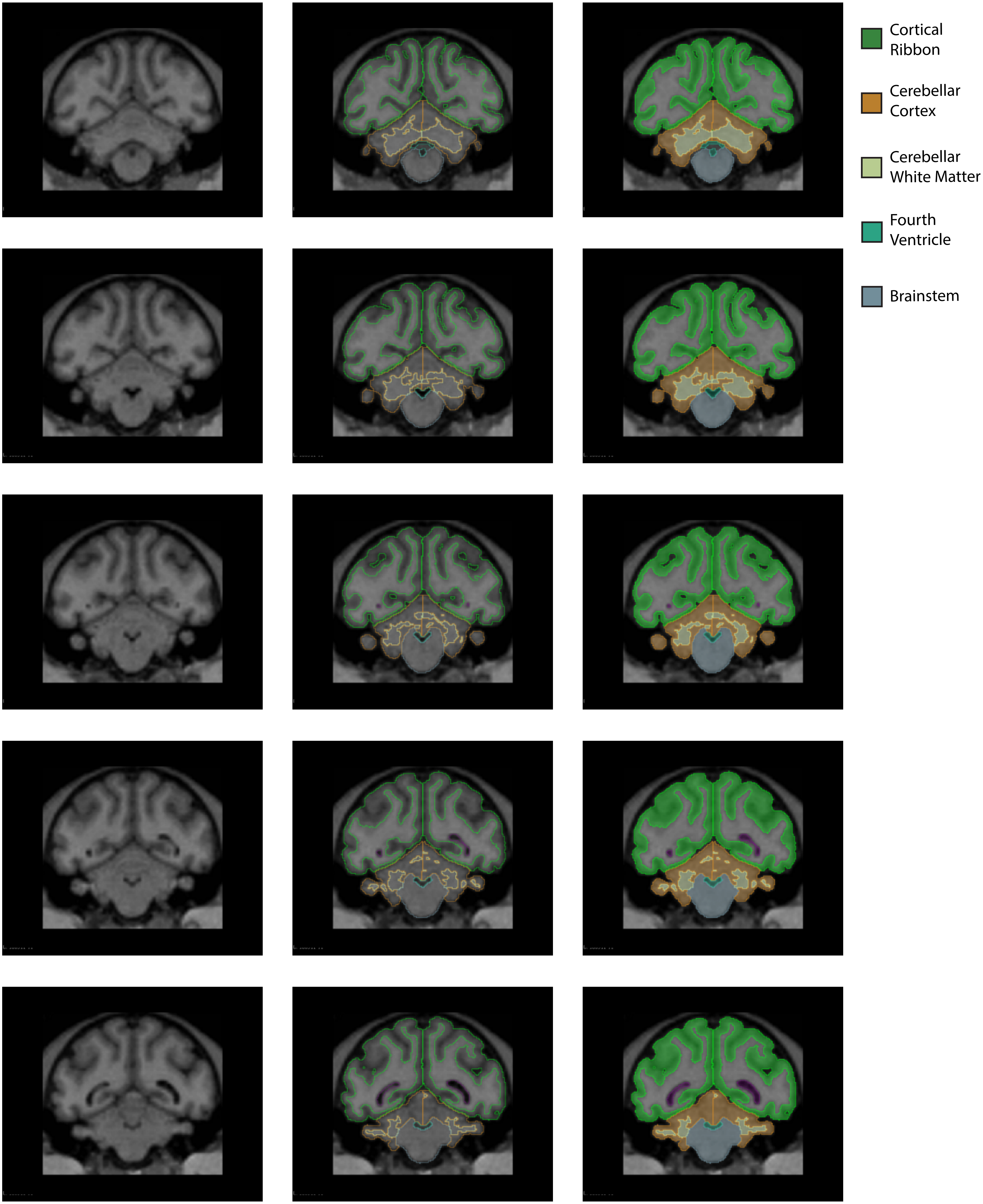
Cortical Ribbon
Cerebellar Cortex
Cerebellar White Matter
Fourth Ventricle
Brainstem

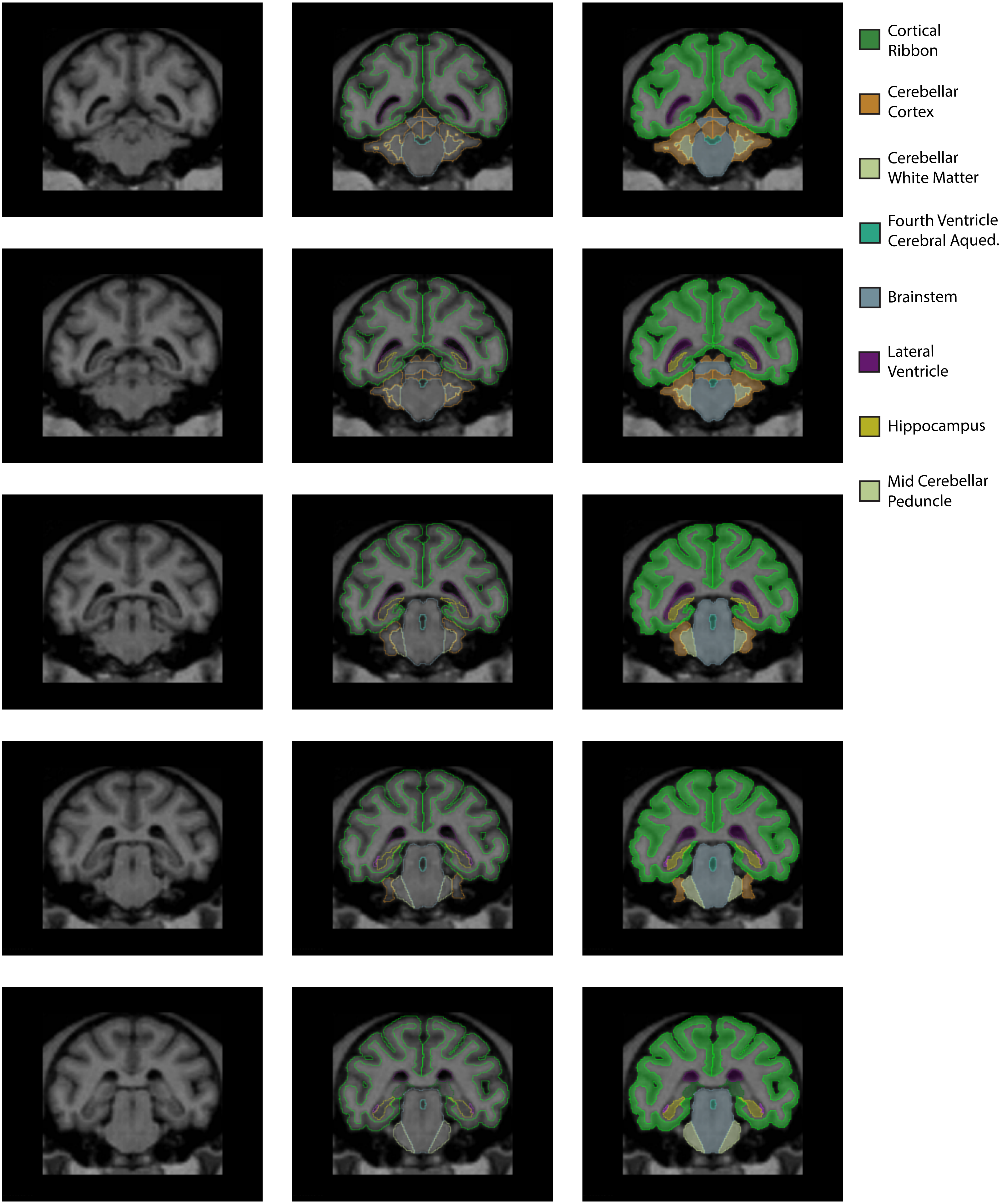
Cortical Ribbon
Cerebellar Cortex
Cerebellar White Matter
Fourth Ventricle Cerebral Aqued.
Brainstem
Lateral Ventricle
Hippocampus
Mid Cerebellar Peduncle

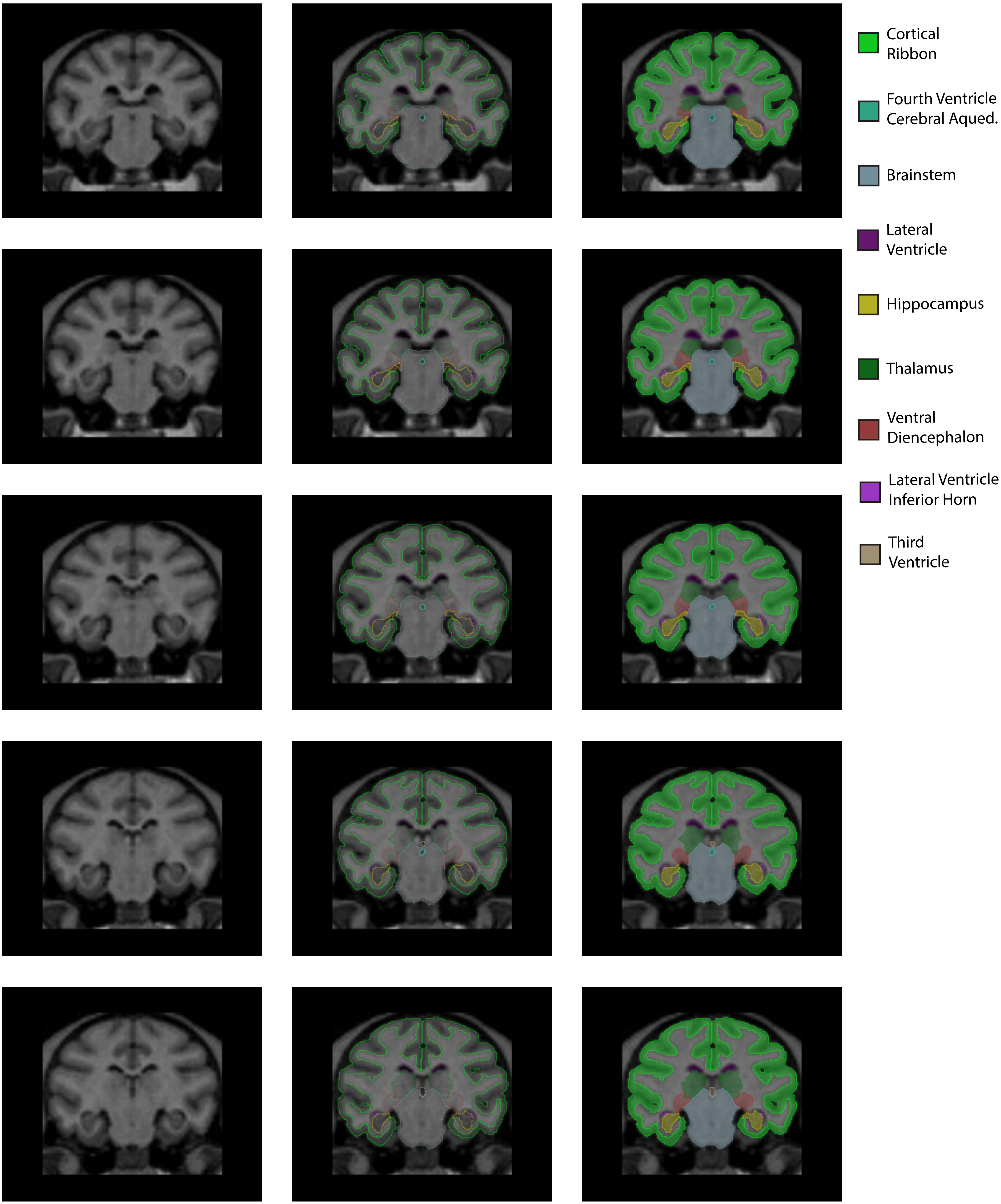

Cortical Ribbon
Fourth Ventricle Cerebral Aqued.
Brainstem
Lateral Ventricle
Hippocampus
Thalamus
Ventral Diencephalon
Lateral Ventricle Inferior Horn
Third Ventricle

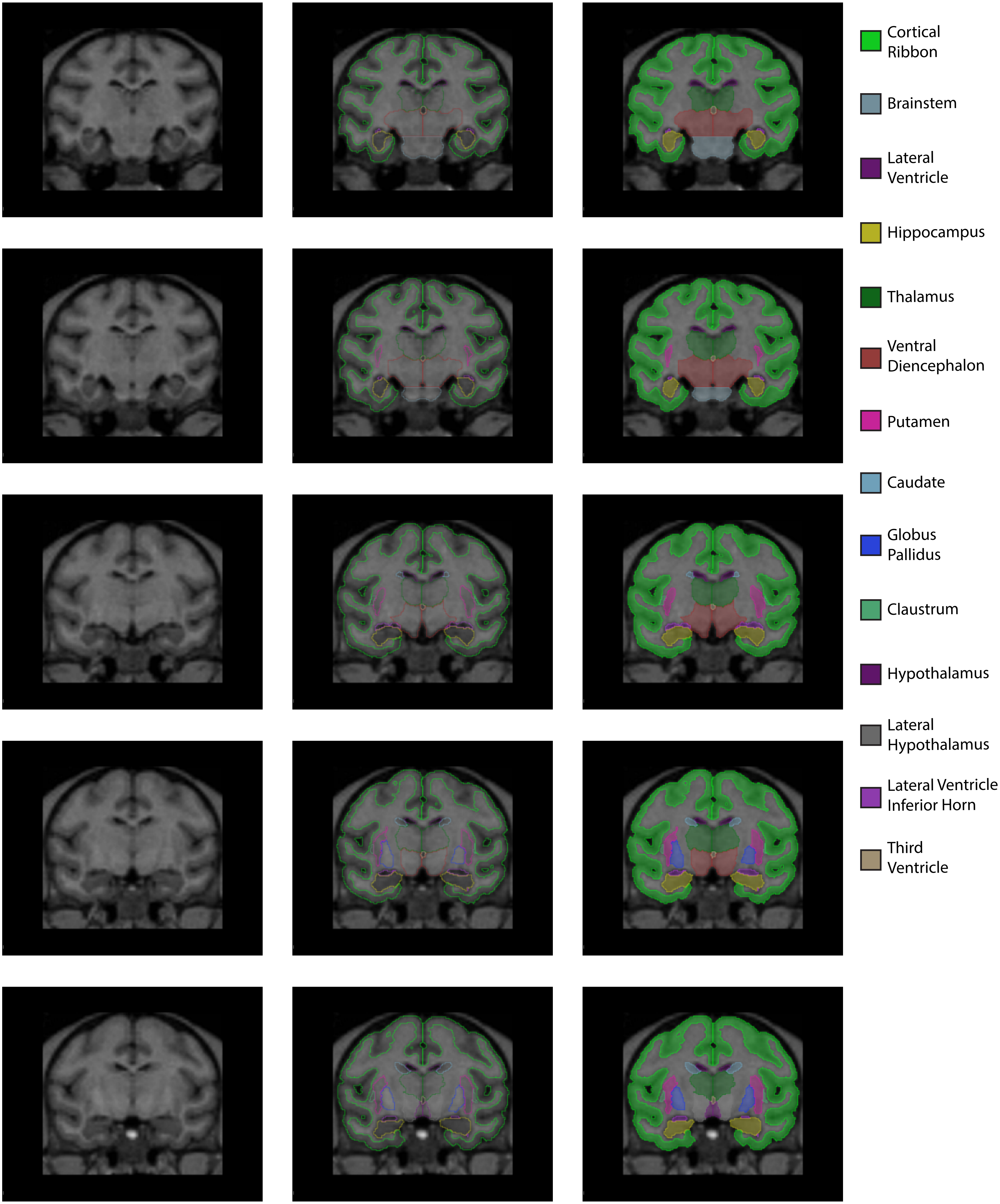
Cortical Ribbon
Brainstem
Lateral Ventricle
Hippocampus
Thalamus
Ventral Diencephalon
Putamen
Caudate
Globus Pallidus
Claustrum
Hypothalamus
Lateral Hypothalamus
Lateral Ventricle Inferior Horn
Third Ventricle

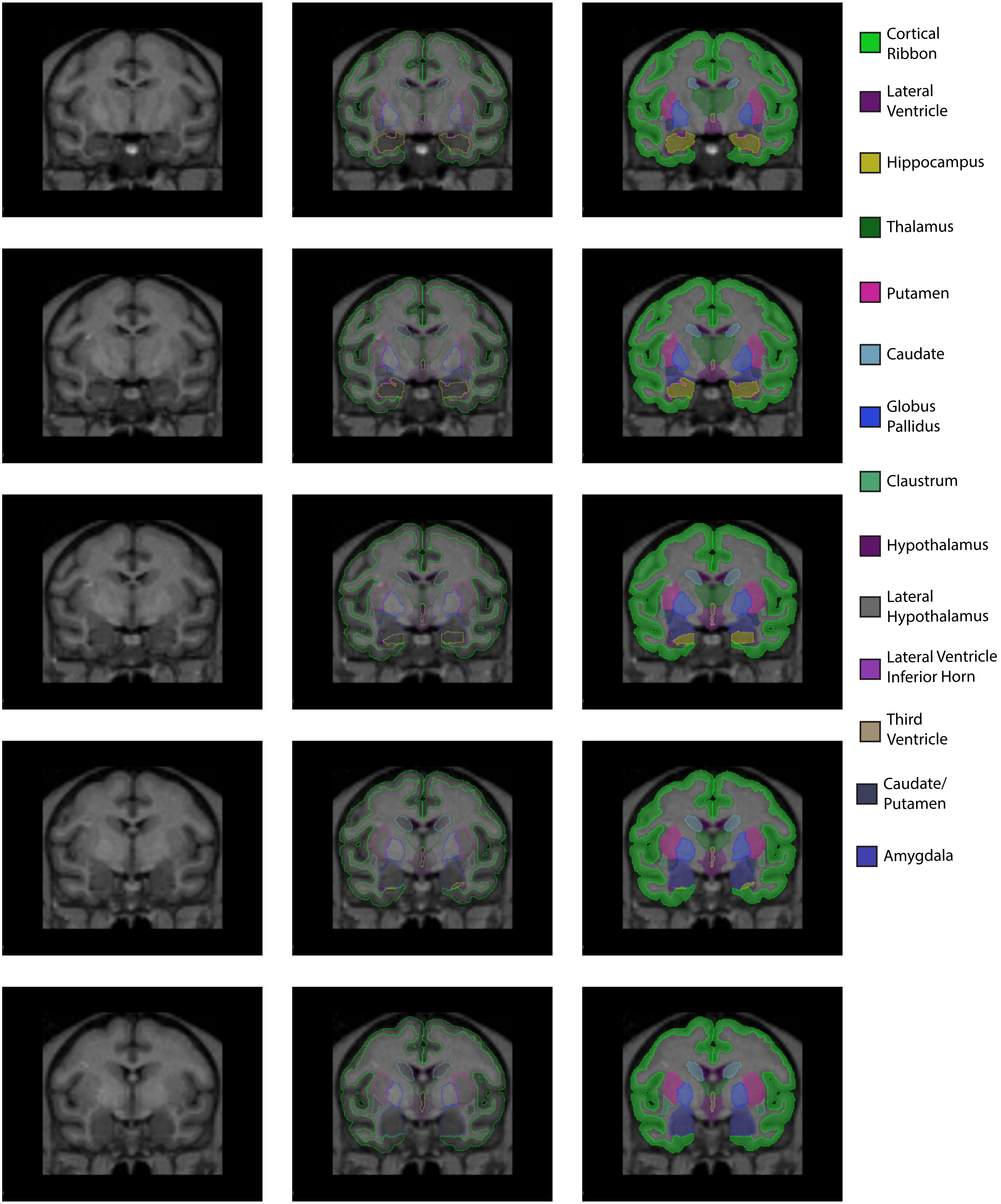
Cortical Ribbon
Lateral Ventricle
Hippocampus
Thalamus
Putamen
Caudate
Globus Pallidus
Claustrum
Hypothalamus
Lateral Hypothalamus
Lateral Ventricle Inferior Horn
Third Ventricle
Caudate/ Putamen
Amygdala

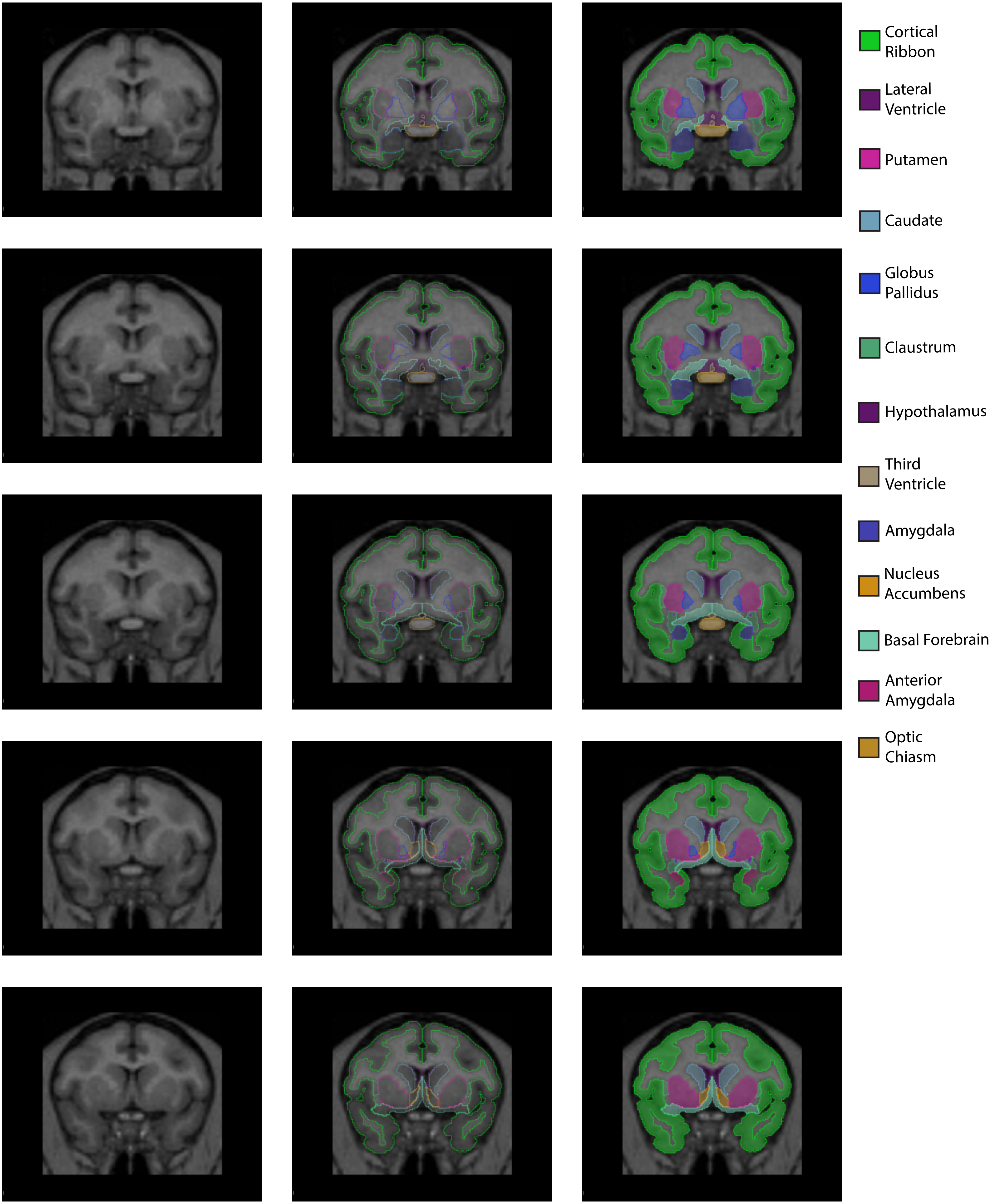
Cortical Ribbon
Lateral Ventricle
Putamen
Caudate
Globus Pallidus
Claustrum
Hypothalamus
Third Ventricle
Amygdala
Nucleus Accumbens
Basal Forebrain
Anterior Amygdala
Optic Chiasm

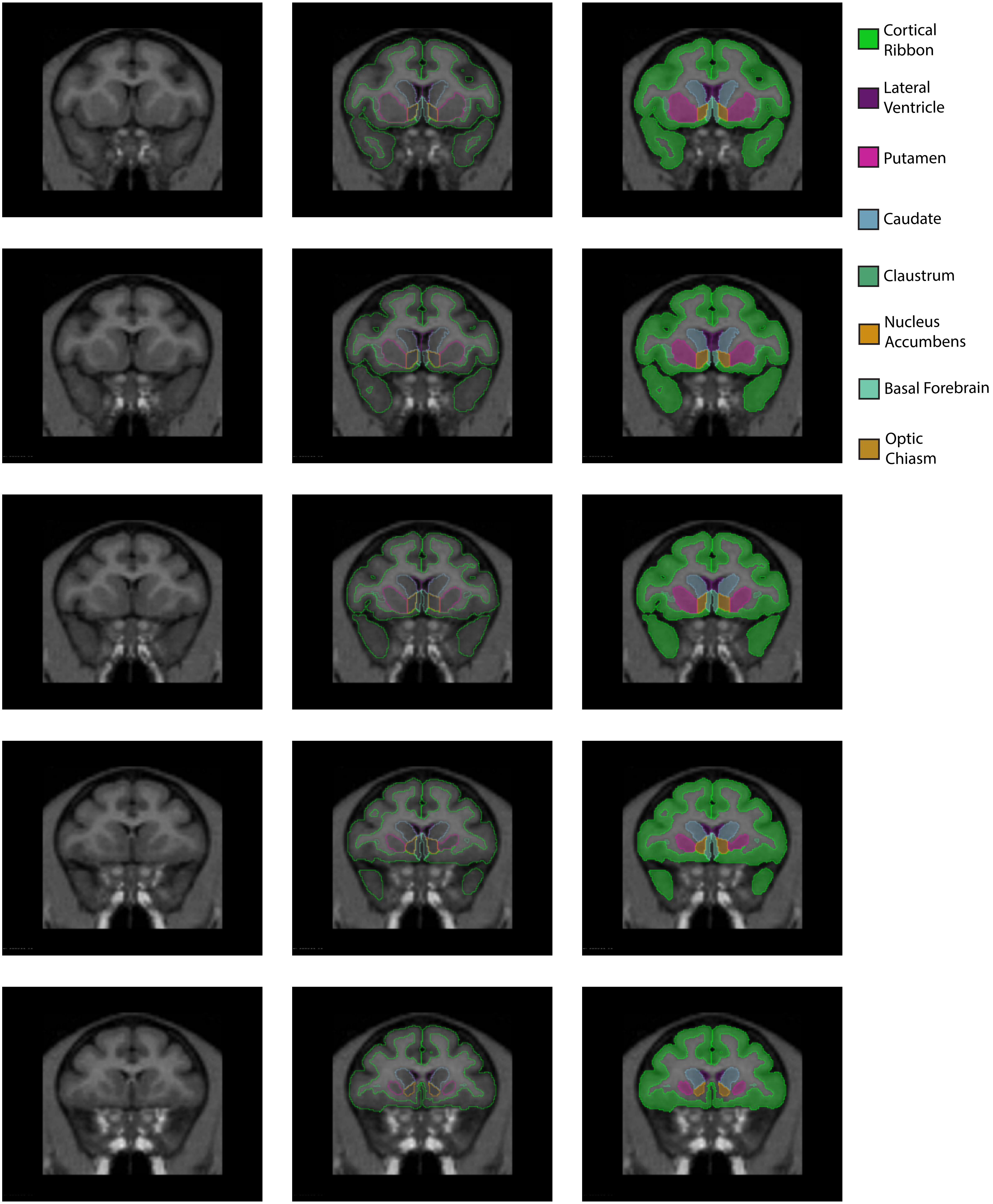
Cortical Ribbon
Lateral Ventricle
Putamen
Caudate
Claustrum
Nucleus Accumbens
Basal Forebrain
Optic Chiasm

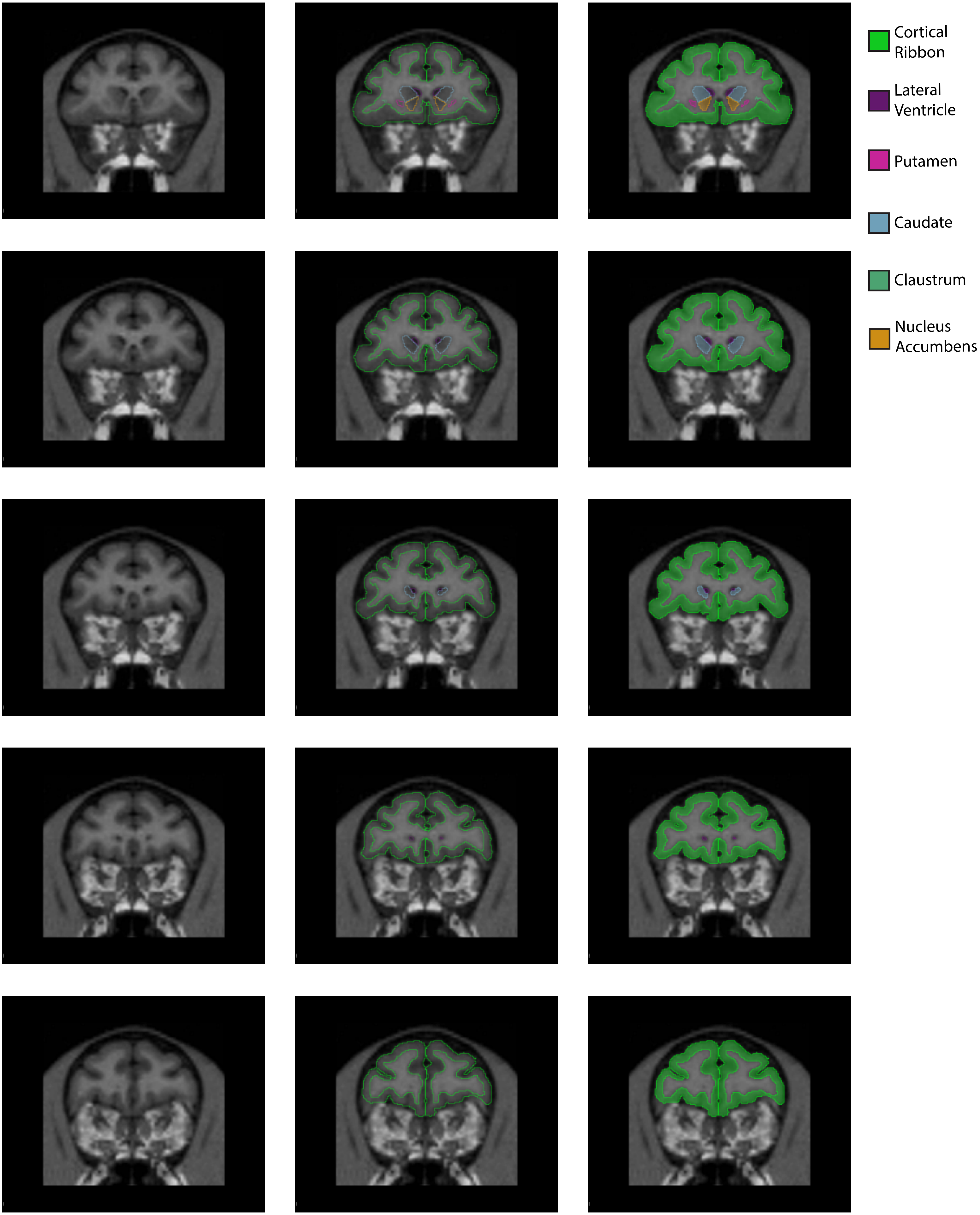
Cortical Ribbon
Lateral Ventricle
Putamen
Caudate
Claustrum
Nucleus Accumbens

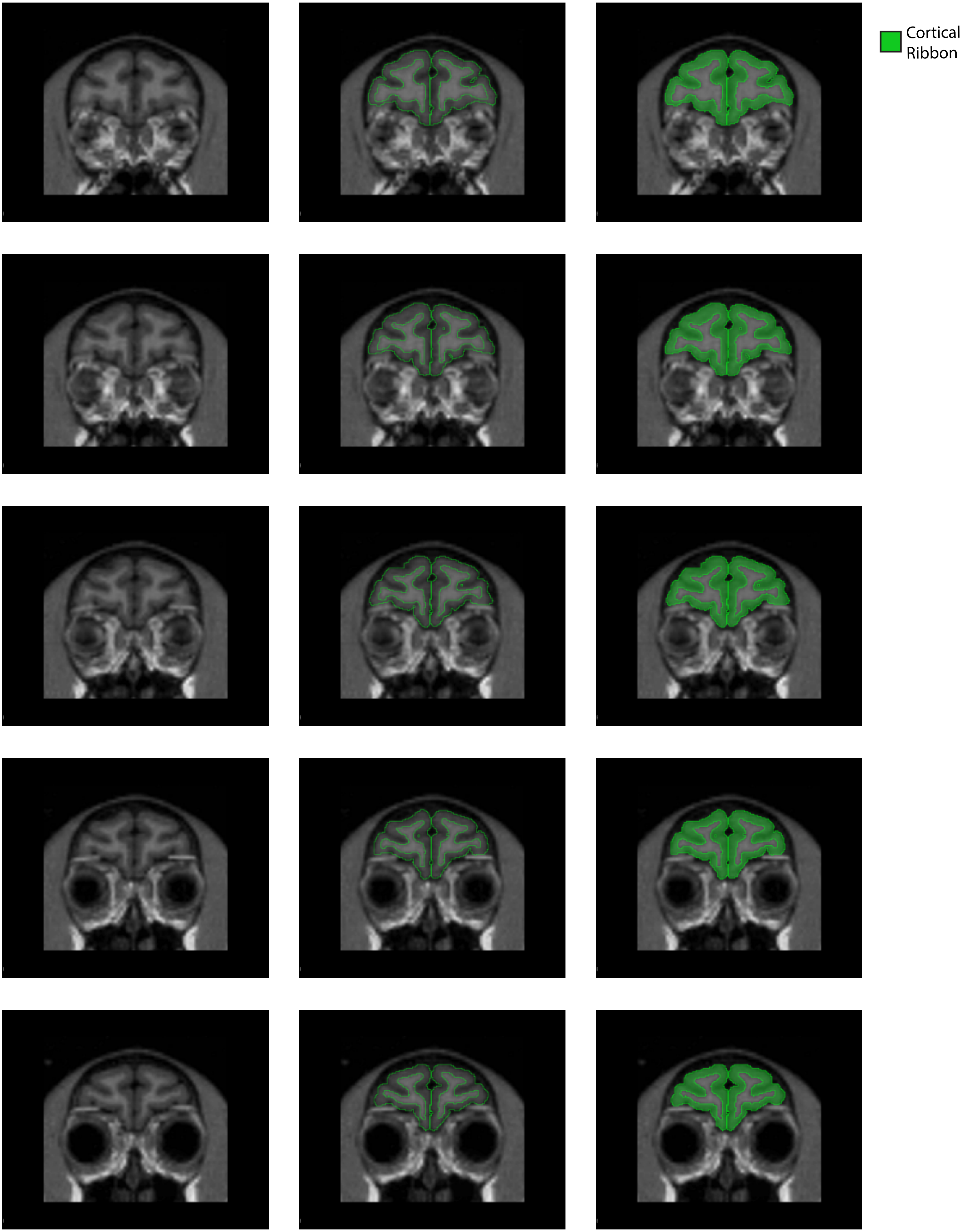
Cortical Ribbon

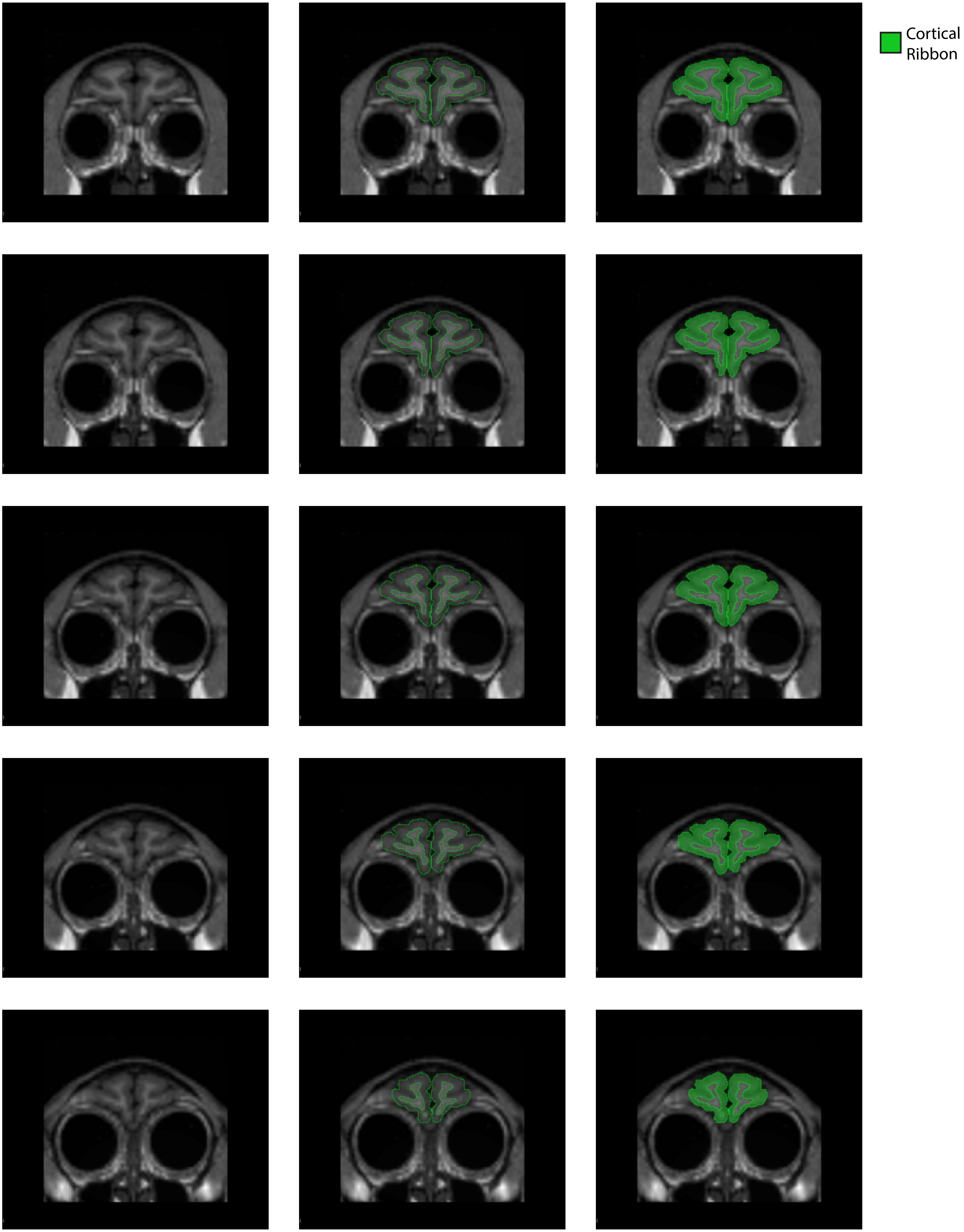
Cortical
Ribbon

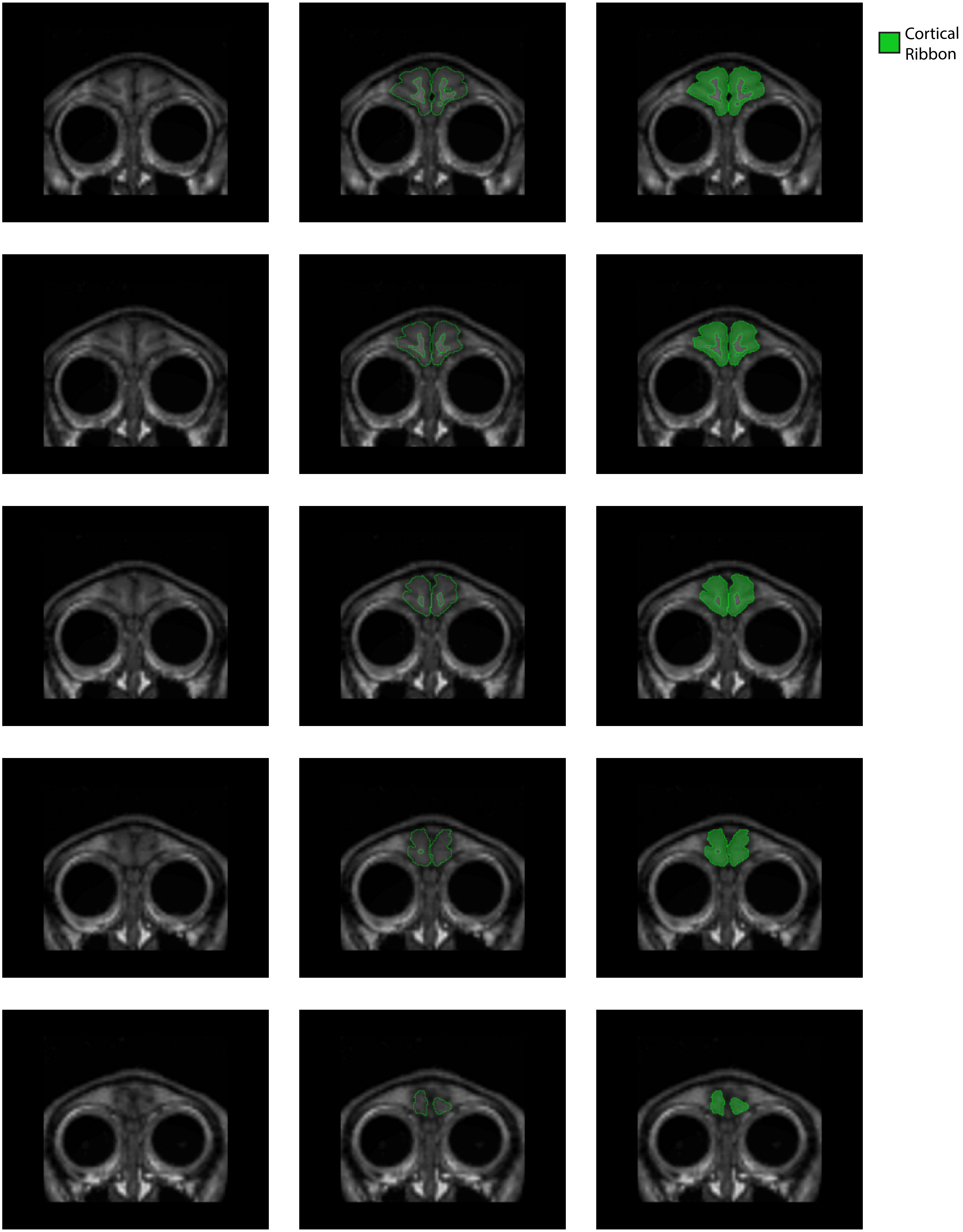
Cortical
Ribbon
