## Supplemental CP images for "MRI-based Parcellation and Morphometry of the Individual Rhesus Monkey Brain: a translational system referencing a standardized ontology"

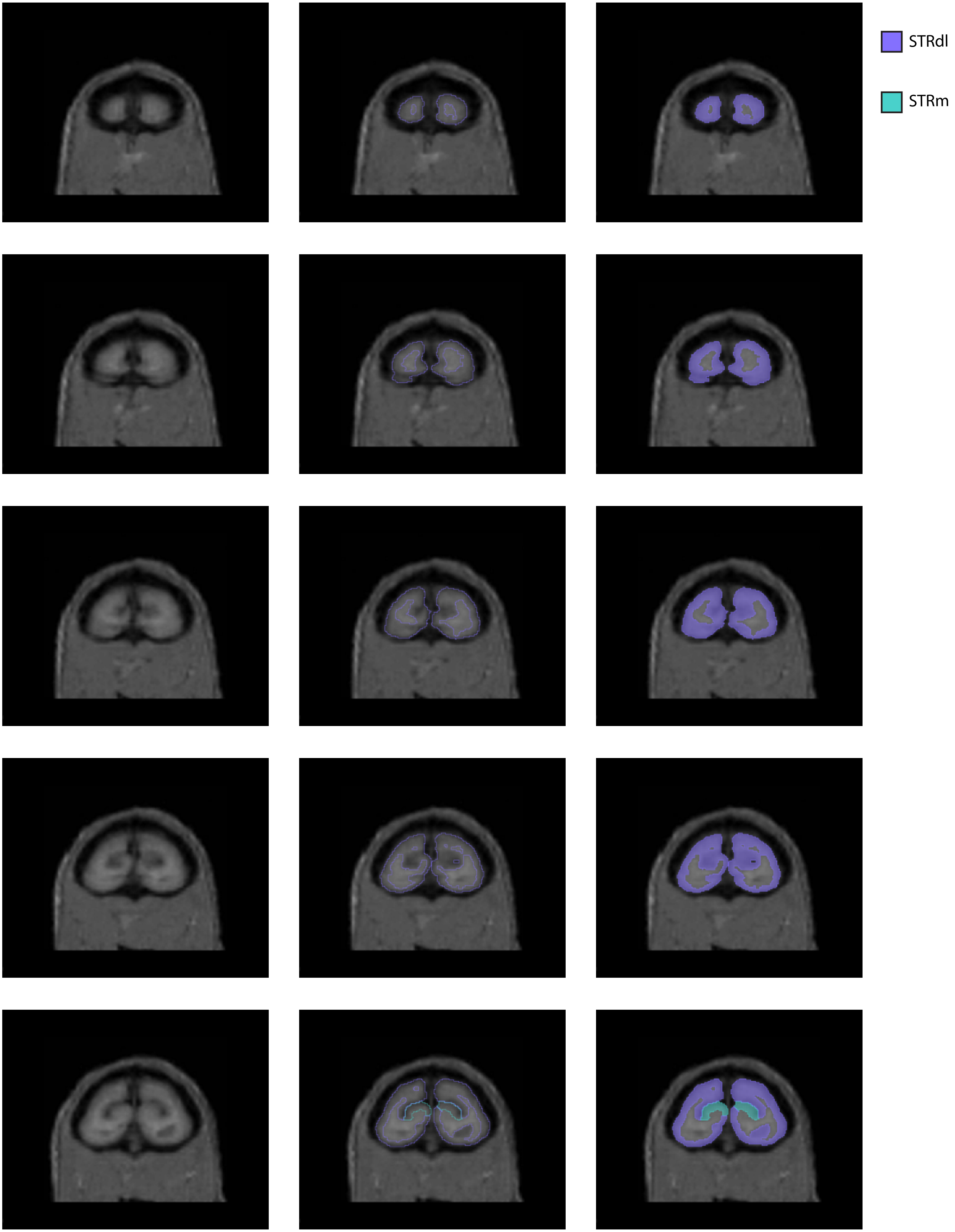
STRdl
STRm

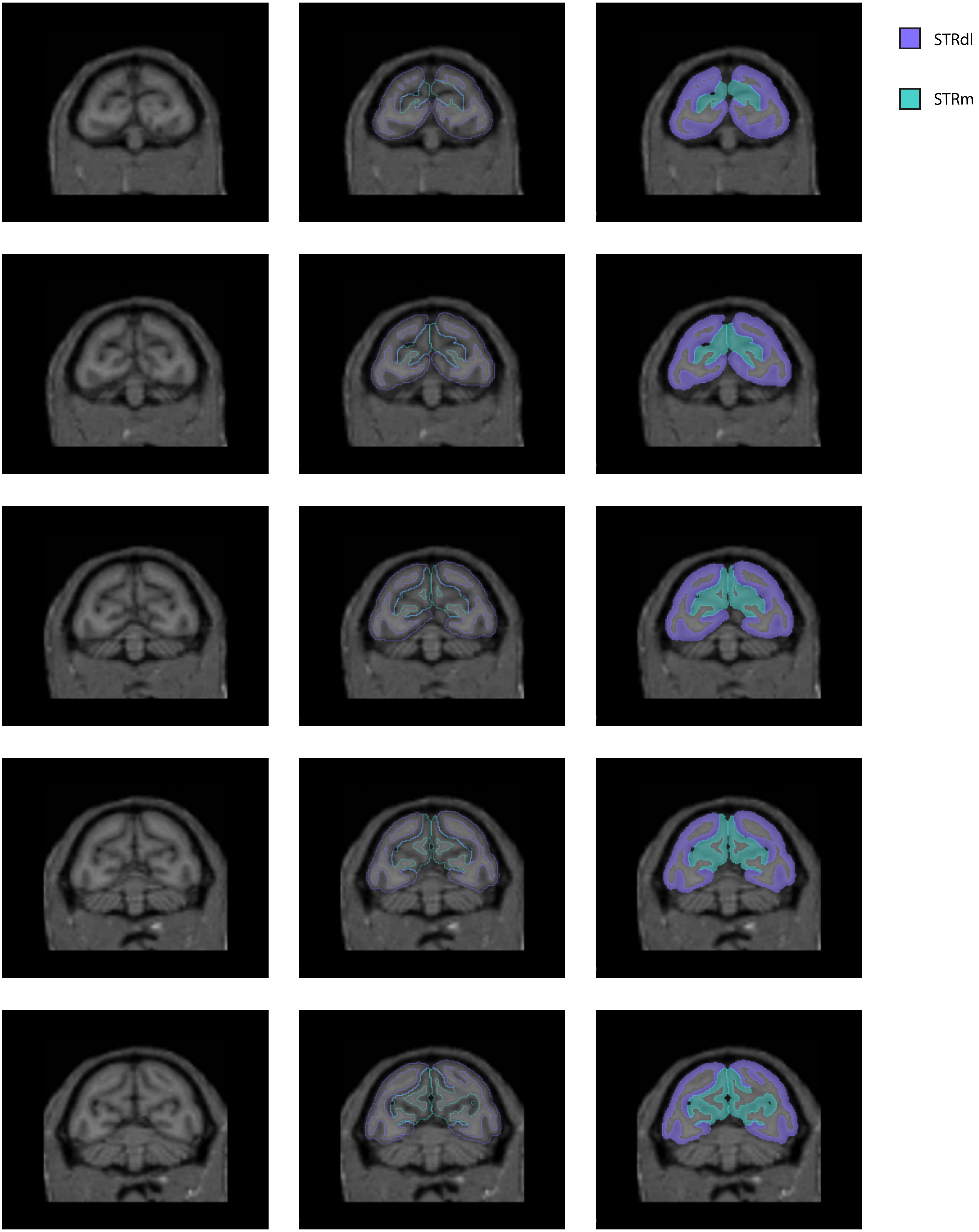

STRdl
STRm

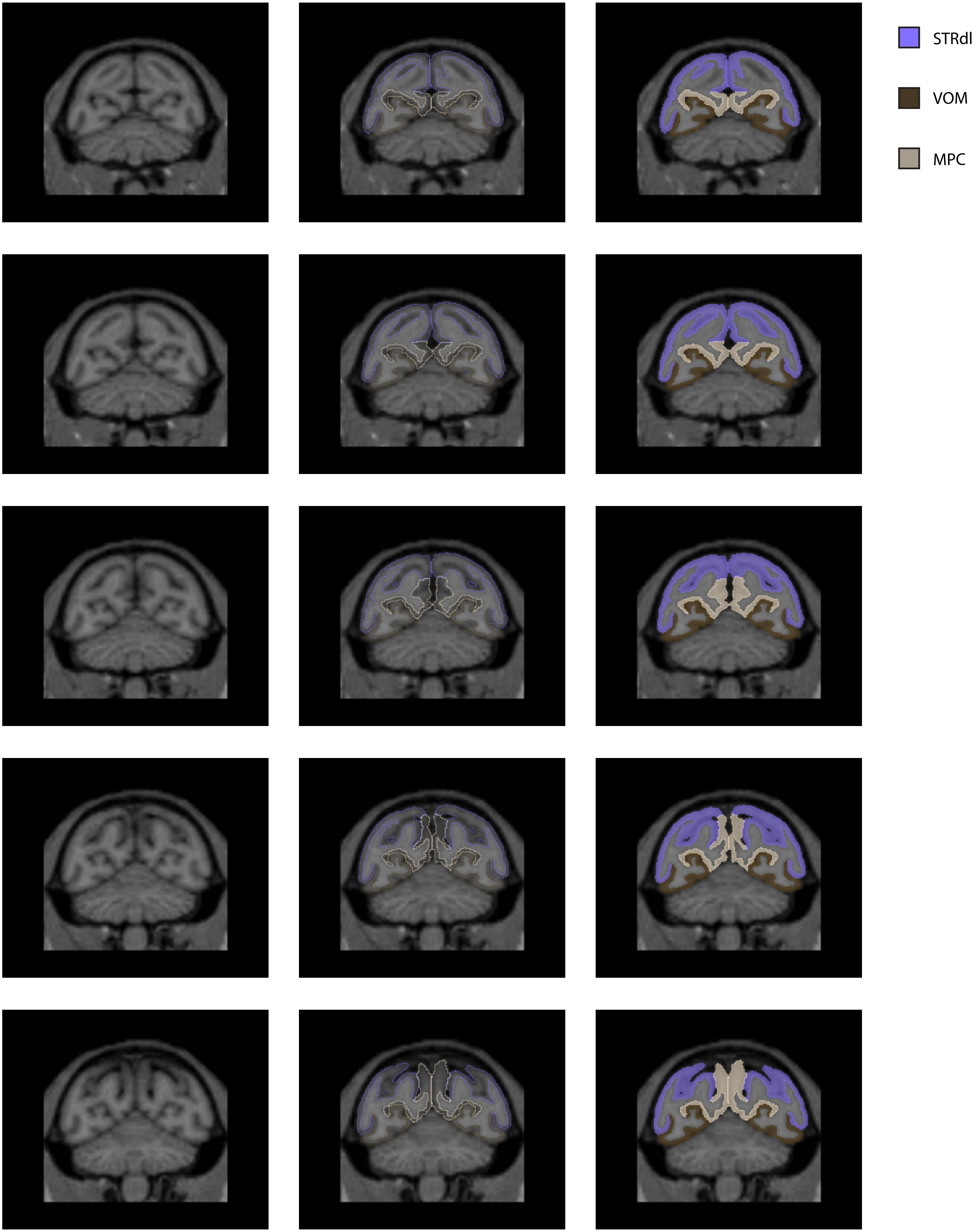
STRdl
VOM
MPC

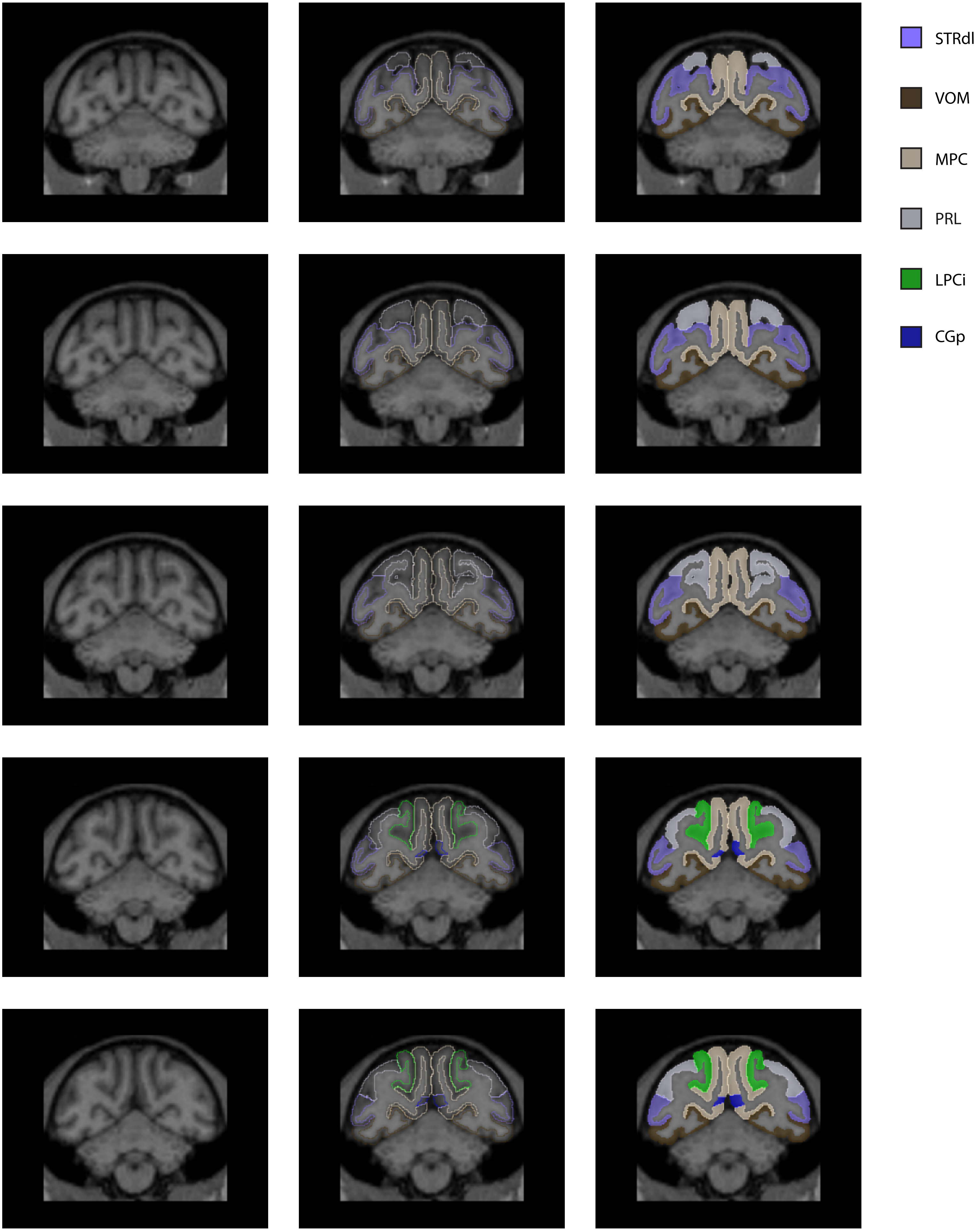

STRdl
VOM
MPC
PRL
LPCi
CGp

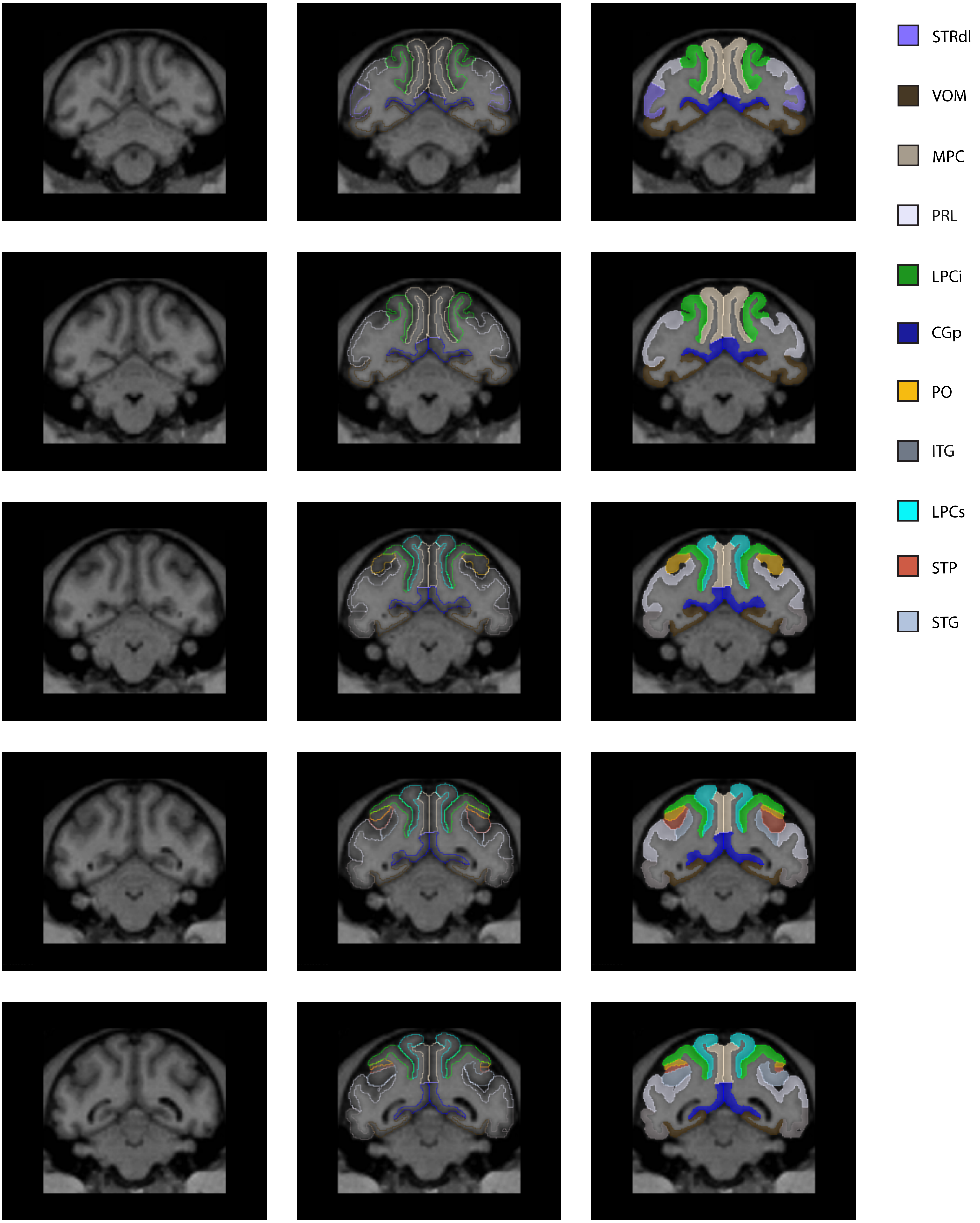
STRdl
VOM
MPC
PRL
LPCi
CGp
PO
ITG
LPCs
STP
STG

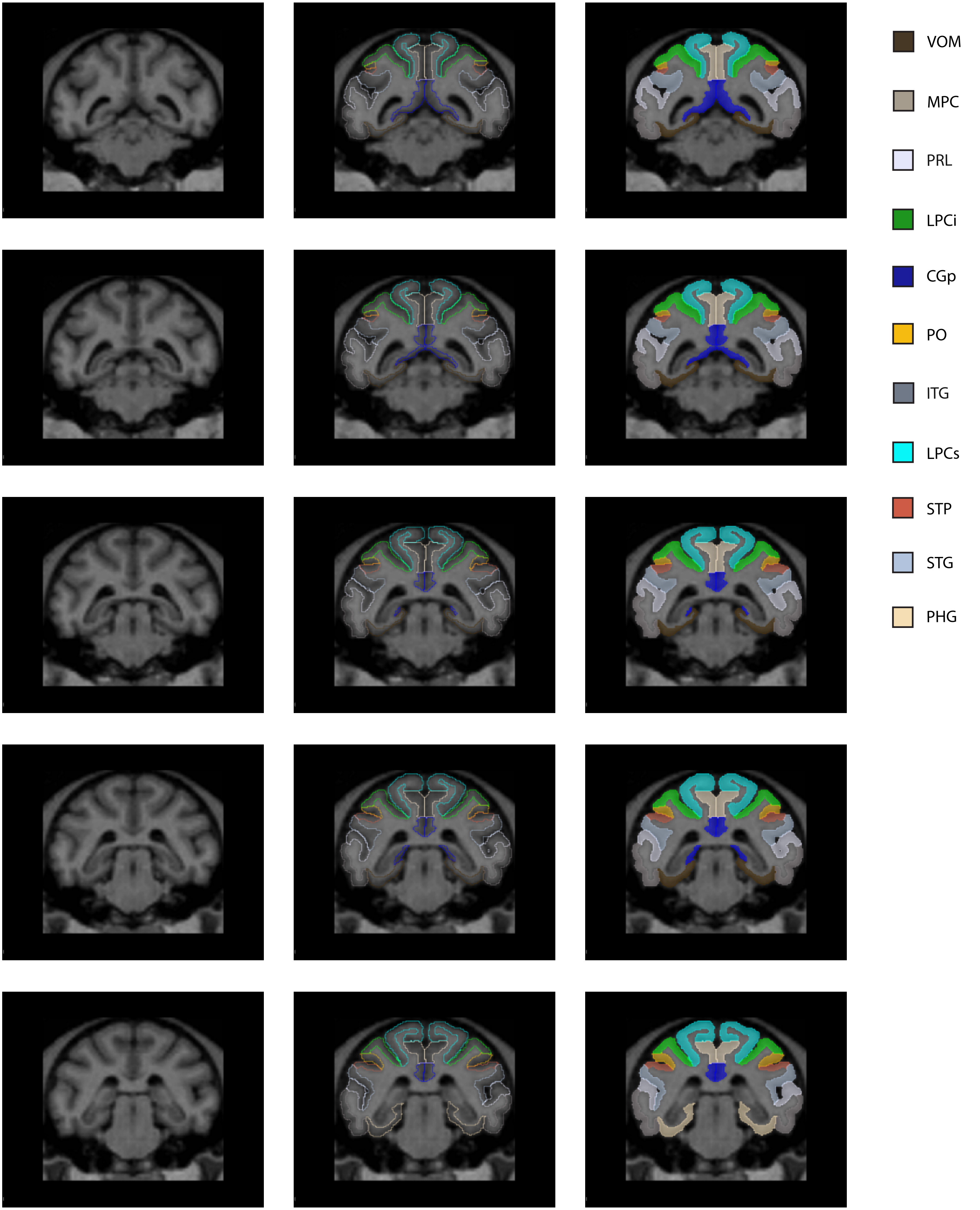
VOM
MPC
PRL
LPCi
CGp
PO
ITG
LPCs
STP
STG
PHG

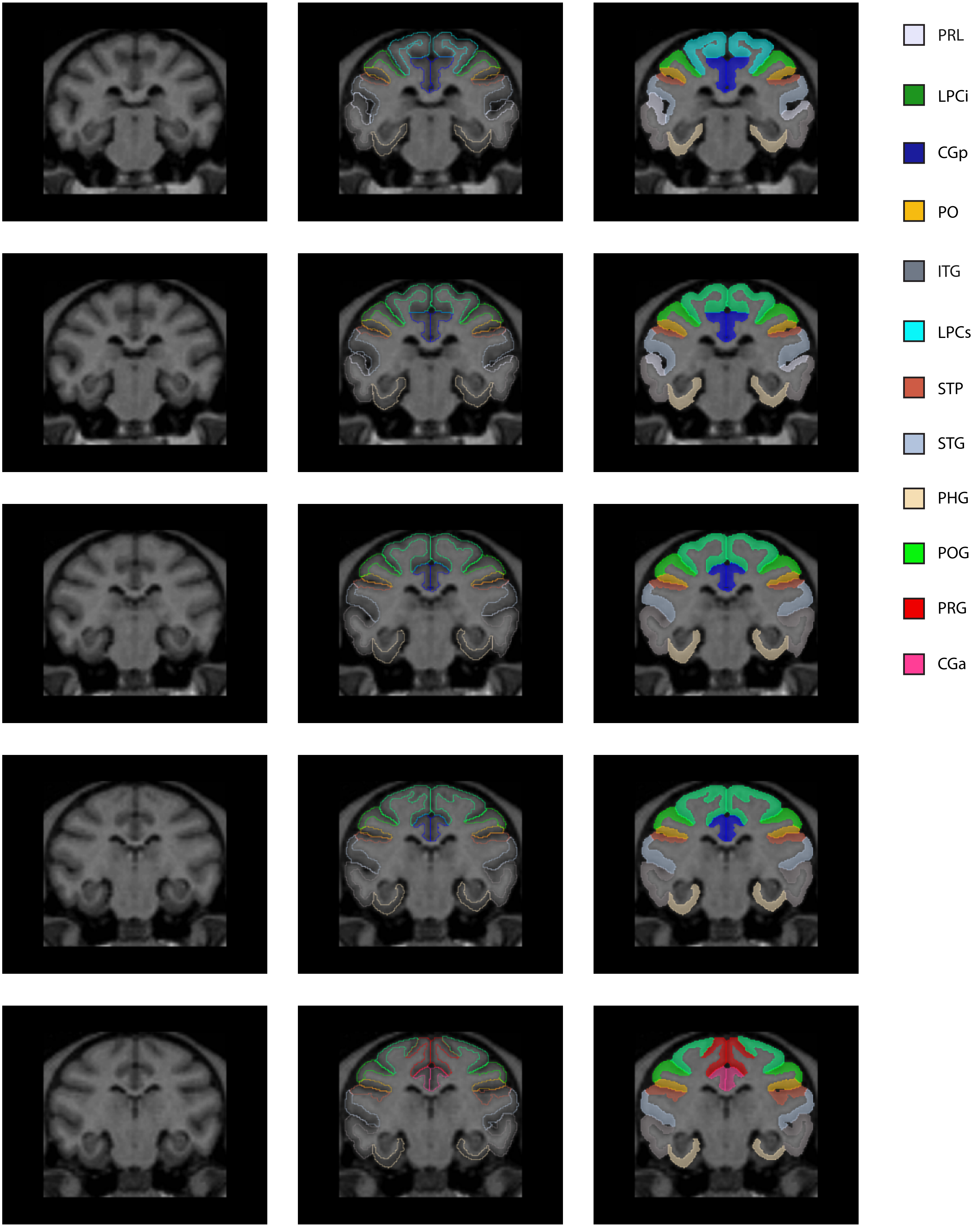
PRL
LPCi
CGp
PO
ITG
LPCs
STP
STG
PHG
POG
PRG
CGa

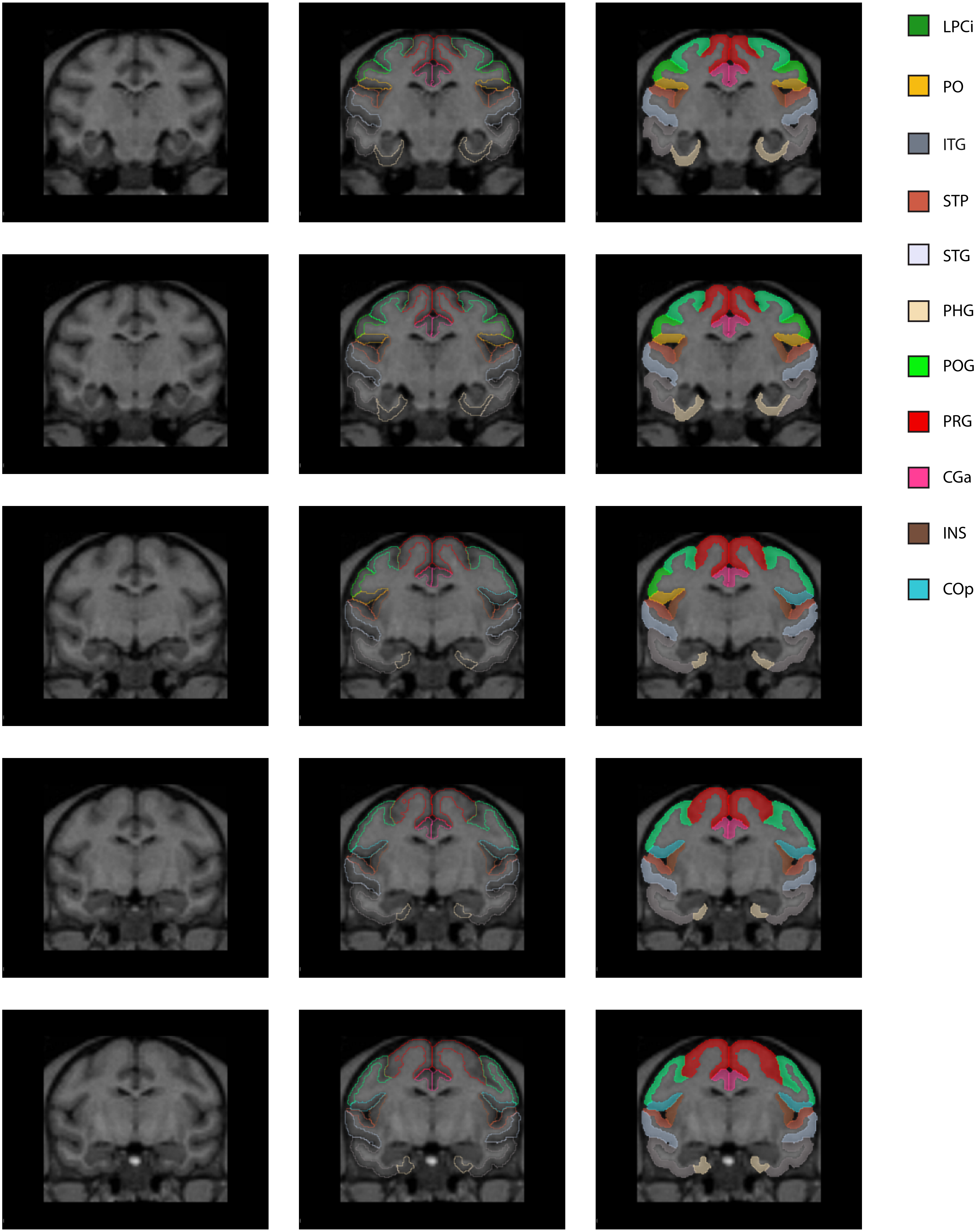
LPCi
PO
ITG
STP
STG
PHG
POG
PRG
CGa
INS
COp

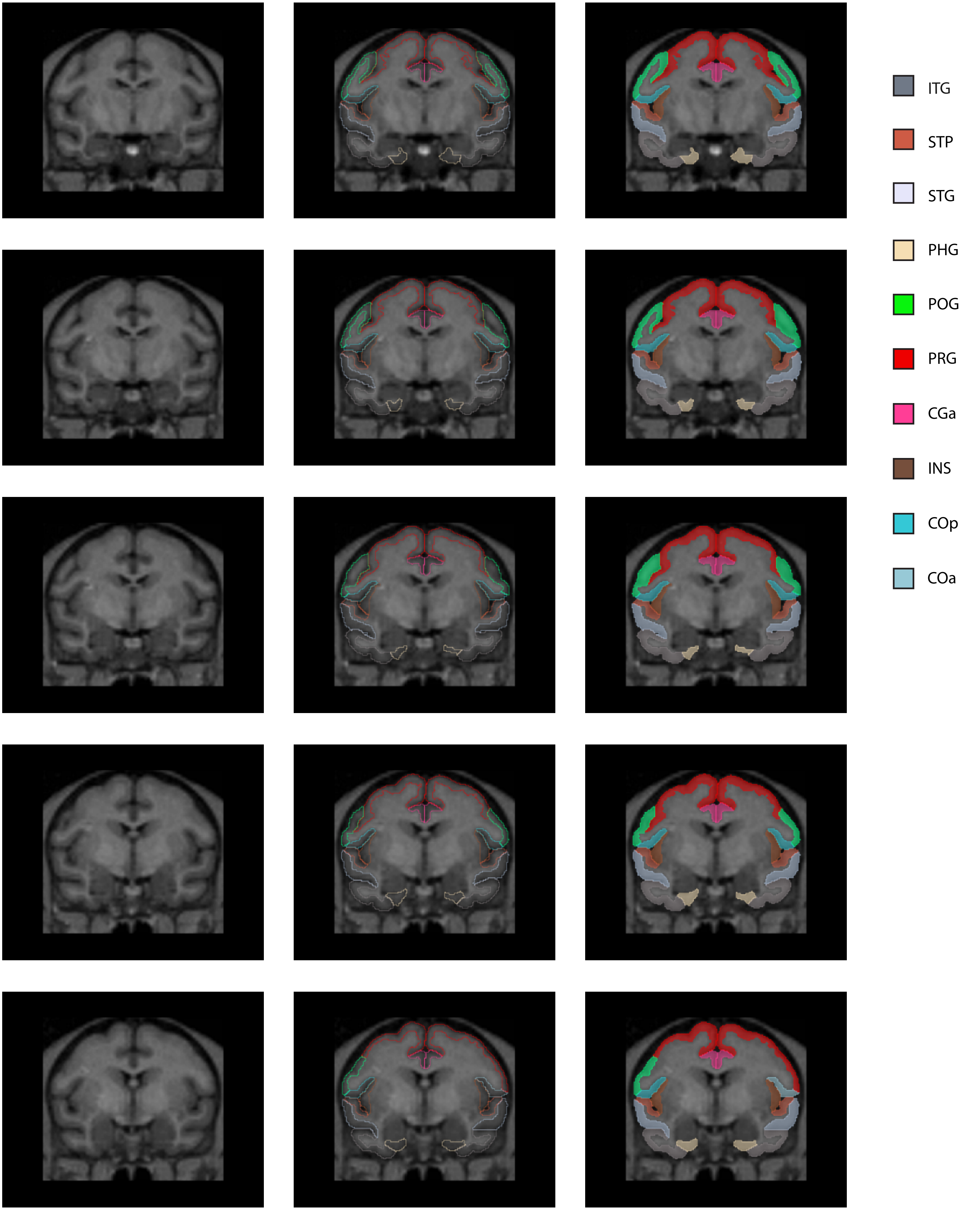
ITG
STP
STG
PHG
POG
PRG
CGa
INS
COp
COa

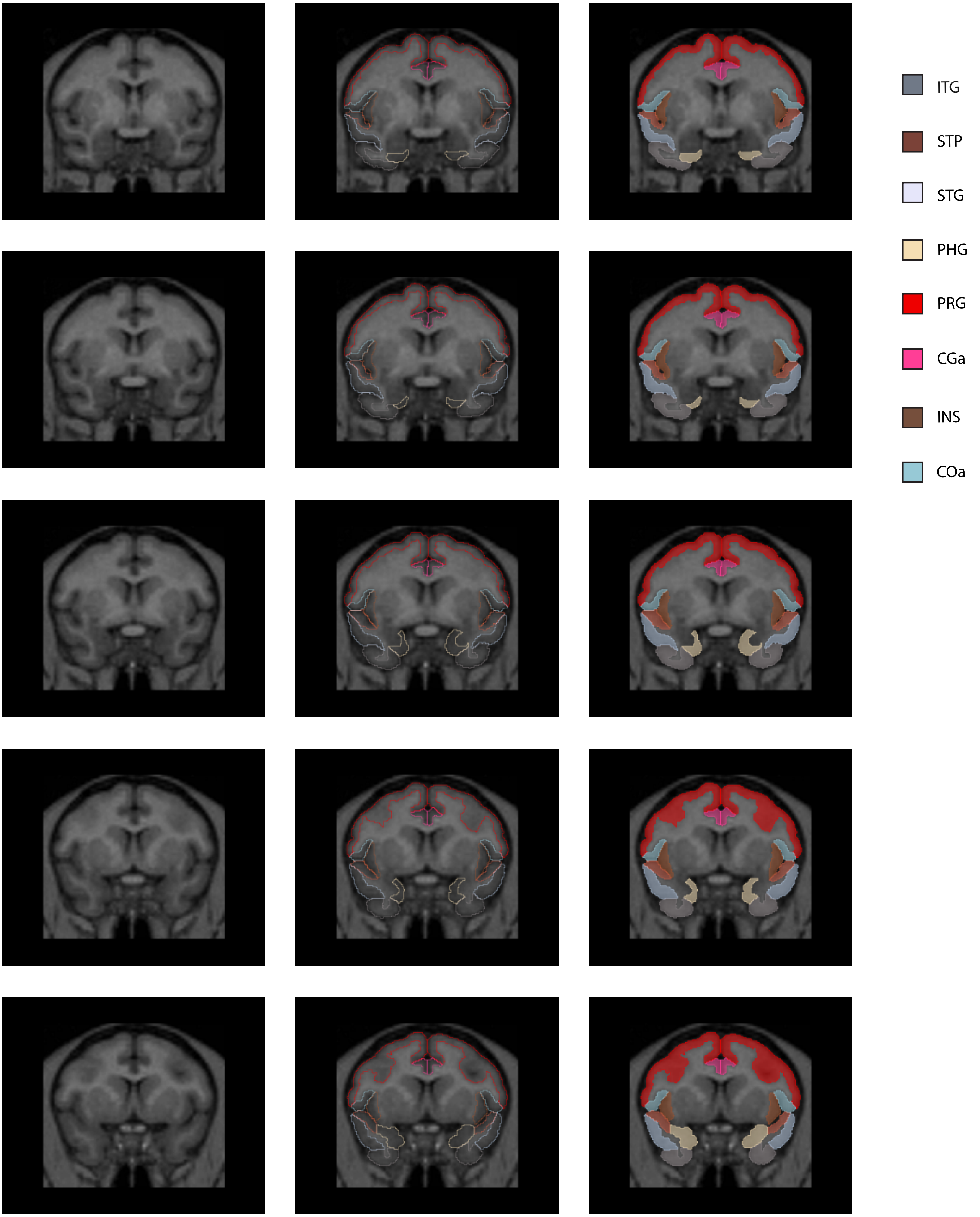
ITG
STP
STG
PHG
PRG
CGa
INS
COa

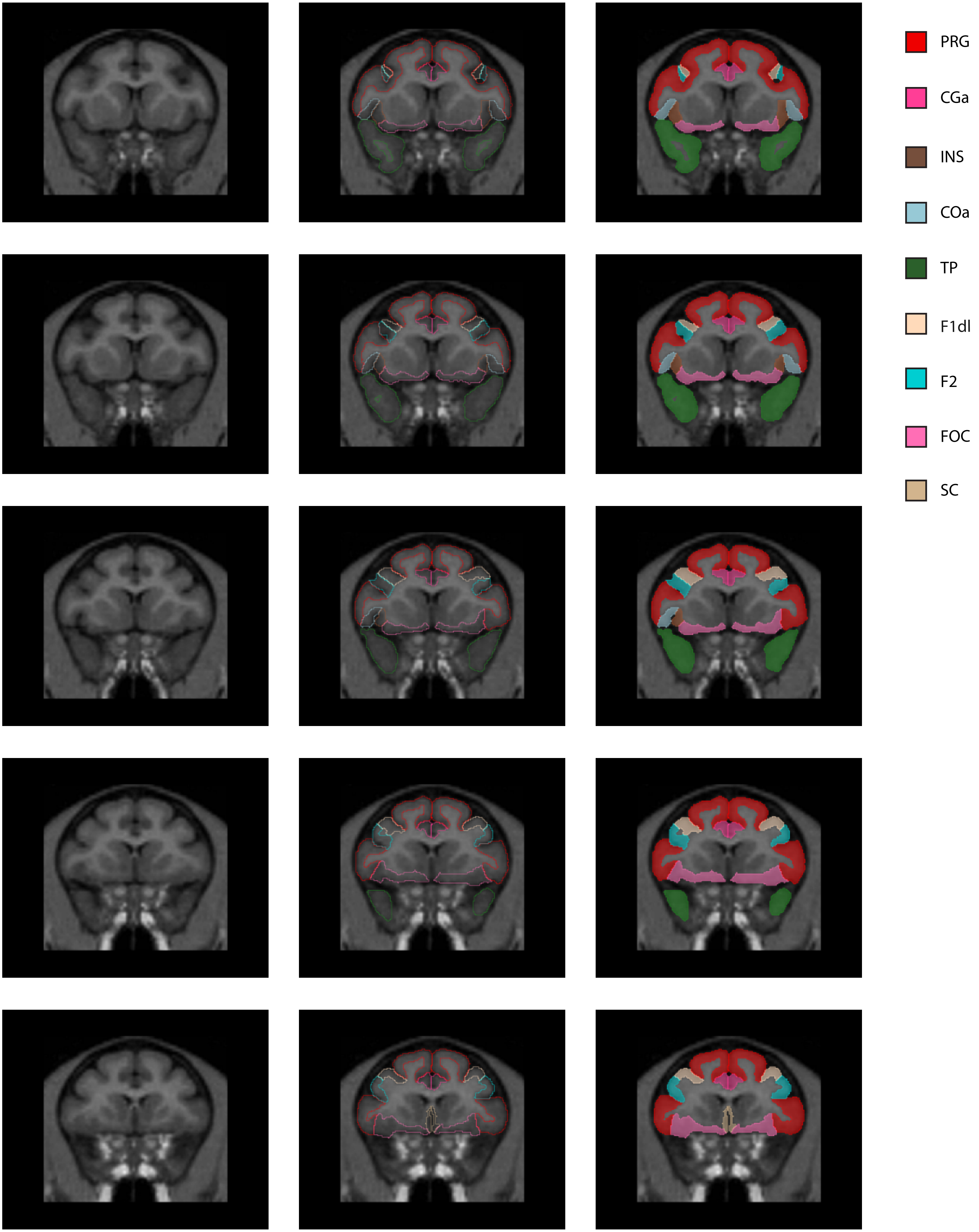
PRG
CGa
INS
COa
TP
F1dl
F2
FOC
SC

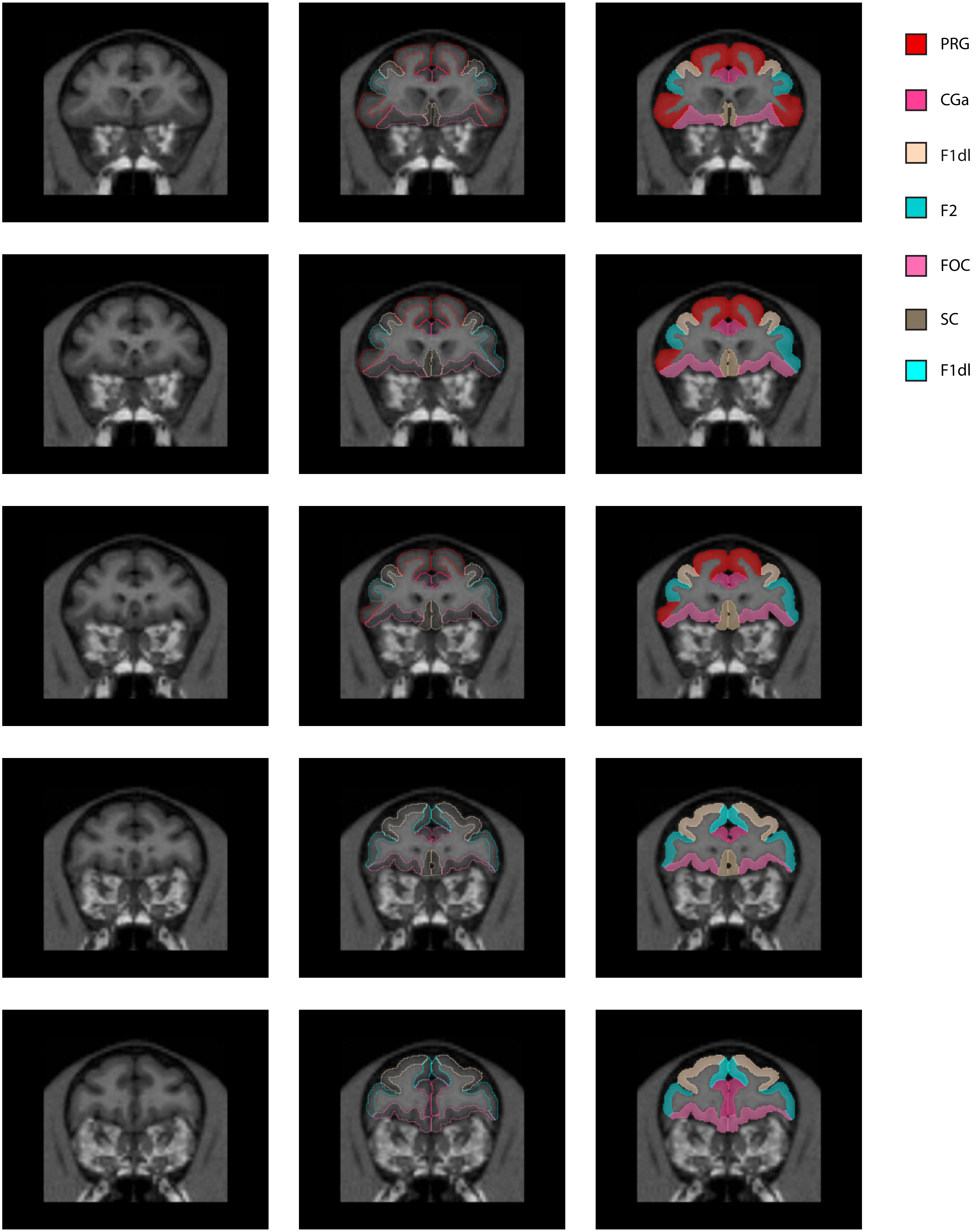

PRG
CGa
F1dl
F2
FOC
SC
F1dl

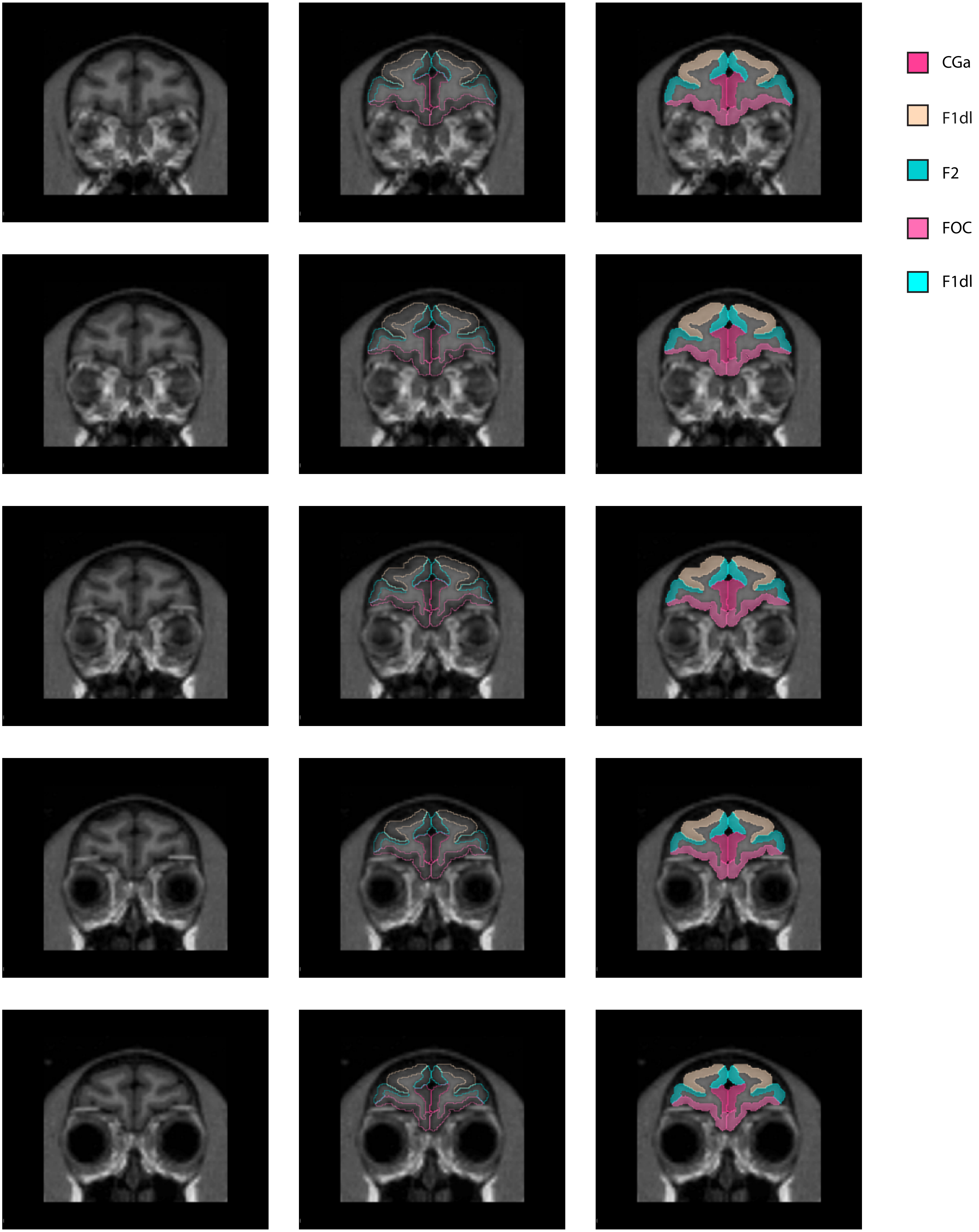
CGa
F1dl
F2
FOC
F1dl

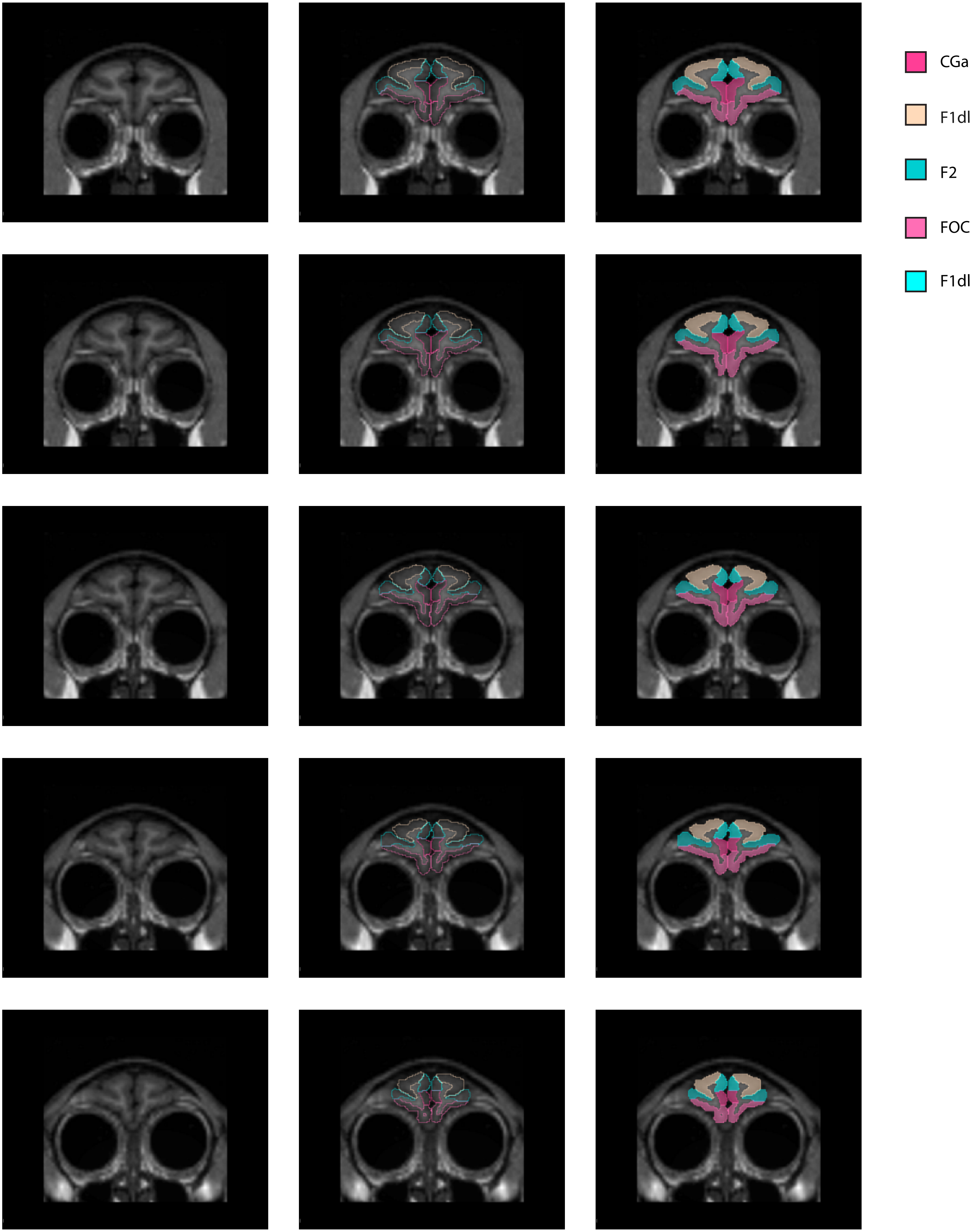
CGa
F1dl
F2
FOC
F1dl

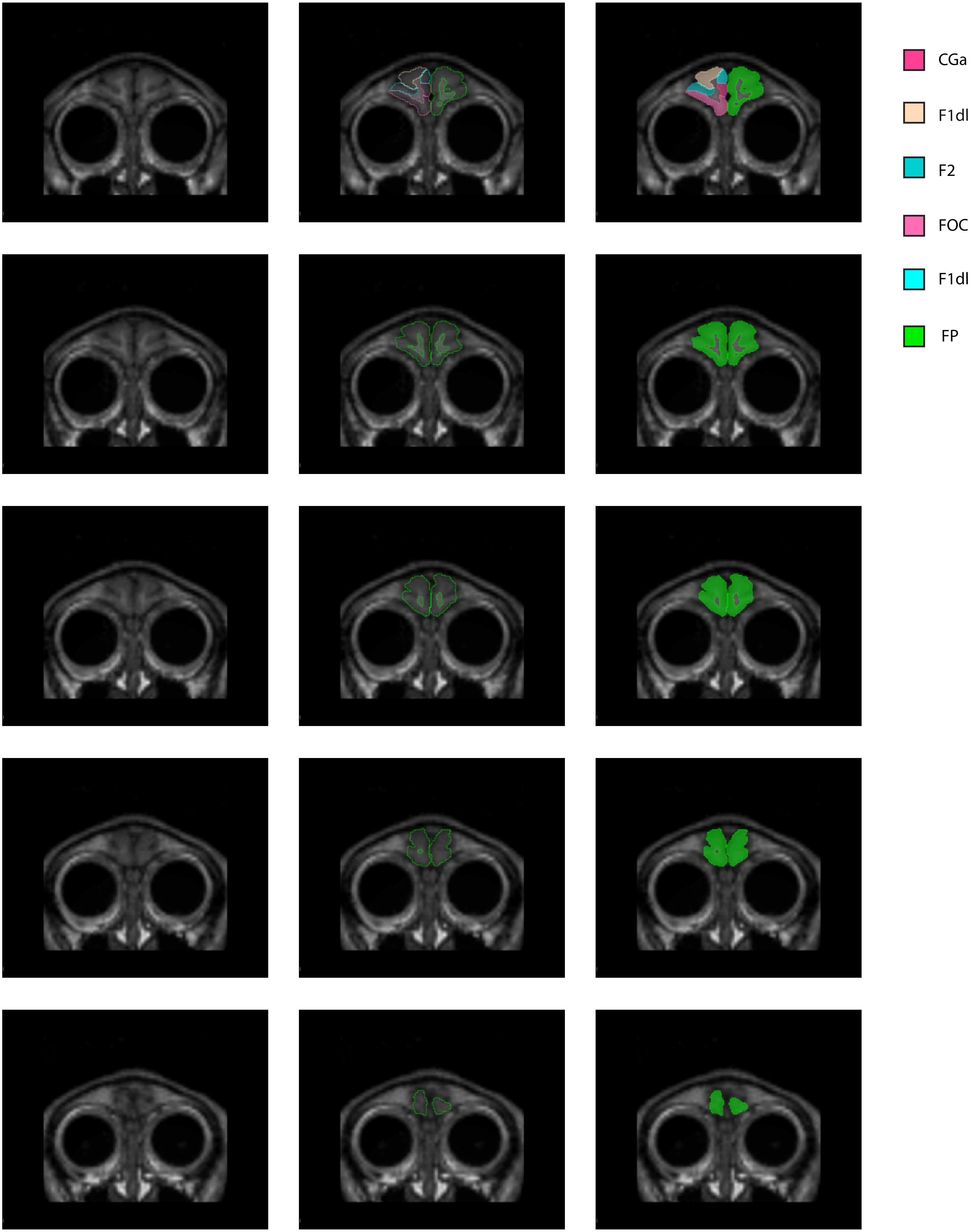
CGa
F1dl
F2
FOC
F1dl
FP
